# Occurrence of *Nigrospora spp.* as the predominant causal agents of leaf spot disease in Cavendish banana in banana plantations in Mindanao Island, Philippines

**DOI:** 10.1101/2025.11.07.685480

**Authors:** Shunsuke Nozawa, Yui Harada, Yoshiki Takata, Keiko Uchida, Mike Andre Malonzo, Reynaldo Valle, Sherman M. Chavez, Aniway F. Penalosa, Kyoko Watanabe

**Affiliations:** School of Agriculture, Tamagawa University, Machida, Tokyo, Japan; BaCaDM project of College of Agriculture, Tamagawa University, 6-1-1 Tamagawa-gakuen, Machida, Tokyo 194-8610, Japan; Unifrutti Tropical Philippines, Inc., Km. 15, Panacan, Davao City 8000, Philippines

## Abstract

Banana leaf diseases are a significant threat to Cavendish banana production. In the Philippines, the main disease has been diagnosed as Black sigatoka disease caused by *Pseudocercospora fijiensis* based on symptoms. However, our study showed that the main pathogen in Mindanao island, the largest banana-producing region in the Philippines, belongs to the genus *Nigrospora*, contradicting previous assumptions. We clarified the phylogenetic positions of 160 *Nigrospora* isolates based on molecular phylogenetic analyses using ITS, *β-tubulin*, and *tef1α* sequences, and compared their morphology with known species. Molecular phylogenetic and morphological analysis revealed that *Nigrospora* isolates comprised *N. chinensis*, *N. lacticolonia*, *N.* cf. *singularis*, *N. sphaerica*, *N. vesicularifera*, and a novel species, *N. nigrocolonia*. Pathogenicity tests on banana leaves confirmed that these species are pathogenic. Species other than *N. sphaerica* were for the first time reported as pathogens of banana leaf. The results of the fungicide sensitivity test using 14 fungicides, including pyrimethanil, spiroxamine, and tebuconazole, for the Sigatoka disease showed 100% inhibition of all isolates at 100 ppm of active ingredients. However, low-sensitivity isolates were observed for the remaining 11 fungicides. Our findings indicated the need for a comprehensive review of banana leaf disease prevention strategies.

## Introduction

Banana is a cash crop in Southeast Asia, Africa, and Latin America, with an annual production of 1.51 million tons in 2023, making it the most produced fruit globally (FAOSTAT 2023). The Cavendish banana is the most widely traded and globally distributed variety among all banana cultivars. Cavendish is exported in significant quantities among the banana cultivars traded internationally. More than 20 million tons of Cavendish banana are exported. The leading exporters of Cavendish banana include Colombia, Costa Rica, Ecuador, Guatemala, and the Philippines, and its major importers are the European Union and United States.

Banana production is affected by diseases such as Fusarium wilt caused by *Fusarium* spp. [1,2], leaf blight by *F. saccari* and *Alternaria* jacinthicola [3,4]; leaf spots by *Colletotrichum* spp., *Pseudocercospora eumusae*, and *Nigrospora* shaerica [5–7]; and Sigatoka disease by *Pseudocercospora* spp. [8]. Among these diseases, Sigatoka disease is one of the most serious, causing an estimated 3% annual reduction in banana yield and economic losses of approximately 1.6 billion dollars annually [9]. The Sigatoka disease is caused by several *Pseudocercospora* spp.; black Sigatoka by *P. fijiensis*, and yellow Sigatoka by *P.* musae [10]. Among these diseases, *P. fijiensis* (black Sigatoka) is the most virulent and causes serious damage. Since its discovery in Fiji in 1964 [11], the disease has spread globally owing to its virulence and infectiousness, becoming one of the most serious threats to banana production.

The Philippines is the world’s fifth-largest producer of bananas, with an annual output of approximately 5.9 million tons in 2023 (FAOSTAT 2023). Bananas are pivotal cash crops for export, with an estimated annual export volume of approximately 2 million tons. However, the country is grappling with a significant challenge: black Sigatoka disease, the management of which has necessitated substantial financial investments [10]. To address this issue, a leaf removal strategy was implemented to eliminate leaves exhibiting disease symptoms and maintain a minimum of five leaves per plant. This approach involves fungicide applications, sprayed approximately 50 to 70 times per year, to prevent disease transmission. According to the Banana Working Group of the Fungicide Resistance Action Committee (FRAC), the following fungicide groups are recommended for the control of Sigatoka disease: amines, anilinopyrimidines (APs), benzimidazoles (BCMs), demethylase inhibitors (DMIs), guanidines, multi-site inhibitors, N-Phenylcarbamates, Qi Inhibitors (QiIs), Qo Inhibitors (QoIs), Succinate Dehydrogenase Inhibitors (SDHIs), and Toluamides. Fungicide application incurs high annual costs and is the most expensive activity in banana production. However, there is a paucity of surveys conducted in the Philippines to investigate the emergence of new pathogens despite the presence of multiple diseases that can present spot symptoms similar to Sigatoka disease in the world [5,7,12].

Our study shows that *Nigrospora* spp., rather than *Pseudocercospora* spp., is the dominant pathogen on the island of Mindanao in the Philippines. This symptom closely resembles that of Sigatoka disease, as it causes the formation of brown to black spots with a yellow halo and the lesions expand along the leaf veins (Fig. 1). Consequently, locally, any disease exhibiting such symptoms is diagnosed as Sigatoka disease.

**Fig. 1.**
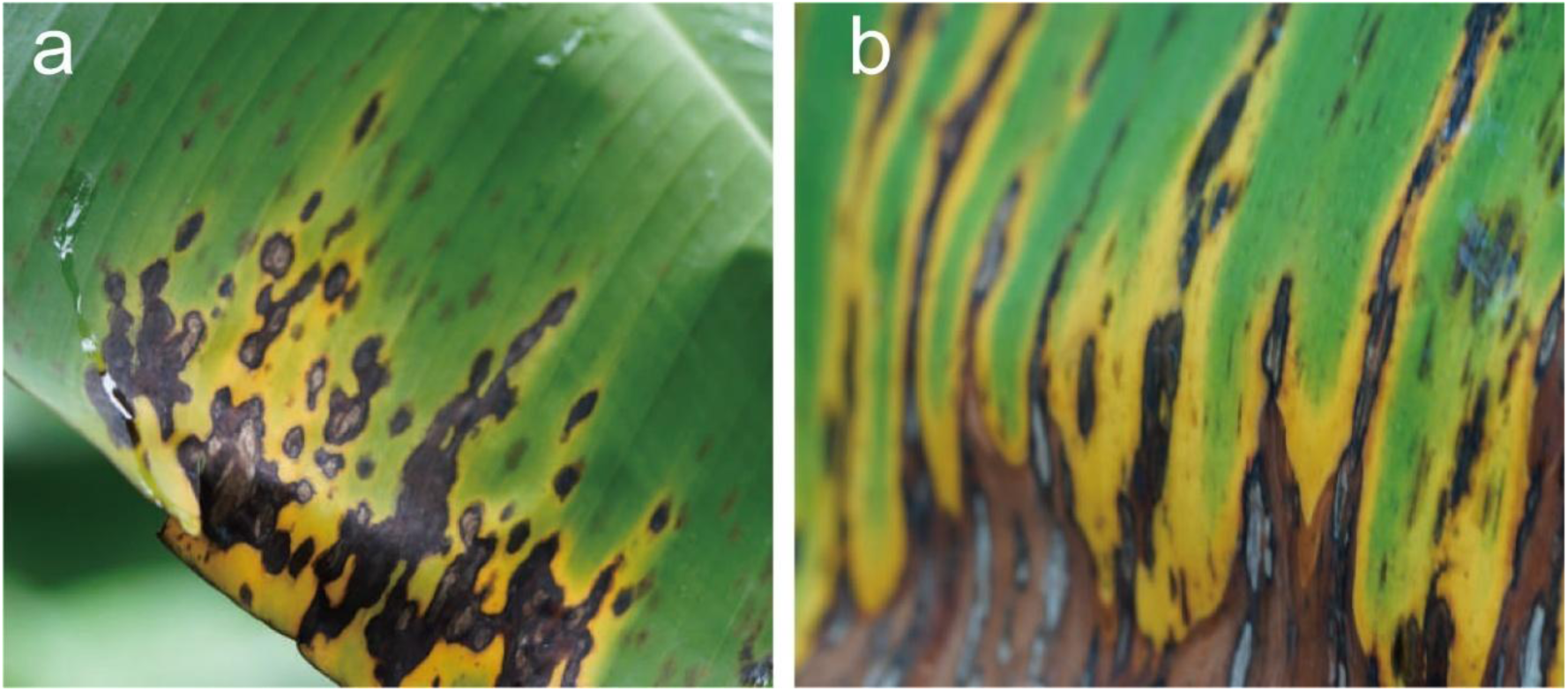
Symptoms of leaf spot on banana leaves in fields. Naturally infected leaves in banana plantations (a, b).

To provide basic information for control measures against this disease, which is mistakenly diagnosed as Sigatoka disease, we identified the pathogens belonging to the genus *Nigrospora* at the species level and conducted haplotype analysis. In addition, in vitro sensitivity tests were conducted to identify effective fungicides currently used to control black Sigatoka disease and hypothesize the potential emergence of resistant *Nigrospora* spp. isolates.

## Results

### Frequency of the genus *Nigrospora*

We obtained 217 isolates from diseased banana leaves. As a preliminary identification to determine the genus, a BLAST search of the ITS region in the NCBI database was conducted. A Basic Local Alignment (BLAST) search showed that our isolates consisted of 18 genera (Table 1). Out of 217 isolates, 160 belonged to the genus *Nigrospora*, which was the most frequently detected fungal genus. In addition, the genus *Nigrospora* was dominant at all the collection locations. The proportions of *Nigrospora* were 96.6%, 70.2 %, 53.5 %, 92.9 %, 69 %, and 75% in Dabong-dabong, Little Panay, Panambon, Patag-Alanib, Tugbok, and Ula, respectively. We did not perform further analyses to identify the species of isolates that did not belong to the genus *Nigrospora*.

**Table 1.** Preliminary identification of 217 fungal isolates from banana leaf samples from six cities of Mindanao Island.

| Genus<sup>a</sup> | Isolate numbers from different cities of Mindanao Island |  |  |  |  |  |
| --- | --- | --- | --- | --- | --- | --- |
|  | Dabong-dabong | Little Panay | Panambon | Patag-Alanib | Tugbok | Ula |
| *Alternaria* | 0 | 0 | 0 | 0 | 1 | 0 |
| *Annulohypoxylon* | 0 | 0 | 0 | 1 | 0 | 0 |
| *Colletotrichum* | 0 | 0 | 0 | 0 | 2 | 3 |
| *Curvularia* | 0 | 0 | 1 | 0 | 1 | 0 |
| *Dimorphiseta* | 0 | 0 | 1 | 0 | 0 | 0 |
| *Exserohilum* | 0 | 1 | 0 | 0 | 0 | 0 |
| *Fusarium* | 0 | 3 | 6 | 0 | 0 | 4 |
| *Lasiodiplodia* | 1 | 0 | 0 | 0 | 1 | 0 |
| *Microdochium* | 0 | 0 | 3 | 0 | 0 | 0 |
| *Musicillium* | 0 | 0 | 0 | 0 | 0 | 1 |
| *Mycosphaerella* | 0 | 1 | 0 | 0 | 0 | 0 |
| *Neosetophoma* | 0 | 2 | 3 | 0 | 0 | 0 |
| *Nigrospora* | 28 | 33 | 21 | 25 | 20 | 33 |
| *Ochrocladosporium* | 0 | 1 | 0 | 0 | 0 | 0 |
| *Phaeosphaeria* | 0 | 5 | 6 | 0 | 3 | 2 |
| *Preussia* | 0 | 0 | 0 | 1 | 0 | 0 |
| *Pseudopestalotiopsis* | 0 | 1 | 0 | 0 | 1 | 0 |
| *Zasmidium* | 0 | 0 | 0 | 0 | 0 | 1 |

<sup>a</sup>The fungal isolates were identified based on ITS sequences.

**Table 2.**
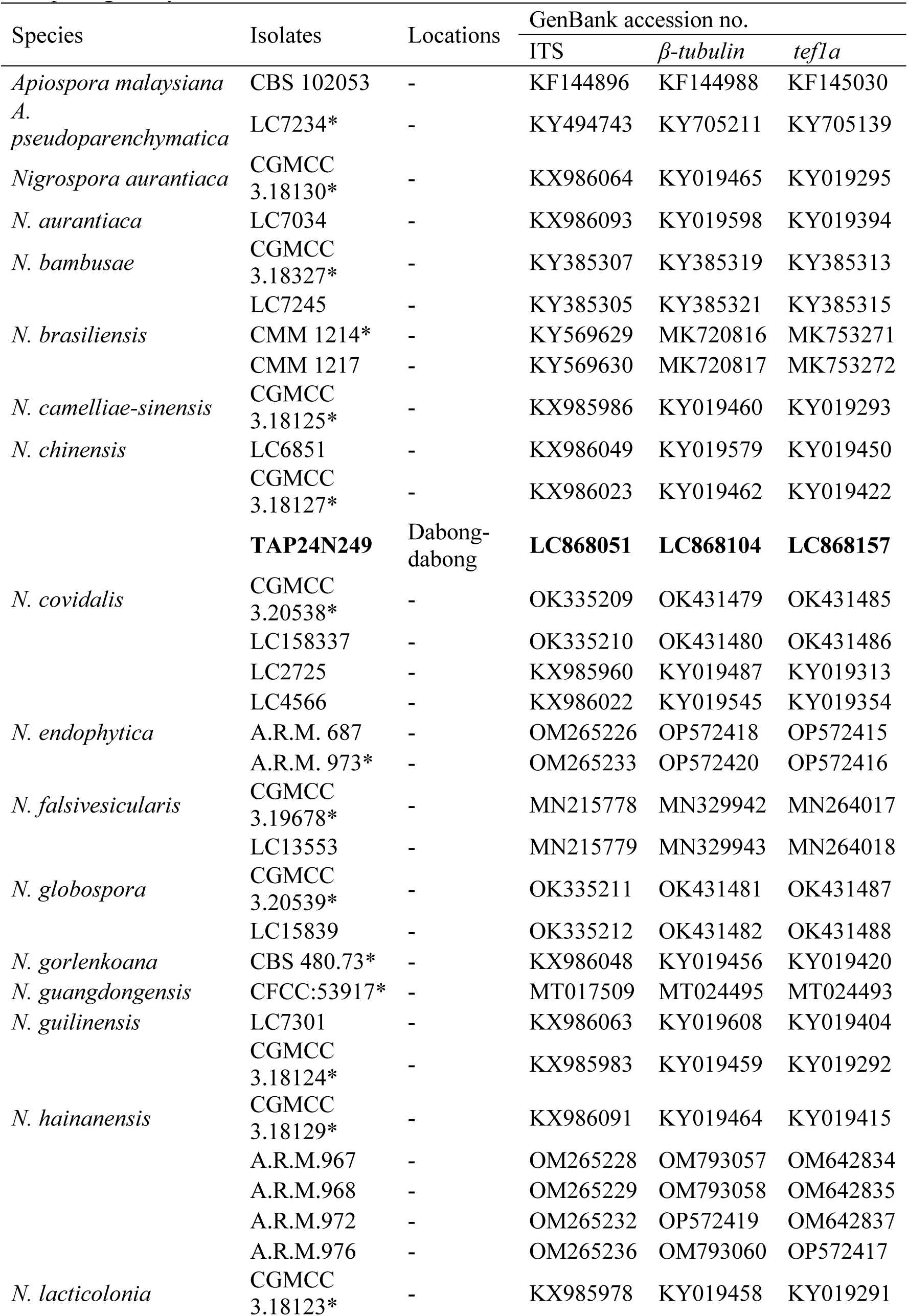
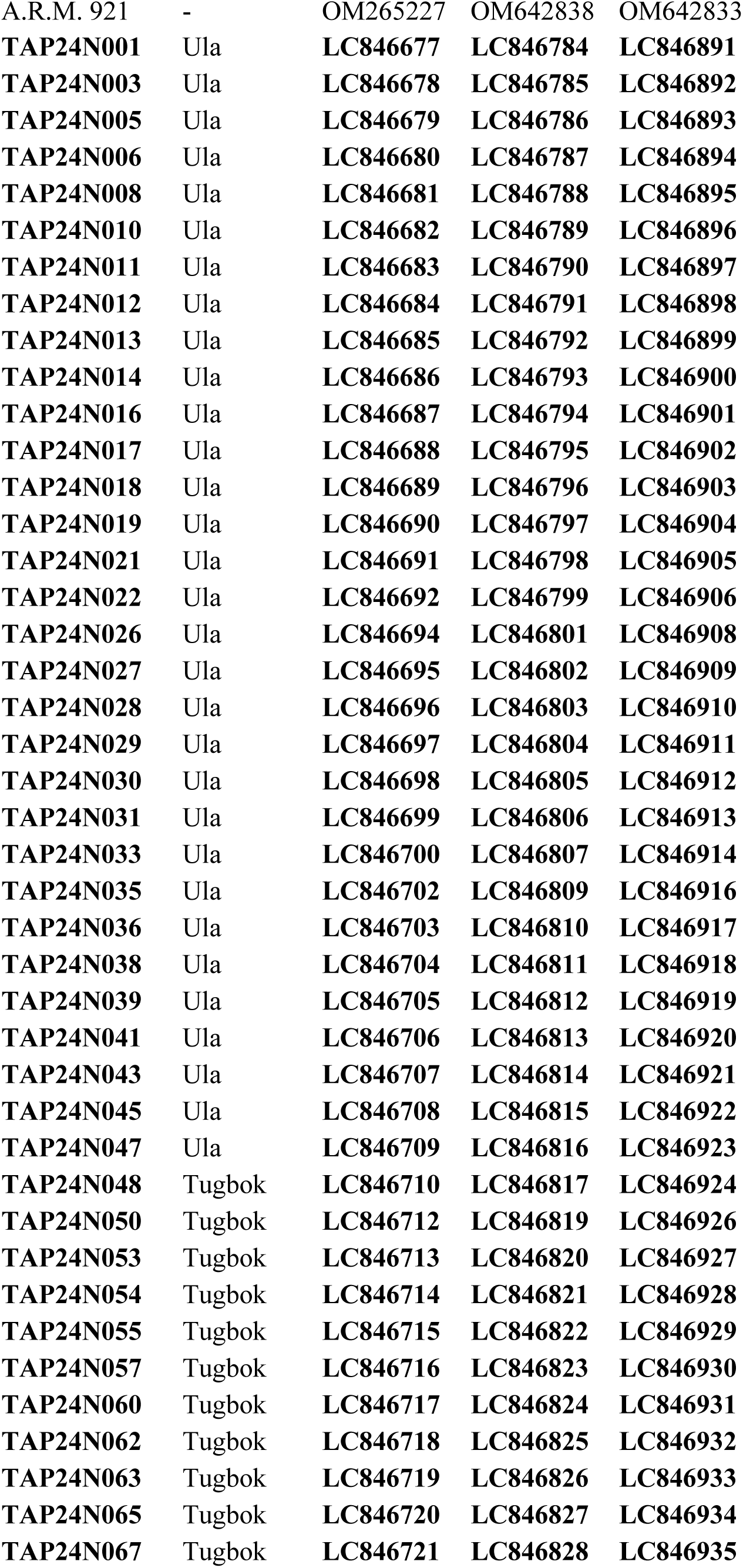
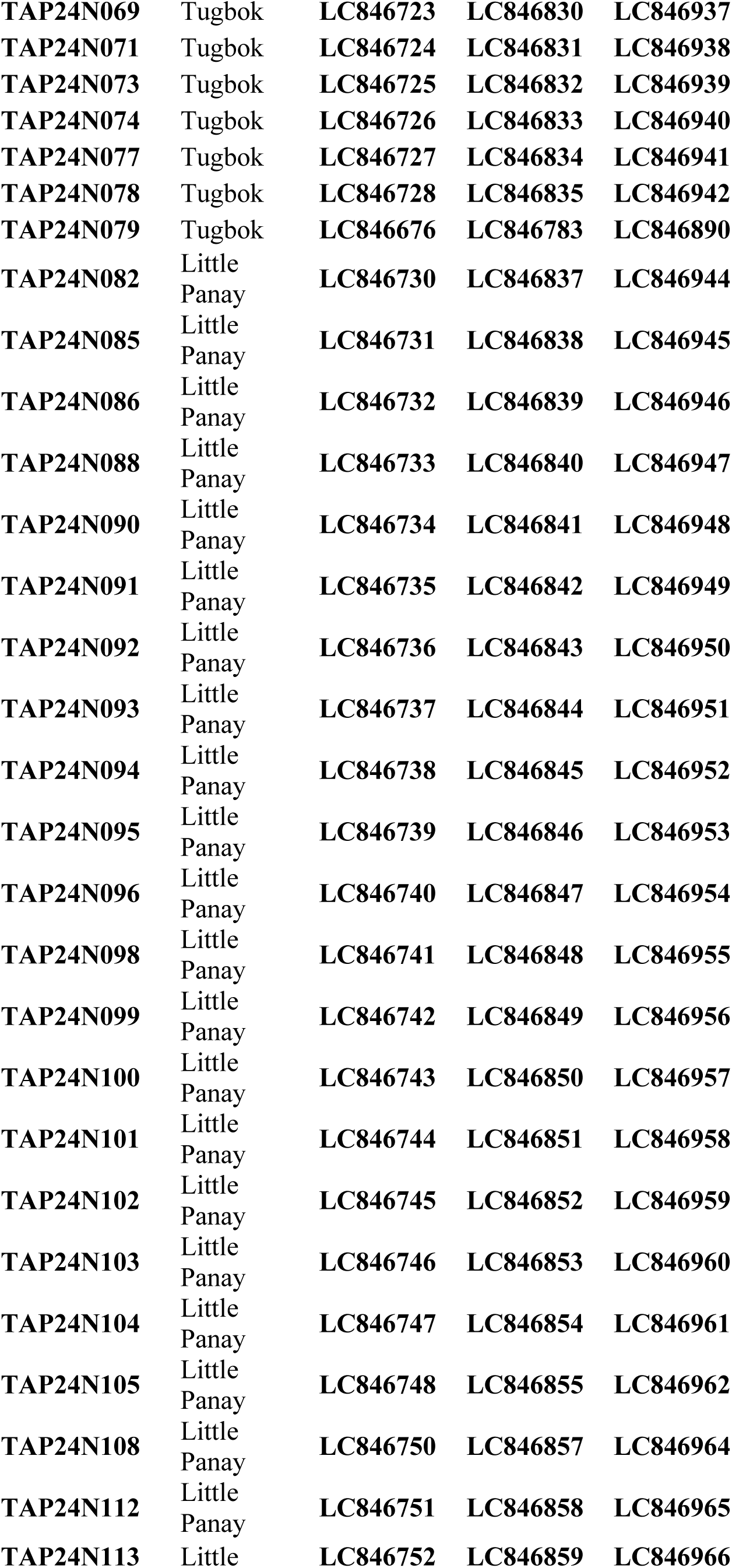
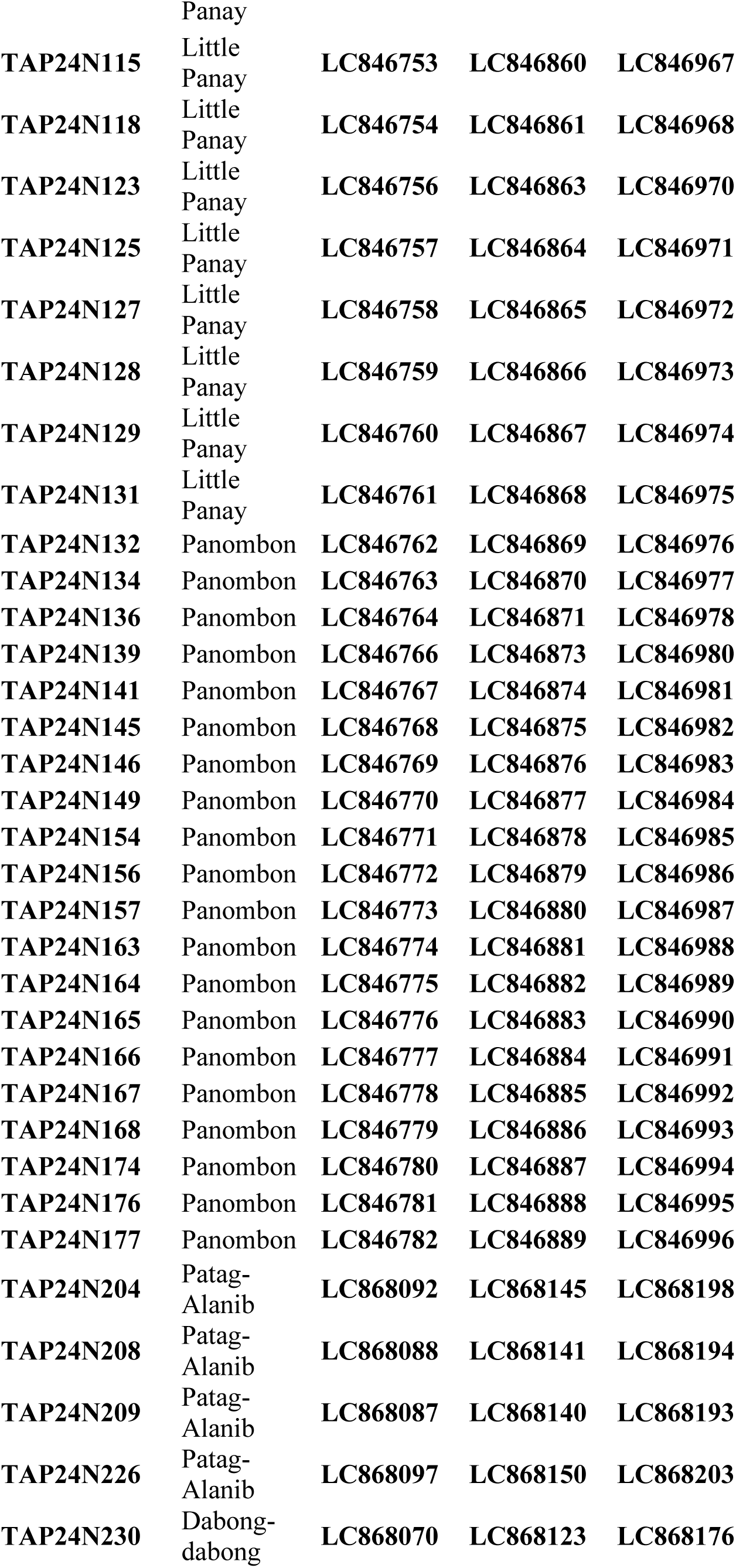
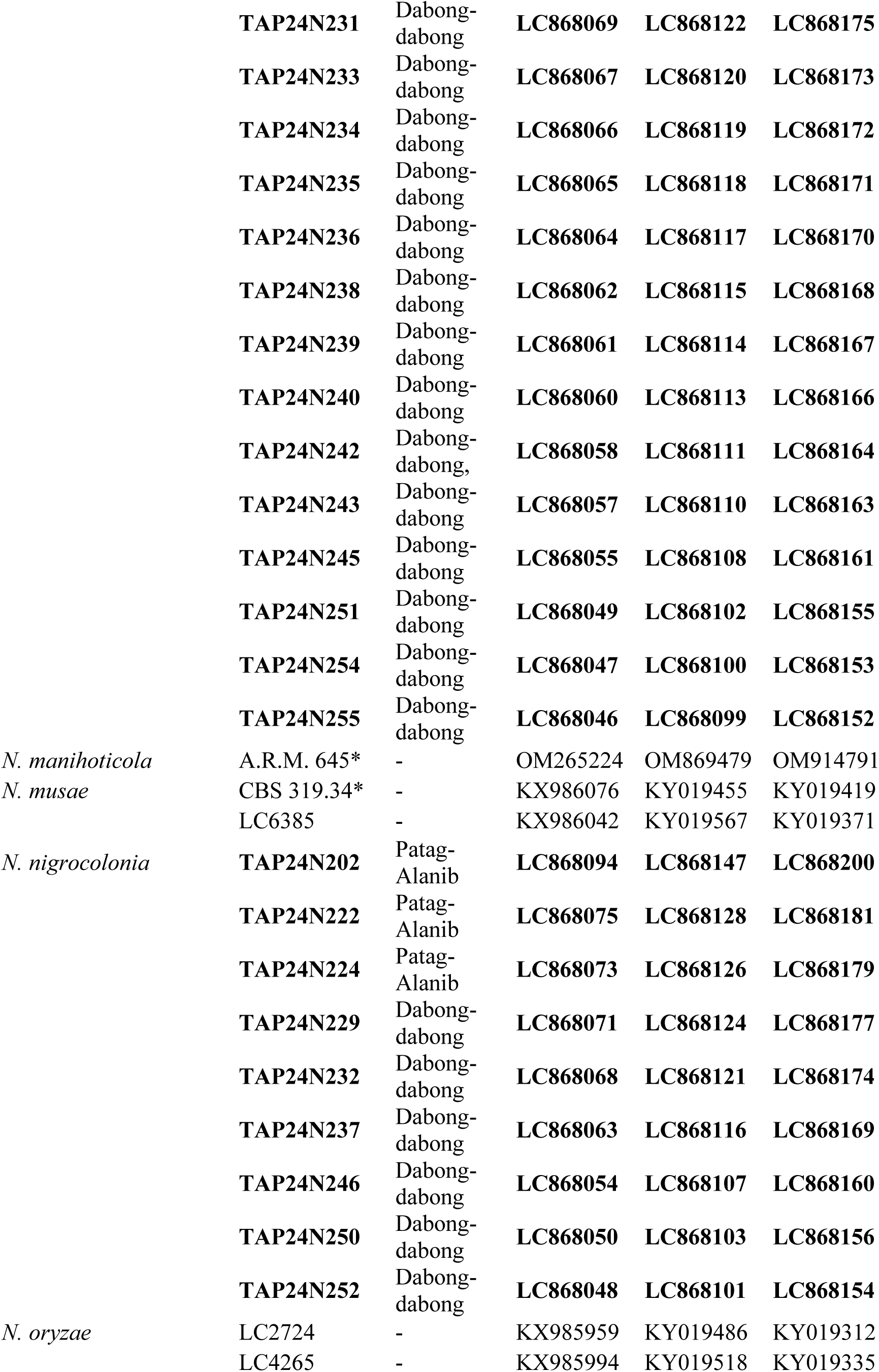
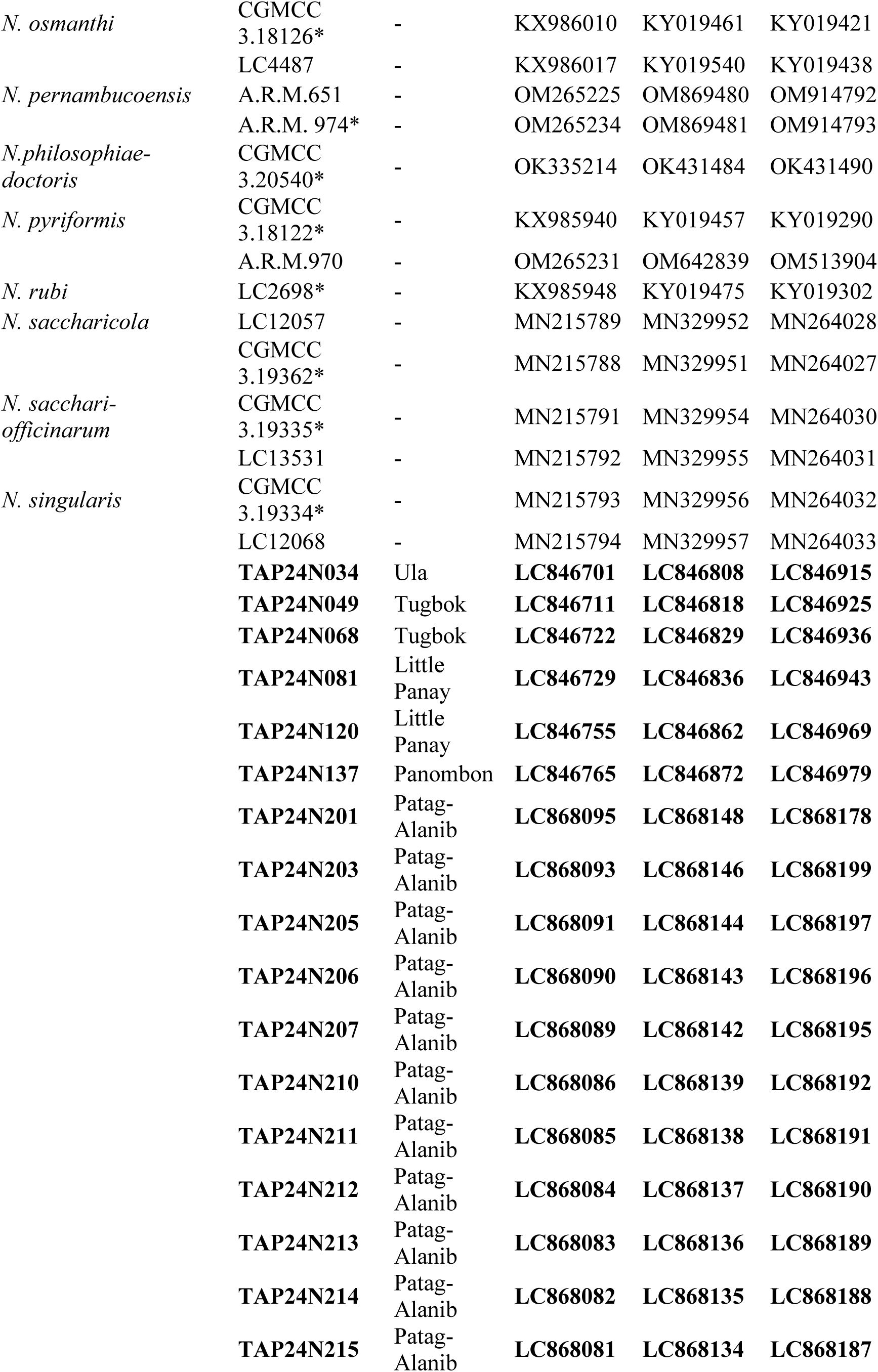
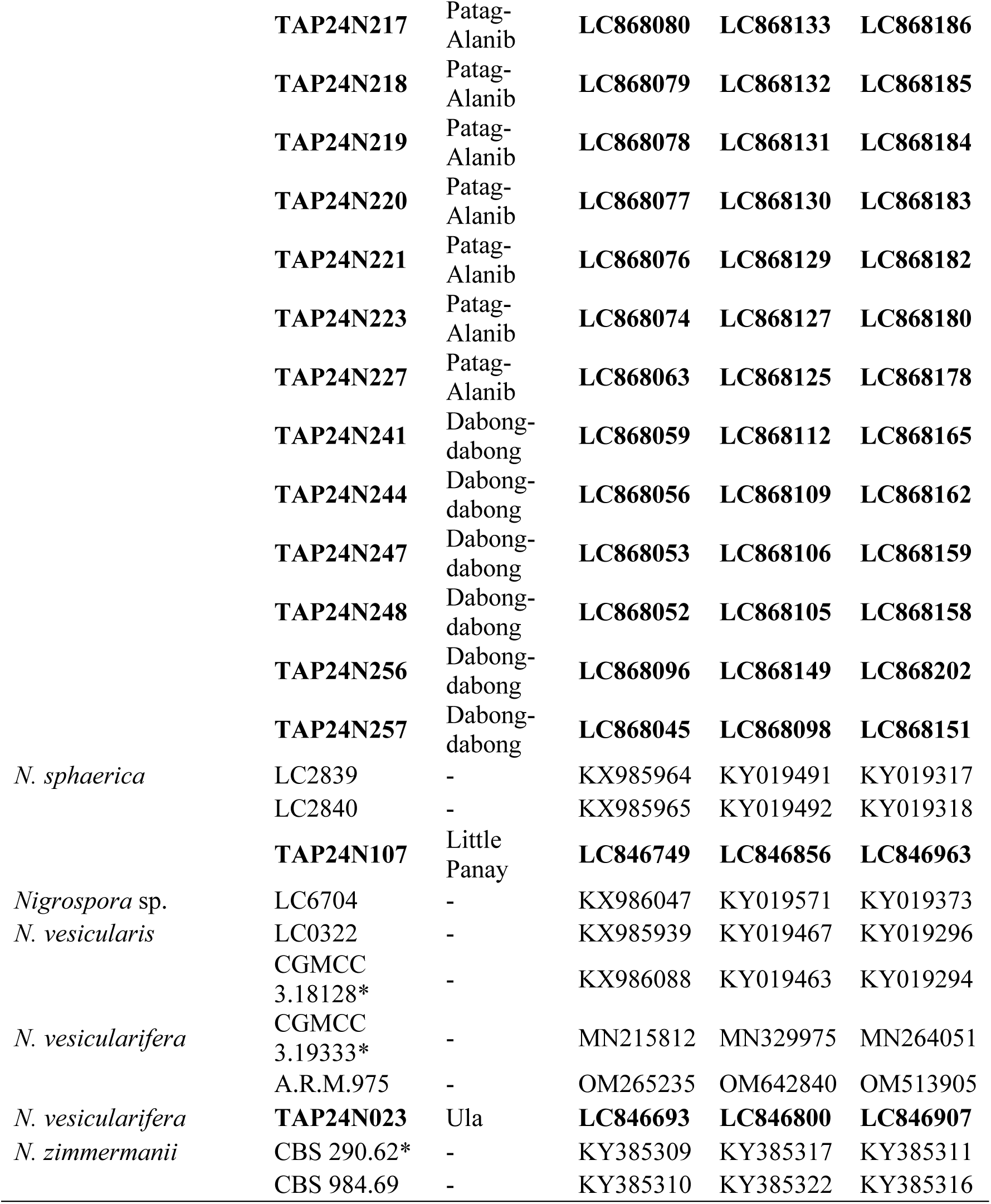
GenBank accession numbers of the DNA sequences used in phylogenetic analysis and pathogenicity of our isolates.

### Phylogeny of isolates belonging to genus *Nigrospora*

To reveal the phylogenetic position of our isolates, we constructed molecular phylogenetic trees based on the combined data set of ITS, *β-tubulin*, and *tef1a* nucleotide sequences. The alignment length, number of variable sites, and number of parsimony-informative sites in the DNA data were 1,156, 545, and 461 bp, respectively (see Supplementary Data Set). The phylogenetic tree (Fig. 2; see Supplementary Data Set) showed that our isolates belonged to a clade comprised of *N. camelliae-sinensis* M. Wang & L. Cai and *N. singularis* M. Raza & L. Cai (30 isolates: 13.4% of all isolates; BPP/ML = 0.99/92, *N. chinensis* clade (1 isolate: 0.5% of all isolates; BPP/ML = 1.0/99), *N. lacticolonia* (118 isolates: 54.6% of all isolates; BPP/ML = 1.0/99), *N. nigrocolonia* Nozawa & Kyoko Watan. clade (9 isolates: 11.1% of all isolates; BPP = 1.0), *N. sphaerica* clade (1 isolate: 0.5% of all isolates; BPP/ML = 1.0/99), and *N. vesicularifera* clade (1 isolate: 0.5% of all isolates; BPP/ML = 1.0/99). Regarding the isolates belonging to a clade comprising of *N. camelliae-sinensis* and *N. singularis*, the evolutionary distance between the isolates and the ex-type strain of *N. singularis* (CGMCC 3.19334) is closer than that to *N. camelliae-sinensis*. However, identifying the isolates collectively as *N. singularis* is not possible because *N. camelliae-sinensis* belongs within the same clade. Therefore, in this study, these isolates were treated as strains closely related to *N. singularis* and designated as *N.* cf. *singularis*. Although the clade comprising *N. camelliae-sinensis* and *N. singularis* warrants further taxonomic examination, in the present study, we focused on identification and therefore did not discuss this limitation.

**Fig. 2.**
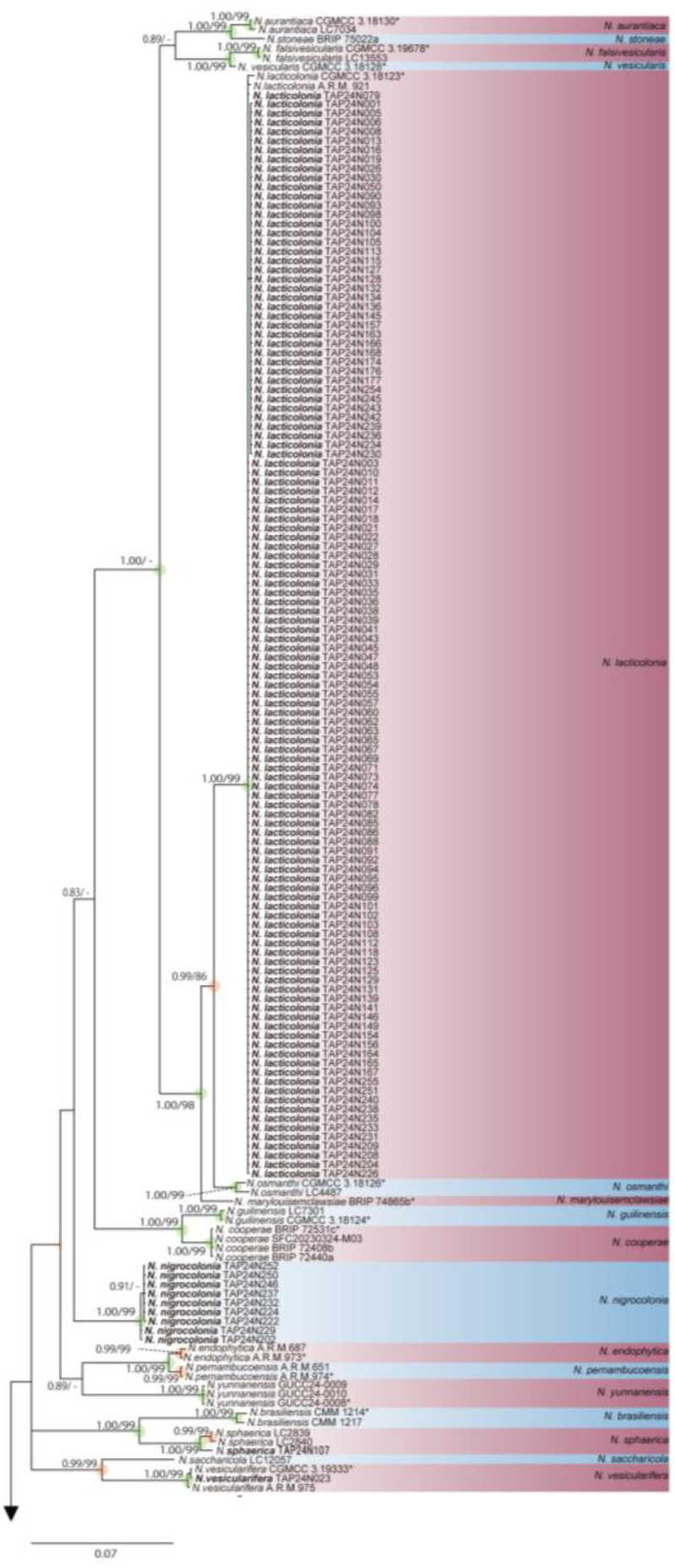
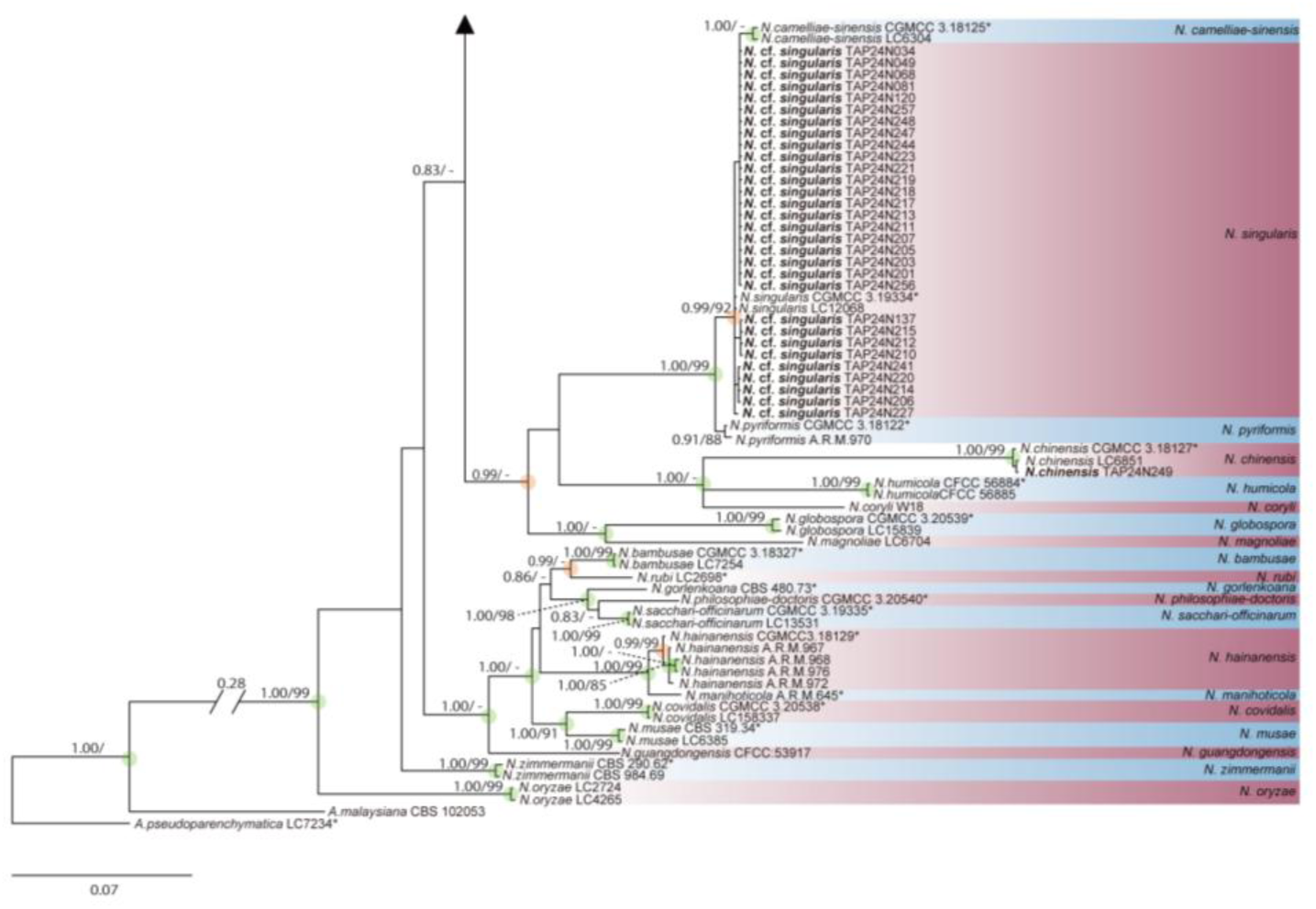
Phylogenetic trees constructed based on the concatenated data of the DNA sequences of ITS, *β-tubulin*, and *tef1α* genes inferred using MrBayes. HKY, GTR, and HKY was used as models in MrBayes for ITS, *β-tubulin*, and *tef1α*, respectively. The numbers on the left and right of each node are posterior probabilities estimated using the software MrBayes and bootstrap values from ML method performed with MEGA 10 with Tamura-Nei model, respectively. Nodes with a PPB of 1.0 are highlighted with green circles, while those with PPB ≥ 0.95 are indicated by orange circles. Strain number with asterisk is ex-type strain. Species name and strains with boldface is our isolate. Branch lengths are proportional to the number of substitutions per site; the scale bar indicates expected substitutions per site.

The phylogenetic positions of the clades to which the isolates belong are as follows. *N. chinensis* clade neighbored *N. humicola* Qin Yang & Ning Jiang (BPP/ML = 1.0/99), *N. lacticolonia* clade neighbored *N. osmanthi* Mei Wang & L. Cai (BPP = 0.99/86), *N. nigrocolonia* clade, a new lineage, neighbored a clade comprising *N. cooperae* Y.P. Tan, Bishop-Hurley, Bransgr. & R.G. Shivas, *N. guilinensis* Mei Wang & L. Cai, *N. lacticolonia* Mei Wang & L. Cai clade, *N. marylouisemclawsiae* Y.P. Tan, Ryley & Bishop-Hurley, and *N. osmanthi*, *N. sphaerica* clade neighbored *N. brasiliensis* A.C.Q. Brito, C. Conforto & A.R. Machado (BPP/ML = 1.0/99), and *N. vesicularifera* clade neighbored *N. saccharicola* M. Raza & L. Cai (BPP/ML = 0.99/99).

Phylogenetic analyses were conducted based on three DNA regions (ITS, β-tubulin, and *tef1a*; Supplementary Figure 1; see Supplementary Data Set) independently. Among the isolates, thirty strains identified as *N.* cf. *sphaerica* formed a monophyletic clade with the reference strains of *N. camelliae-sinensis*, *N. falsivesicularis* M. Raza & L. Cai, and *N. pyriformis* Mei Wang & L. Cai in the phylogenetic tree based on ITS region (ITS tree; BPP < 0.95). In contrast, these strains formed a strongly supported monophyletic clade (BPP = 1.0) with *N. camelliae-sinensis*, *N. pyriformis*, and *N. sphaerica* (Sacc.) E.W. Mason in the phylogenetic tree based on the *β-tubulin* region *(β-tubulin* tree; Supplementary Figure 2). In the phylogenetic tree based on *tef1a* (*tef1a* tree; Supplementary Figure 3), although posterior probabilities were not indicated, the isolates formed a monophyletic group with *N. singularis*.

The isolate identified as *N. chinensis* Mei Wang & L. Cai consistently formed a well-supported monophyletic clade (BPP = 1.0 for all loci) with the reference strains of *N. chinensis*, including the ex-type strain, across all ITS, β-tubulin, and *tef1a* trees (Supplementary Figure 1, 2, 3). The isolates identified as *N. lacticolonia* formed a monophyletic clade with *N. osmanthi* and *N. vesicularifera* M. Raza & L. Cai in the ITS tree; however, they clustered with the reference strains of *N. lacticolonia* (including the ex-type strain) in both the β-tubulin and *tef1α* trees (BPP = 1.0/0.83: β-tubulin/*tef1a*). Similarly, the isolate identified as *N. sphaerica* formed a monophyletic clade with the reference *N. sphaerica* (including the ex-type strain) across all loci (BPP = 0.93/1.0/1.0 for ITS/*β-tubulin*/*tef1a*). The isolate identified as *N. vesicularifera* clustered with the reference strains of *N. vesicularifera* (including the ex-type strain) in both the β-tubulin and *tef1a* trees (BPP = 1.0/1.0), while in the ITS tree, it formed a monophyletic group with both *N. vesicularifera* and *N. saccharicola* (BPP = 1.0). Although species-level resolution was not always clearly delineated in phylogenies based on single locus, all isolates were positioned within clades closely related to the corresponding identified species. Furthermore, the newly established species *N. nigrocolonia* was independent from any known species in all phylogenetic trees (BPP = 1.0/1.0/1.0: ITS/*β-tubulin* /*tef1a*).

Focusing on isolation frequency at each collection site, *N. lacticolonia* was the most frequently isolated species in Little panay, Panambon, Tugbok, and Ula. Meanwhile, *N. singuralis* M. Raza & L. Cai was the most frequently isolated species from Dabong-dabong and Patag-Alanib (Fig. 3a, b).

**Fig. 3.**
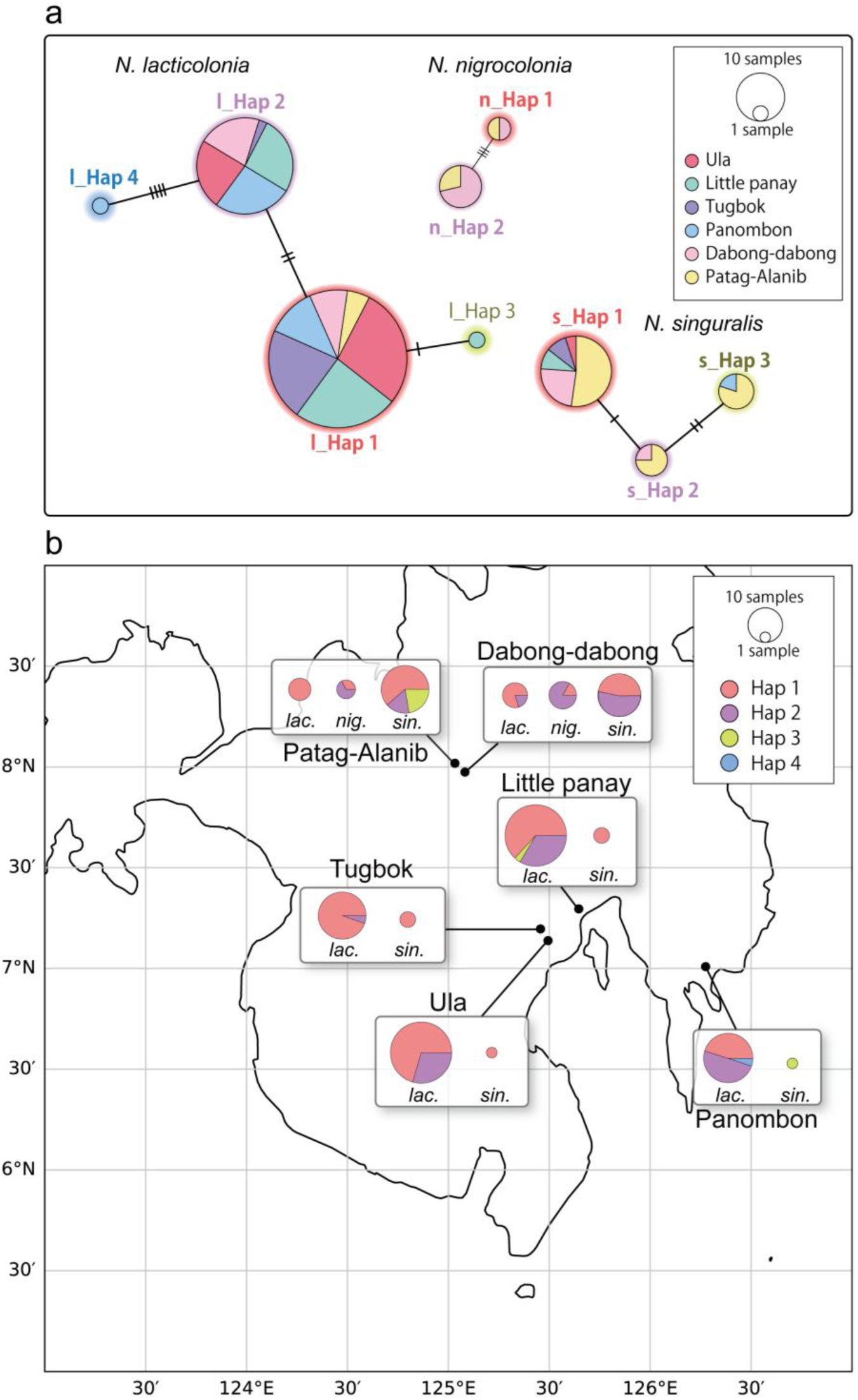
Haplotype analyses of *N. lacticolonia*, *N. niglospora*, and *N. singularis*. Haplotype networks (a). l_Hap; *N. lacticolonia* haplotype, n_Hap; *N. nigrospora* haplotype, and s_Hap; *N. singularis* haplotype. Distribution of species and haplotypes in the Mindanao Island, Philippines (b). *lac*; *N. lacticolonia*, *nig*; *N. nigrospora*, and *sin*; *N. singularis*. The size of the circle depends on the number of samples.

### Taxonomy

#### Nigrospora nigrocolonia

Nozawa & Kyoko Watan., sp. nov. (Fig. 4.; Supplementary Figure 4) MycoBank no: MB 858325.

**Fig. 4.**
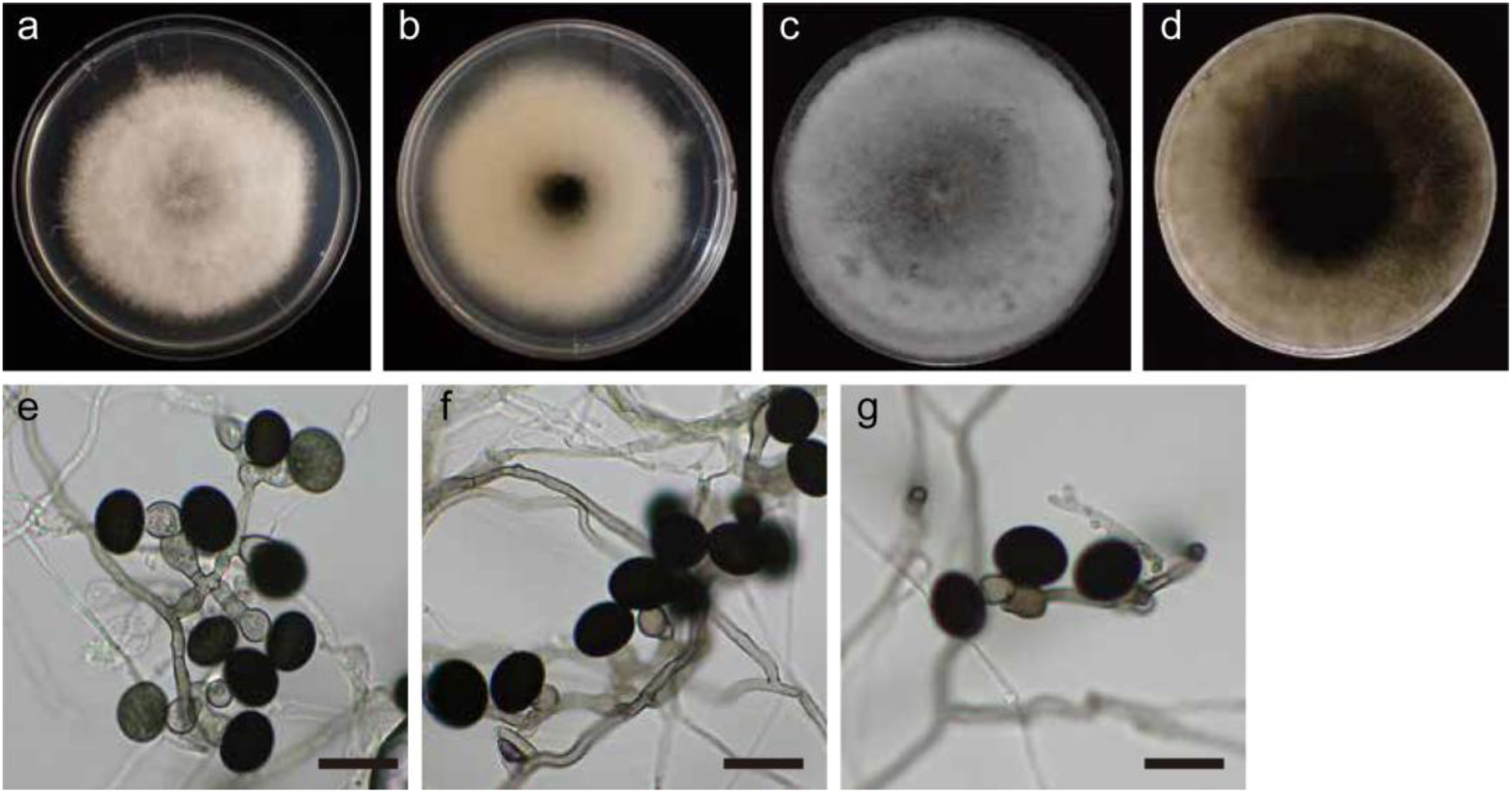
*Nigrospora nigrocolonia* (TAP24N237) Surface and reverse of colony on PDA after 3 days (a, b) and 7 days (c, d). Conidiogenous cells and conidia on SNA plate (e, f, g). Scale bars: = 20 μm

Culture characteristics: Colonies on PDA reached 90 mm in diameter. After 5 days of incubation at 25 ± 0.5 °C, the colonies had smooth margins, initially white to whitish cream with abundant cottony aerial mycelia; the reverse was initially whitish cream (Fig. 4a, b). With age, the colonies on the surface became grayish, and dark olive-to-black on the reverse (Fig. 4c, d).

Asexual morph on SNA: Hyphae are hyaline to brown, highly-branched, septate, smooth, and 2.2–4.8 (mean ± SD = 3.1±0.1, n = 30) μm wide. Conidiophores are micronematous or semi-macronematous, flexuous, olivaceous or brown, smooth (Fig. 4e), measuring 5.1– 10.6×2.5–4.3 μm (mean ± SD = 8.5±1.2×3.3±0.4, n = 4), frequently reduced to conidiogenous cells (Fig. 4e-g). Conidiogenous cells are aggregated in clusters or dispersed on hyphae, monoblastic, determinate, variable in shape, mostly subspherical to cylindrical, and olivaceous or brown, measuring 5.4–15.2×5.9–10.8 μm (mean ± SD = 8.7±0.3×7.9±0.2, n = 30). Conidia are solitary, aseptate, globose 13.2–18.7×14–18 (mean ± SD = 16.3±0.2×16.5±0.2, n = 30), and subglobose to ellipsoidal 14.4–19.2×11.7–15 (mean ± SD = 16.2±0.2×13.1±0.2, n = 30) (Fig. 4e-g). No chlamydospores were observed in this study. Sexual morphs were not observed.

Etymology: Specific epithet refers to the black coloration of the colony on the back.

Type: Dried culture specimen grown on SNA (TAP H-24N202, holotype); ex-type strain TAP24N202 isolated from diseased leaf of *Musa acuminata*, in Patag-Alanib, Lantapan, Bukidnon, Philippines, on December 11, 2024, isolated by Shunsuke Nozawa. The holotype specimen and ex-type strain were deposited in the culture collection of Tamagawa University, Japan. The GenBank accession numbers for the type strain of Nigrospora nigrocolonia are as follows: ITS: LC868094, *β-tubulin*: LC868147, and *tef1a*: LC868200.

Note: *N. nigrocolonia* forms a well-supported clade as a neighbor of a clade comprising *N. aurantiaca*, *N. cooperae*, *N. falsivesicularis*, *N. guilinensis*, *N. lacticolonia*, *N. marylouisemclawsiae*, *N. osmanthi*, *N. stoneae*, and *N. vesicularis* in the phylogenetic tree based on cancate. The holotype of *N. nigrocolonia* exhibited sequence differences with other *Nigrospora* species as follows: *N. aurantiaca* CGMCC 3.18130 showed ITS 1.1%, *β-tubulin* 12.4%, and *tef1a* 20.1%; *N. cooperae* BRIP 72440a showed ITS 1.9%, β*-tubulin* 13.0%, and *tef1a* 21.1%; *N. falsivesicularis* CGMCC3.19678 showed ITS 2.4%, *β-tubulin* 12.3%, and *tef1a* 21.4%; *N. guilinensis* CGMCC 3.18124 showed ITS 1.6%, β-tubulin 11.7%, and *tef1a* 20.5%; *N. lacticolonia* CGMCC 3.18123 showed ITS 1.6%, *β-tubulin* 9.0%, and *tef1a* 20.7%; *N. marylouisemclawsiae* BRIP 74865b showed ITS 1.3%, *β-tubulin* 9.0%, and *tef1a* 20.4%; *N. osmanthi* CGMCC 3.18126 showed ITS 1.6%, *β-tubulin* 10.0%, and *tef1a* 19.0%; *N. stoneae* BRIP 75022a showed ITS 1.1%, *β-tubulin* 15.7%, and *tef1a* 20.3%; and *N. vesicularis* CGMCC 3.18128 showed ITS 1.3%, *β-tubulin* 12.1%, and *tef1a* 20.6%.

*N. nigrocolona* differed from related species in exhibiting dark olivaceous conidiogenous cells upon maturation. The species is further distinguished by its conidial size. The conidial size of *N. nigrocolonia* was notably bigger—measuring 13.2–18.7×14–18 (mean ± SD = 16.3±0.2×16.5±0.2)—than those of *N. aurantiaca* (mean ± SD = 14.82 ± 0.79×11.78 ± 1.07 µm), *N. cooperae* (10–12.5×8.5–10.5 µm), *N. falsivesicularis* (mean ± SD = 13.62 ± 1.8 µm), *N. guilinensis* (mean ± SD = 12.46 ± 0.62 × 9.69 ± 0.71 µm), *N. lacticolonia* (mean ± SD = 14.36 ± 1.04 µm), *N. osmanthi* (mean ± SD = 14.87 ± 0.63 µm), and *N. vesicularis* (mean ± SD = 14.8 ± 0.76 µm). Morphological data for *N. marylouisemclawsiae* and *N. stoneae* are currently unavailable and were therefore excluded from conidial comparisons.

In addition to the type strain identified as *N. nigrocolonia*, the following isolates— TAP24N222, TAP24N224, TAP24N229, TAP24N232, TAP24N237, TAP24N246, TAP24N250, and TAP24N252—have been deposited in the Tamagawa University Mycological Bio-bank of Plant Diseases.

### Morphological descriptions of isolates

In this section, species other than *N. nigrocolonia* isolated in this study are described. For species from which multiple strains were obtained, one strain was randomly selected for morphological characterization.

#### Nigrospora cf. singularis

Isolate no. TAP24N068: Culture characteristics: Colonies on PDA reached 90 mm in diameter. After 4 days of incubation at 25 ± 0.5 °C, the colonies had smooth margins and were white to whitish gray with abundant cottony aerial mycelia; the reverse was initially whitish cream (Fig.5a, b). Asexual morph on SNA: Hyphae are hyaline to pale brown, branched, septate, smooth, and 1.2–5.1 (mean±SD = 2.4±0.13, n = 30) μm wide. Conidiophores are micronematous or semi-macronematous, flexuous, hyaline, and smooth. Conidiogenous cells are dispersed on hyphae, monoblastic, determinate, variable in shape, mostly subspherical to oblate, hyaline, measuring 6.4–16.1×6.4–17.3 (mean±SD = 9.4±0.41×8.1±0.35, n = 30) in size (Fig. 5c, d). Conidia are solitary, aseptate, and subglobose to ellipsoidal 9.8–13×9–13 (mean±SD = 11.6±0.18×11±0.22, n = 30; Fig.4c, d). No chlamydospores were observed in this study. Sexual morphs were not observed. The size of the conidia of this strain closely matched the original description of *N. singularis* (CGMCC 3.19334) ^13^: 9.5–13 × 11–15 (mean ± SD = 11.2 ± 1.11 × 13.4 ± 1.12) μm.

**Fig. 5.**
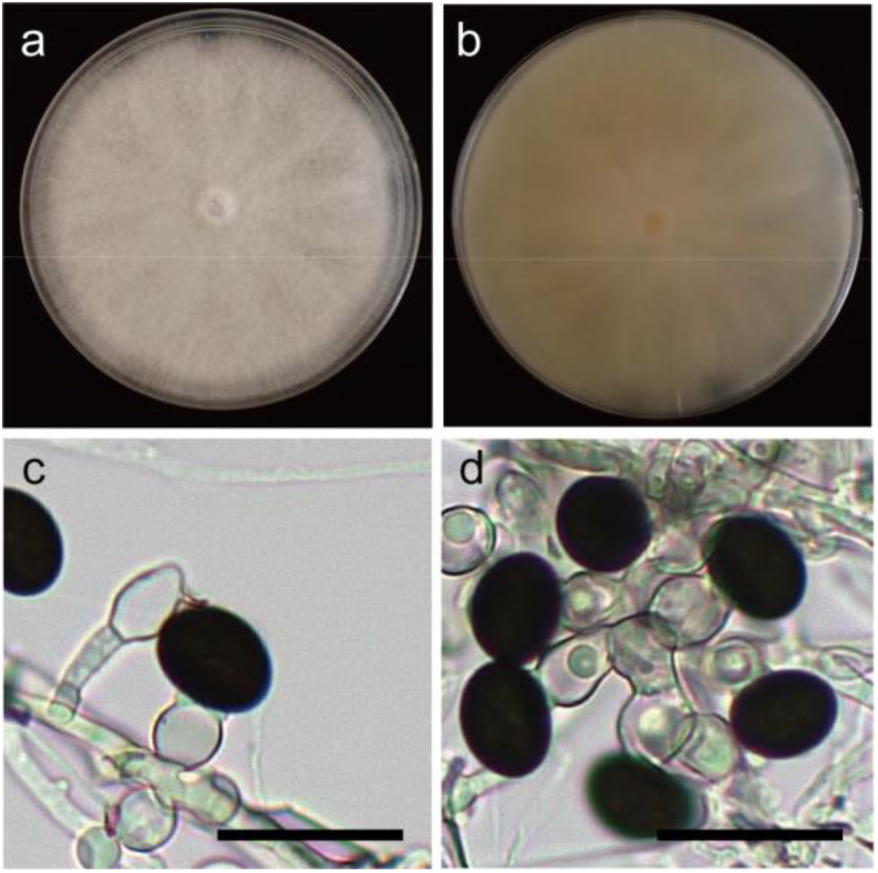
*Nigrospora chinensis* (TAP24N249) Surface and reverse of colony on PDA after 3 days (a, b). Conidiogenous cells and conidia on SNA plate (c, d). Scale bars: = 20 μm

#### N. chinensis

Isolate no. TAP24N249: Culture characteristics: Colonies on PDA reached 90 mm in diameter. After 7 days of incubation at 25 ± 0.5 °C, the colonies had smooth margins and were white to whitish cream, with abundant cottony aerial mycelia. the reverse was whitish gray to pale brown (Fig. 6a, b). Asexual morph on SNA: Hyphae are hyaline to pale brown, highly-branched, septate, smooth, and 1.5–4.6 (mean±SD = 2.7±0.14, n = 30) μm wide. Conidiophores are micronematous or semi-macronematous, flexuous, hyaline, smooth, frequently reduced to conidiogenous cells (Fig. 6c, d). Conidiogenous cells are dispersed on hyphae, monoblastic, determinate, variable in shape, mostly subspherical to oblate, hyaline, and 4.8–12.6×6.9–10.5 (mean±SD = 8.8±0.31×8±0.16, n = 30) in size. Conidia are solitary, aseptate, and subglobose to ellipsoidal 10.5–15.6×7.8–13 (mean±SD = 12.8±0.23×10.2±0.2, n = 30; Fig. 6c, d). No chlamydospores were observed in this study. Sexual morphs were not observed. The size of conidia of this strain closely matched the original description of *N. chinensis* (CGMCC 3.18127)^14^: 10–14.5 × 7.5–11 (mean±SD = 11.78 ± 0.75 × 9.18 ± 0.61) μm.

**Fig. 6.**
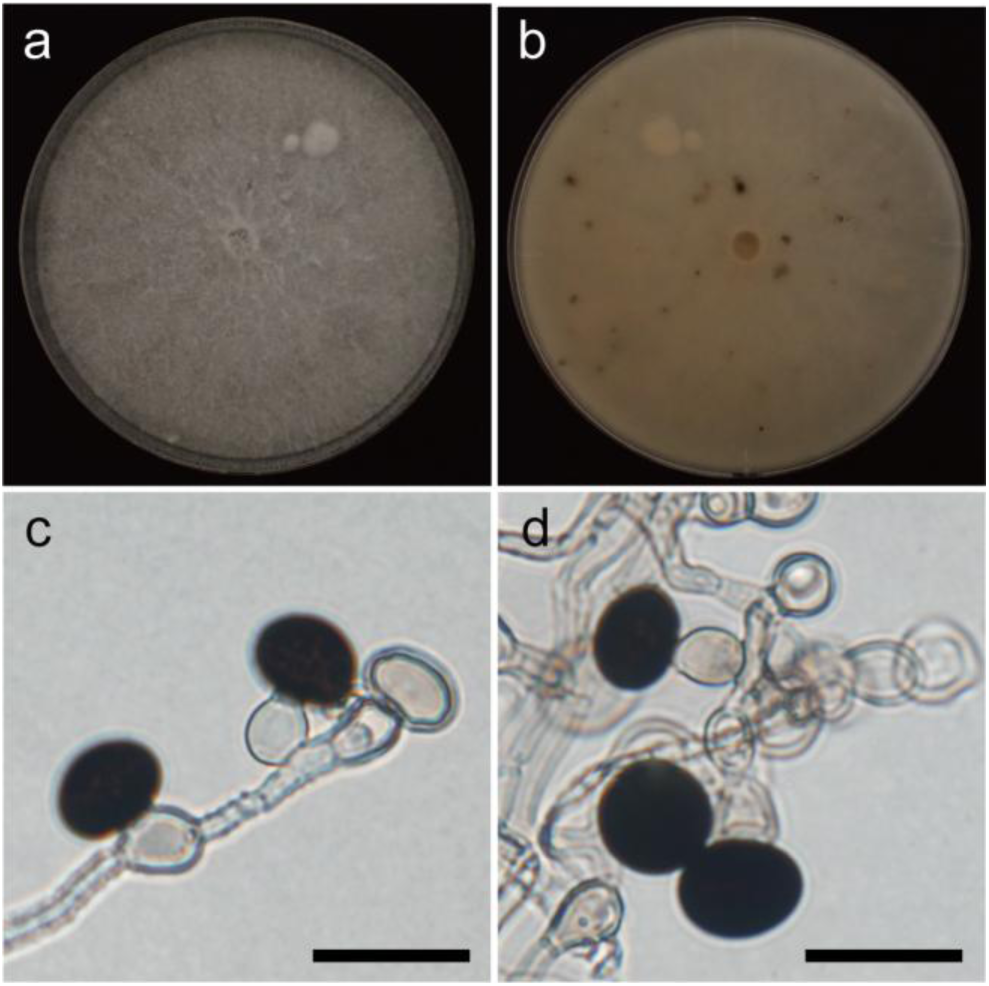
*Nigrospora lacticolonia* (TAP24N011) Surface and reverse of colony on PDA after 3 days (a, b). Conidiogenous cells and conidia on SNA plate (c, d). Scale bars: = 20 μm

#### N. lacticolonia

Isolate no. TAP24N011: Culture characteristics: Colonies on PDA reached 90 mm in diameter. After 3 days of incubation at 25 ± 0.5 °C, the colonies had smooth margins, and were white to whitish gray with abundant cottony aerial mycelia; the reverse was whitish cream to pale brown (Fig. 7a, b). Asexual morph on SNA: Hyphae are hyaline to pale brown, highly-branched, septate, smooth or verrucose, and 2–5.8 (mean±SD = 3.7±0.16, n = 30) μm wide. Conidiophores are micronematous or semi-macronematous, flexuous, hyaline, smooth or verrucose, frequently reduced to conidiogenous cells. Conidiogenous cells are dispersed on hyphae, monoblastic, determinate, variable in shape, mostly subspherical to oblate, hyaline, and 4.6–14.4×5.7–10.9 (mean±SD = 8.2±0.35×7.8±0.2, n = 30) in size (Fig. 7c, d). Conidia are solitary, aseptate, subglobose to ellipsoidal 13.2–16.5×12.2–16.1 (mean±SD = 15.3±0.17×14.9±0.15, n = 30; Fig 7c, d). No chlamydospores were observed in this study. Sexual morphs were not observed. The size of conidia of this strain closely matched the original description of *N. lacticolonia* (CGMCC 3.18123) [14]: 11.5–16.5 (mean±SD. = 14.36 ± 1.04) μm diameter.

**Fig. 7.**
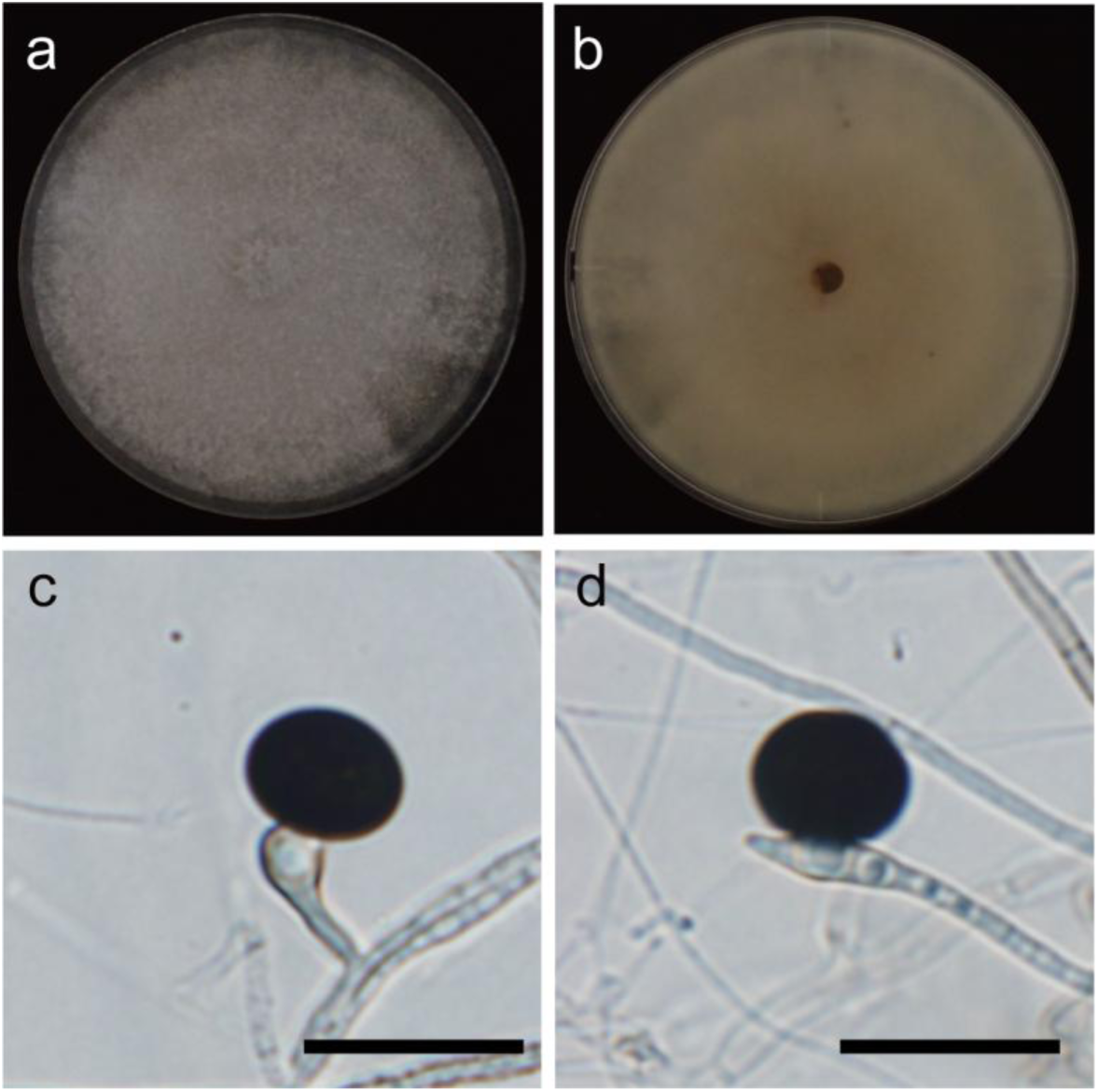
*Nigrospora singuralis* (TAP24N068) Surface and reverse of colony on PDA after 3 days (a, b). Conidiogenous cells and conidia on SNA plate (c, d). Scale bars: = 20 μm

#### Nigrospora sphaerica

Isolate no. TAP24N107: Culture characteristics: Colonies on PDA reached 90 mm in diameter. After 3 days of incubation at 25 ± 0.5 °C, the colonies had smooth margins, and were white to whitish gray with abundant cottony aerial mycelia; the reverse was whitish cream (Fig. 8a, b). Asexual morph on SNA: Hyphae are hyaline to pale brown, branched, septate, smooth, and 1.9–4.2 (mean±SD = 3.1±0.12, n = 30) μm size. Conidiophores are micronematous or semi-macronematous, flexuous, hyaline, smooth, and frequently reduced to conidiogenous cells. Conidiogenous cells are dispersed on hyphae, monoblastic, determinate, variable in shape, mostly subspherical to oblate, hyaline to pale brown, and 6.6–11.7×5.8–17.2 (mean±SD = 9.3±0.23×8.7±0.38, n = 30) in size (Fig. 8c, d). Conidia are solitary, aseptate, subglobose to ellipsoidal 15.6–22.6×15.2–22.9 (mean±SD = 19.6±0.33×19.5±0.36, n = 30; Fig. 8c, d). No chlamydospores were observed in this study. Sexual morphs were not observed. The size of conidia of this strain closely matched the description of *N. sphaerica* (LC2840) measured by Wang et al. [14]: 16–21 (mean±SD = 18.22 ± 1.0) μm diameter.

**Fig. 8.**
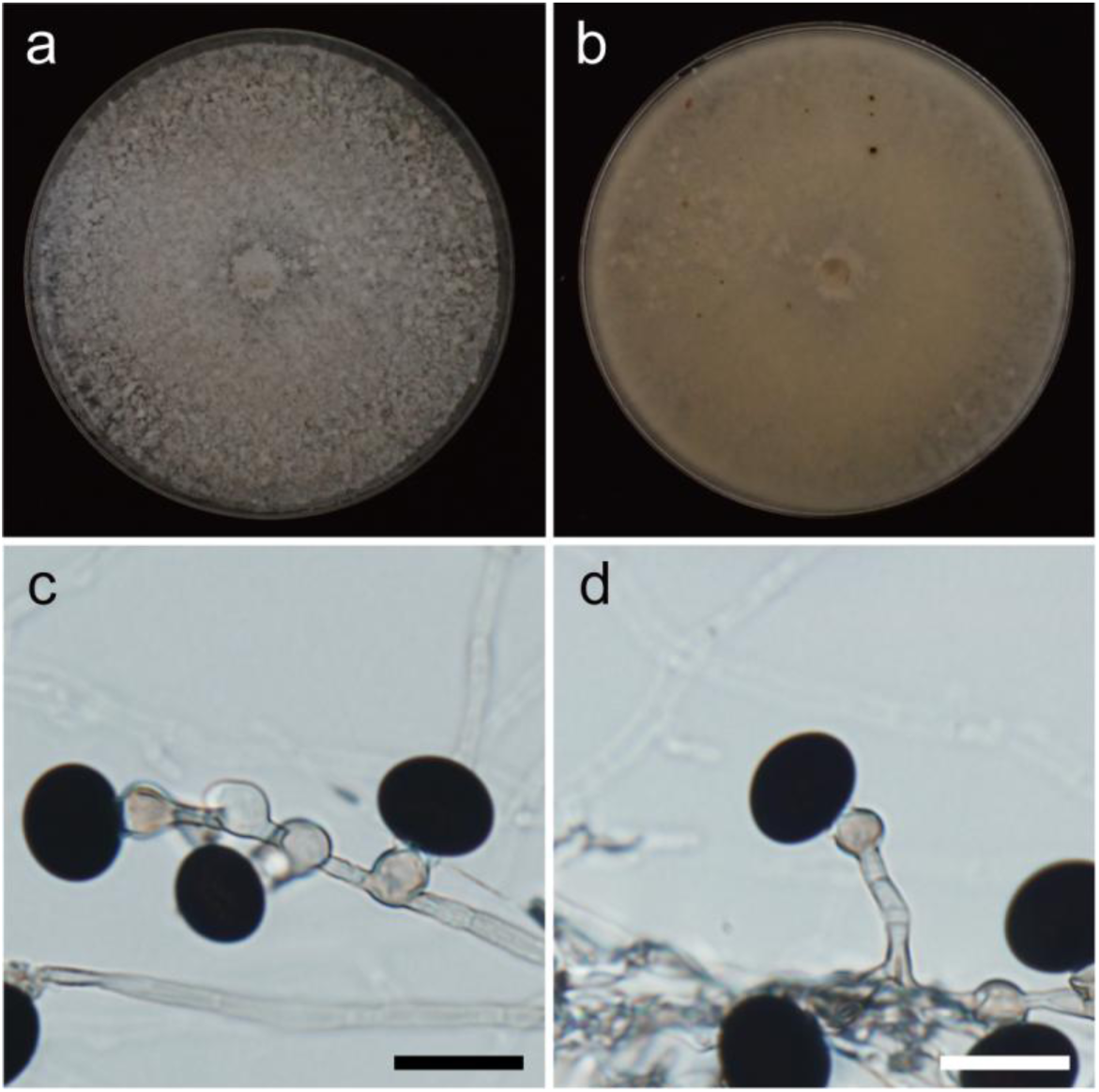
*Nigrospora sphaerica* (TAP24N107) Surface and reverse of colony on PDA after 3 days (a, b). Conidiogenous cells and conidia on SNA plate (c, d). Scale bars: = 20 μm

#### N. vesicularifera

Isolate no. TAP24N023: Culture characteristics: Colonies on PDA reached 90 mm in diameter. After 3 days of incubation at 25 ± 0.5 °C, the colonies had smooth margins, and were white to whitish gray with abundant cottony aerial mycelia; the reverse was whitish cream to brown (Fig. 9a, b). The isolate did not produce conidia. Sexual morphs were not observed.

**Fig. 9.**
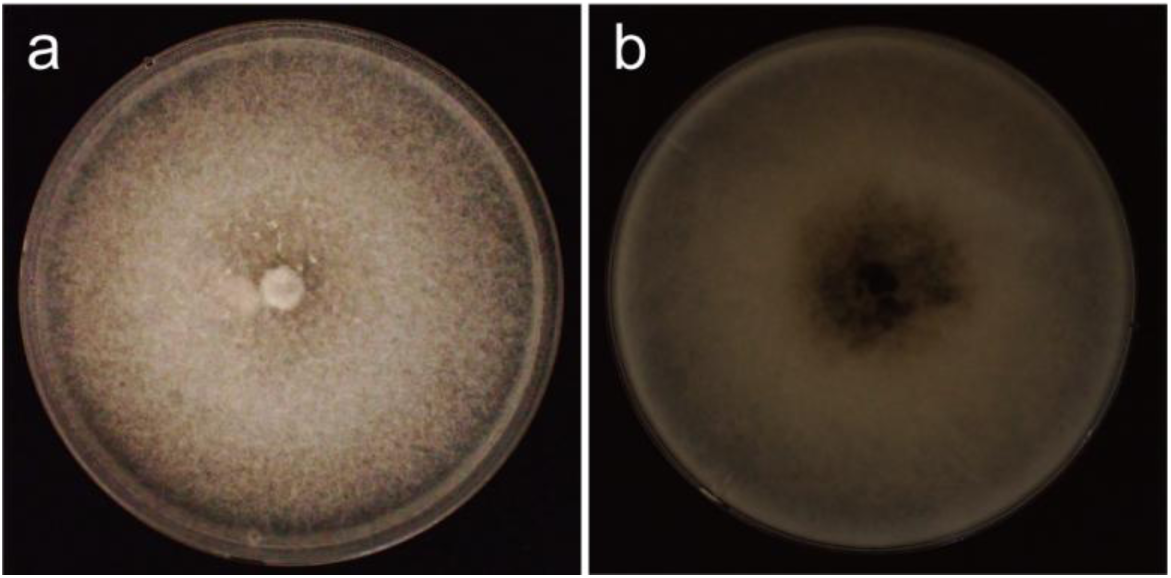
*Nigrospora vesicularifera* (TAP24N023) Surface and reverse of colony on PDA after 3 days (a, b).

### Haplotype analyses

Haplotype analysis based on the *tef1α* gene sequences was conducted for *N. lacticolonia*, *N. nigrocolonia*, and *N. singularis*, which were obtained as multiple isolates (Fig. 3a, b). *N. lacticolonia* included 118 isolates, and the alignment consisted of 616 sites. This species was divided into four haplotypes, characterized by six variable sites (S) and a nucleotide diversity (Pi) of 0.001586. Among all the isolates, 78 belonged to Hap_1, 38 to Hap_2, and 1 each to Hap_3 and Hap_4. Hap_1 was found at all six collection sites, whereas Hap_2 was found at five collection sites, excluding Patag-Alanib. Hap_3 was isolated solely from Little Panay, and Hap_4 was isolated solely from Panombon.

*N. nigrocolonia* included 9 isolates, and the alignment consisted of 714 sites. This species was divided into two haplotypes, with three variable sites (S) and a nucleotide diversity (Pi) of 0.001634. Two isolates belonged to Hap_1, and seven to Hap_2. Both haplotypes were isolated from Dabong-dabong and Patag-Alanib cultivars.

*N. singularis* included 30 isolates, and the alignment consisted of 604 sites. This species was divided into three haplotypes, with three variable sites (S) and a nucleotide diversity (Pi) of 0.001254. Among all isolates, 21 belonged to Hap_1, 4 to Hap_2, and 5 to Hap_3. Hap_1 was found at five of the collection sites, excluding Panombom, whereas Hap_2 was found at Dabong-dabong and Patag-Alanib. Hap_3 was isolated from the panomboms and patag-alanib.

### Pathogenicity to leaves

All 160 isolates were tested on wounded and non-wounded banana leaves. The aggressiveness of the tested isolates on wounded leaves differed significantly, with lesion diameters ranging from 2.33 mm to 22.61 mm (Fig. 10; Table 3; Supplementary Figure 5). Based on diameter sizes of the disease spots, aggressiveness was classified as strongly virulent (≧6.6 mm), moderately virulent (≧3 to <6.6 mm), weakly virulent (> 0 to <3 mm), or very weakly virulent (0 mm with brown or yellow discoloration at or around the wound site). Among the 160 isolates investigated, 54 (33.75%) were strongly virulent, 52 (32.5%) moderately virulent, 43 (26.9%) weakly virulent, and 8 (5%) very weakly virulent; 2 (1.25%) isolates were non-pathogenic. No symptoms appeared on non-wounded banana leaves.

**Fig. 10.**
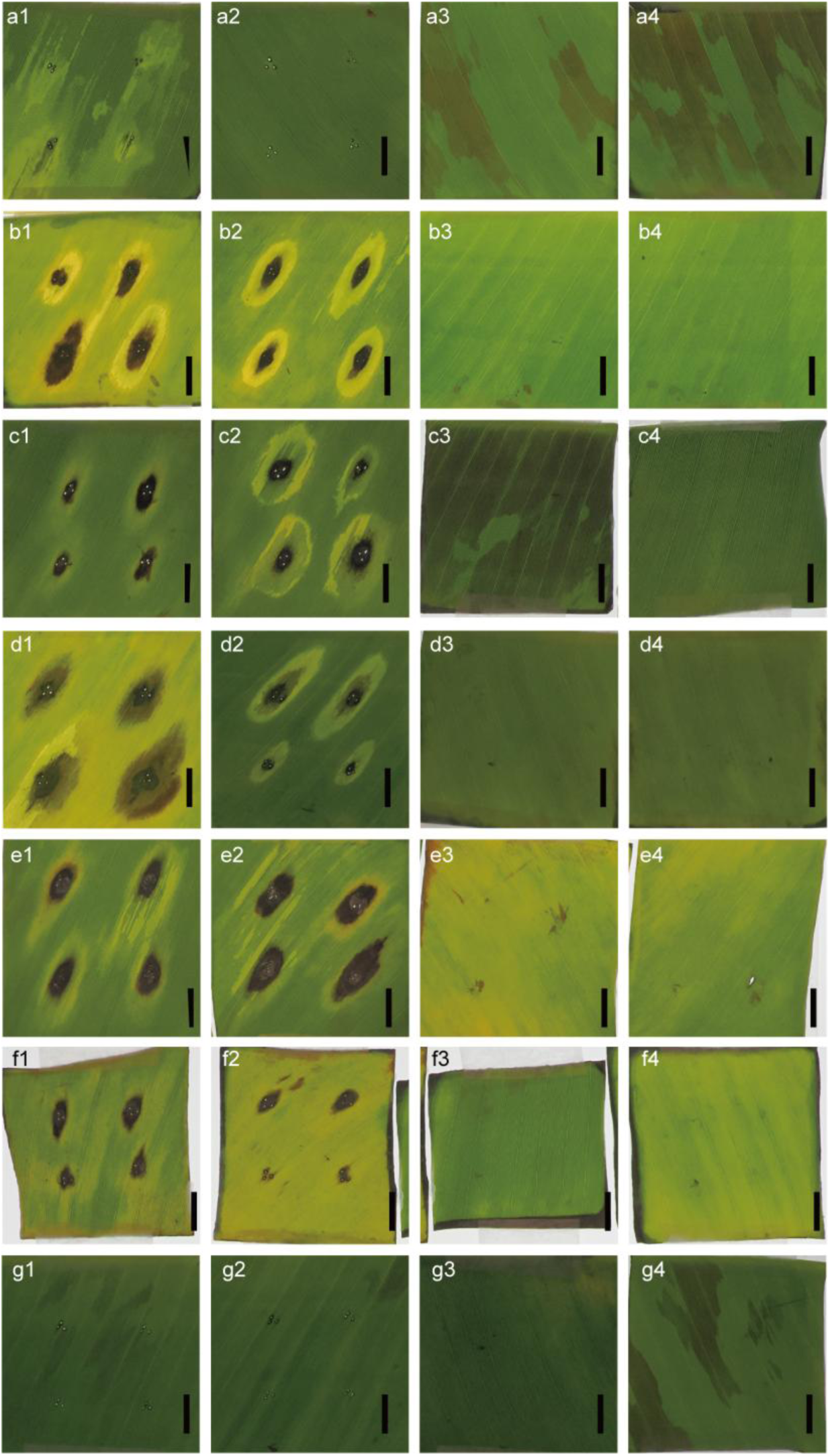
Samples of banana leaves 4 days after inoculation with mycelial plugs. Inoculated leaves with *N. chinensis* (isolate TAP24N011) with wounds (a1, a2) and without wound (a3, a4), *N. lacticolonia* (isolate TAP24N078) with wounds (b1, b2) and without wound (b3, b4), *N. nigrocolonia* (isolate TAP24N237) with wounds (c1, c2) and without wound (c3, c4), *N.* cf. *singularis* (isolate TAP24N257) with wounds (d1, d2) and without wound (d3, d4), *N. sphaerica* (isolate TAP24N107) with wounds (e1, e2) and without wound (e3, e4), *N.vesicularifera* (isolate TAP24N023) with wounds (f1, f2) and without wound (f3, f4). Controls inoculated with potato dextrose agar discs with wounds (g1, g2) and without wound (g3, g4). Bars: 1 cm.

These 160 isolates included six *Nigrospora* species: *N. chinensiss*, *N. lacticolonia*, *N.* cf. *singularis*, *N. sphaeric*a, *N. vesicularifera*, and *N. nigrocolonia*. Among them, *N. chinensis*, *N. sphaerica*, and *N. vesicularifera* were each represented by a single isolate, while multiple isolates were available for *N. lacticolonia*, *N.* cf. *singularis*, and *N. nigrocolonia*. The single isolates of *N. chinensis* exhibited weak virulence. *N. sphaerica*, and *N. vesicularifera* exhibited strong virulence. In contrast, isolates of *N. lacticolonia*, *N.* cf. *singularis*, and *N. nigrocolonia* showed variable levels of aggressiveness. Among the *N. lacticolonia* isolates, 39.8% (47/118 isolates) were classified as strongly virulent, 37.3% (44/118 isolates) as moderately virulent, 18.6 (22/118 isolates) as weakly virulent, 2.5% (3/118 isolates) as very weakly virulent, and 1.7% (2/118 isolates) as non-pathogenic. For *N.* cf. *singularis*, 16.7% (5/30 isolates) were classified as strongly virulent, 20% (6/30 isolates) as moderately virulent, 50% (15/30 isolates) as weakly virulent, and 13.3% (4/30 isolates) as very weakly virulent. For the isolates of *N. nigrocolonia*, 11.1% (1/9 isolates) were classified as strongly virulent, 22.2% (2/9 isolates) as moderately virulent, 55.5% (5/9 isolates) as weakly virulent, and 11.1% (1/9 isolates) as very weakly virulent.

### In vitro fungicide sensitivity test

The heat map in Fig. 11 presents the sensitivity of each isolate of *Nigrospora lacticolonia* (118 isolates), *N. singularis* (30 isolates), *N. nigrocolonia* (9 isolates), *N. chinensis*, *N. sphaerica,* and *N. vesicularifera* (1 isolate each) to each fungicide (active ingredient), using the rate of inhibition of mycelial growth as an indicator (Supplemental Table 1).

**Fig. 11.**
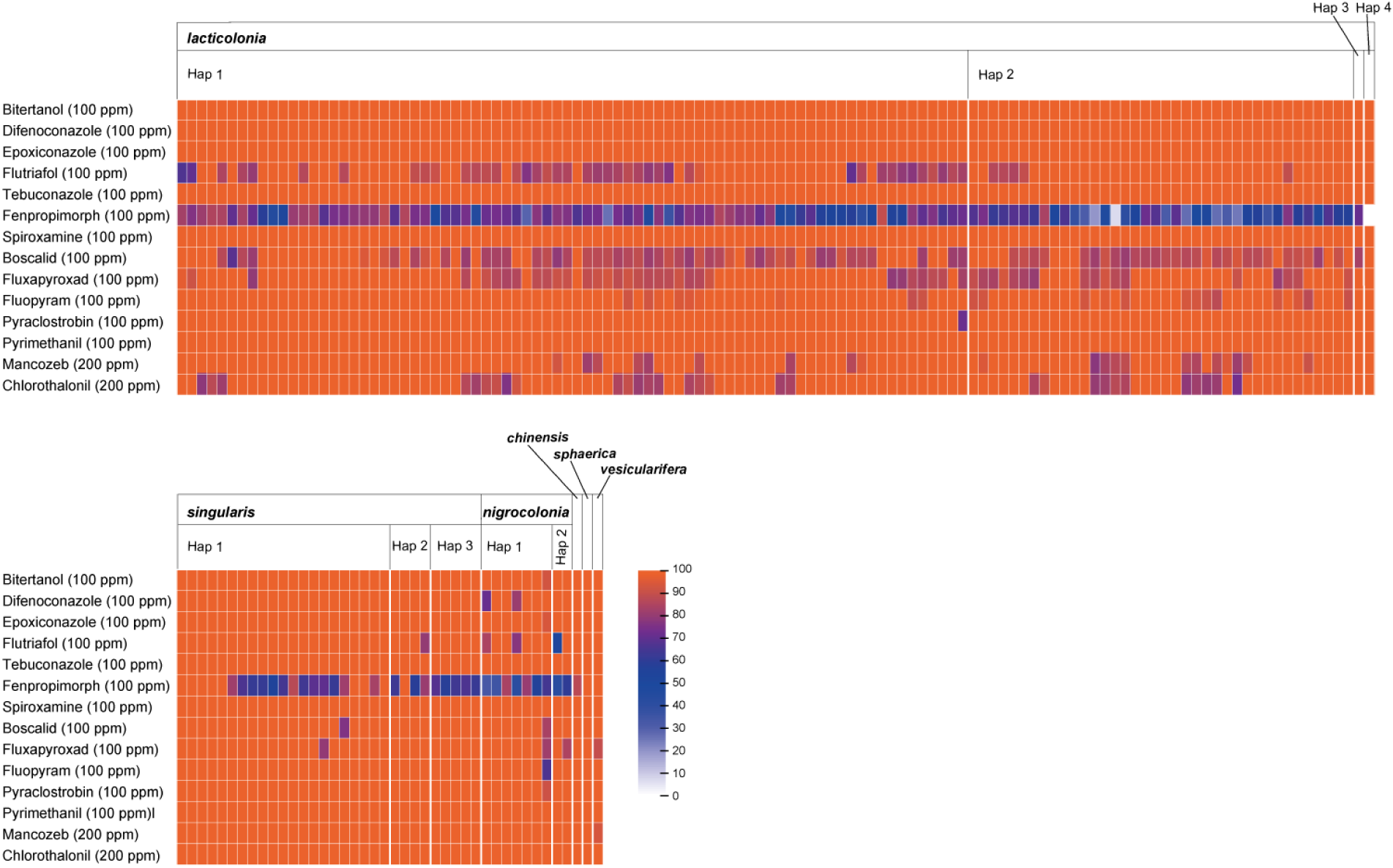
Heat map of the inhibition percentages for the isolates.

All the isolates exhibited high sensitivity (100% inhibition rate) to pyrimethanil (AP) and tebuconazole (DMI). Against pyraclostrobin (QoI), one isolate of *N. lacticolonia* and one isolate of *N. nigrospora* showed less than 100% sensitivity (72.7% and 92.3%, respectively). Twenty-one isolates were less than 100% (78.7-96.7%) sensitive to chlorothalonil, and 30 isolates less than 100% (74.4-88.5%) sensitive to chlorothalonil. Both fungicides resulted in 100% inhibition of the remaining isolates. Seventy-seven isolates of *N. lacticolonia* exhibited less than 100% (70.1-86.6%) sensitivity to boscalid (SDHI). One isolate (76.9%) of *N. singularis* and one isolate (84.7%) of *N. nigrocolonia* exhibited less than 100% sensitivity. Sixty-nine isolates of *N. lacticolonia* exhibited a range of 79.9–98% sensitivity to fluxapyroxad. One isolate (80.6%) of *N. singularis* and two (83.2-85.8%) of *N. nigrocolonia* had less than 100% sensitivity. Twenty-two isolates of *N. lacticolonia* displayed sensitivities ranging from 88.8–98.1% to fluopyram. One isolate (67.9%) of *N. nigrocolonia* exhibited less than 100% sensitivity. All isolates were highly sensitive to the GMI agents bitertanol, difenoconazole, and epoxiconazole, except for one to two isolates of *N. nigrocolonia* (72-94.8% inhibition rate). With the exception of nine isolates of *N. singularis* and one isolate each of *N. sphaerica* and *N. vesicularifera*, the inhibition rates of fenpropimorph (SBI class 2) were less than 100% (16.9-90.7%). In contrast, all fungi were highly sensitive (100%) to spiroxamine, which belongs to the same class as fenpropimorph.

The dominant species, *N. lacticolonia*, exhibited low sensitivity to 8 of the 14 fungicides. The second most dominant species, *N. singularis*, presented low sensitivity to the four species. *N. nigrocolonia* exhibited low sensitivity to nine fungicides, while *N. chinensis* and *N. vesicularifera* exhibited low sensitivity to one.

### Principal component analysis (PCA) for fungicide sensitivety

PCA was performed based on inhibition rates (Supplemental Table 1). In the final PCA results, data for TAP24N093, TAP24N094, TAP24N163, and TAP24N252, which appeared as outliers, were excluded (Supplementary Figure 6). These outliers were determined based on principal component scores obtained from PCA, using a threshold of three standard deviations from the mean for PC1 and PC2. This criterion is grounded in the statistical property of the normal distribution, where approximately 99.7% of data points fall within ±3 standard deviations. PC1 and PC2 together explained 37.99% of the total variance in the dataset (Supplementary Figure 7; Supplementary Table 2). For PC1, the top three contributors were Chlorothalonil (22.7%), Mancozeb (21.6%), and Fluopyram (16.5%). For PC2, the top three contributors included Flutriafol (53.8%), Fluxapyroxad (18.8%), and Difenoconazole (14.8%). Cluster analysis, based on the distances between the coordinates in the two-dimensional scatter plot of PC1 and PC2, revealed that no distinct clusters were formed within the specific regions or fungal lineages (Fig. 12). In cases where PCA was performed including the four strains identified as outliers, no clustering by geographic region or haplotype was observed, consistent with the results obtained after excluding the outliers (Supplementary Figure 8).

**Fig. 12.**
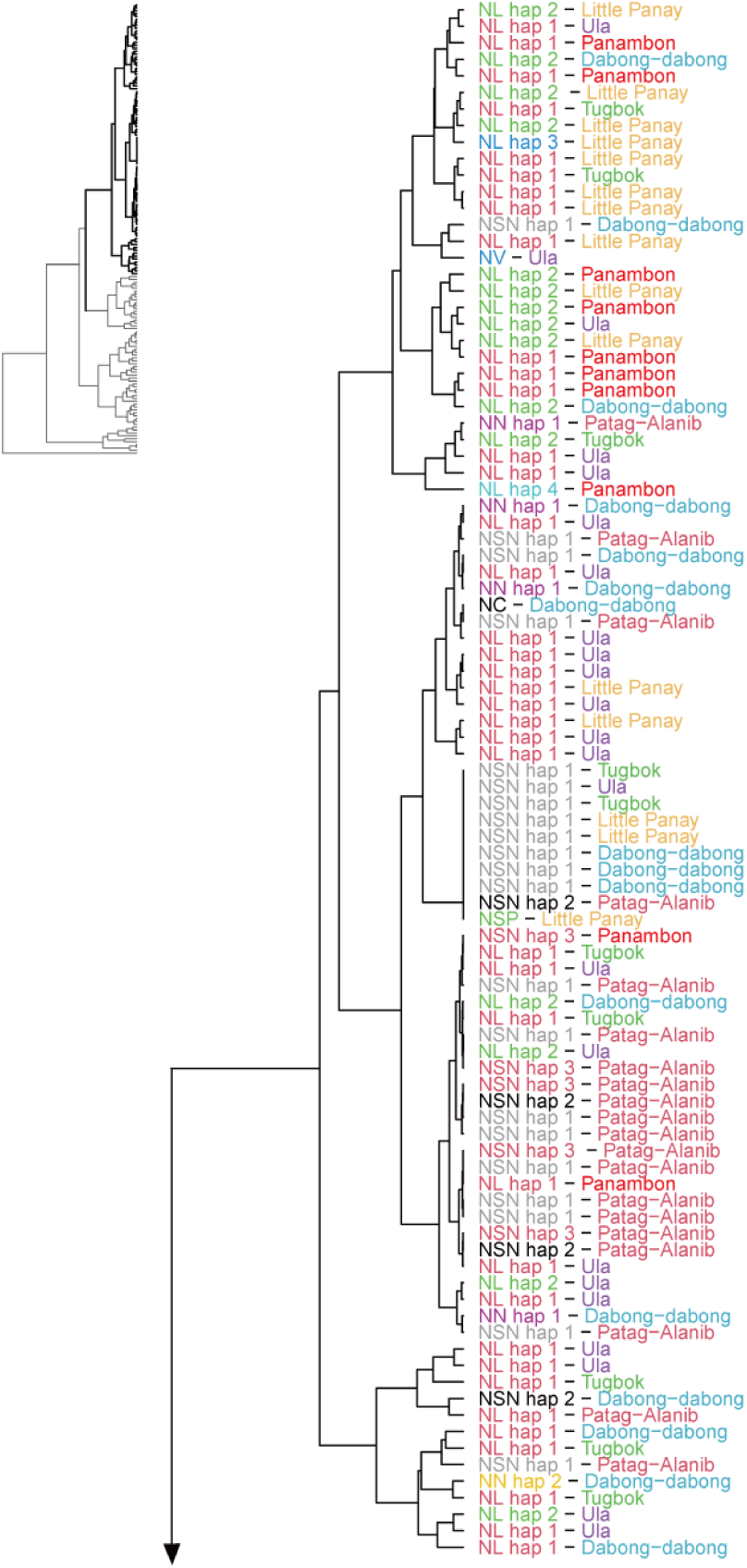
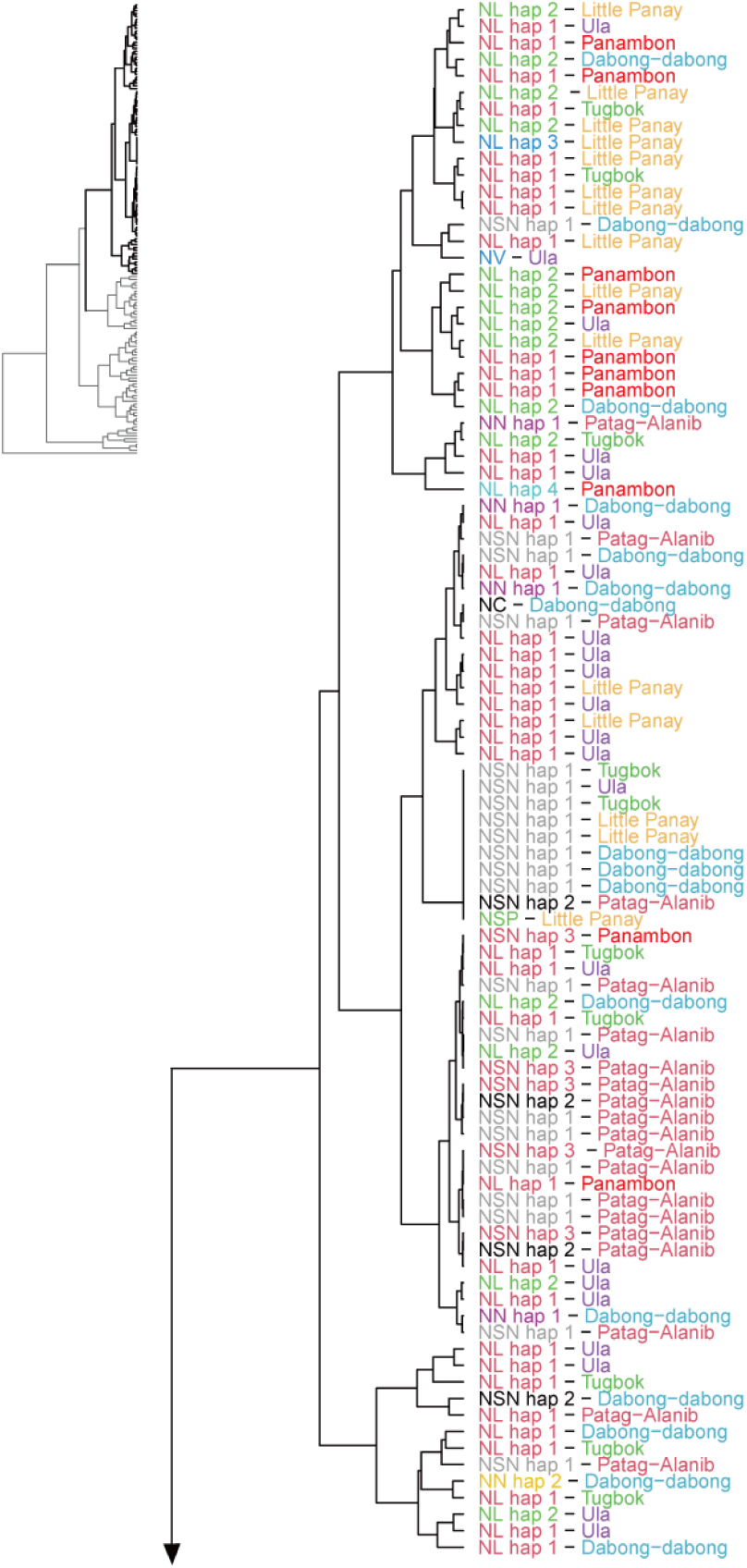
Dendrogram based on coordinates on scatter plot of PCA derived from values of inhibition rates of fungicide. Label on the left and right of hyphen are species_haplotype and sampling site, respectively.

## Discussion

This study aimed to provide essential information for managing leaf spot diseases in the Philippines. To this end, we identified pathogens responsible for banana leaf diseases in Mindanao, the main banana-producing region in the Philippines, and evaluated the efficacy of 14 fungicides currently in use against these pathogens.

On Mindanao Island, the disease “black Sigatoka,” caused by *Pseudocercospora fijiensis*, was suspected but not detected in any of the sampled locations. Eighteen genera of fungi from symptomatic leaves were collected from six farms in Mindanao, and the most prevalent genus was *Nigrospora* spp., accounting for 72% of the total fungal isolates.

The isolated isolates were identified as six species: *N. chinensis*, *N. lacticolonia*, *N. nigrocolonia*, *N. singularis*, *N. sphaerica*, and *N. vesicularifera*, all of which are pathogens that infect banana leaves. To the best of our knowledge, this is the first report of *N. chinensis*, *N. lacticolonia*, *N. nigrocolonia*, *N. singularis*, and *N. vesicularifera* as causative agents of banana leaf spot worldwide. *N. sphaerica* has previously been reported as the causal agent of banana “leaf spot” in Bangladesh^7^, but this is the first report of *N. sphaerica* in the Philippines.

In the Philippines, chemical spraying is a common agricultural practice employed 50–70 times per year to manage the local incidence of Sigatoka disease. Frequent use of fungicides may have altered the leaf biota community and created a selective pressure that allowed for the persistence of resistant species or strains. However, this cross-sectional study cannot establish causality. *Pseudocercosporaceae* fungi are inherently slow-growing; therefore, during this study, they may have been overgrown by other co-infecting fungi and remained undetected. Therefore, the presence of *P. fijiensis* at the survey sites is not discussed in this study. Nevertheless, it should be hypothesized that *P. fijiensis* may have been subjected to selection pressure from fungicide application, potentially contributing to the emergence of the genus *Nigrospora* as the dominant pathogen at the sampling sites. Future verification through pathogen-specific PCR detection on infected leaves is considered necessary.

An example of consequence fungicide application contributing to changes in the fungal composition of pathogenic communities has been reported in wheat [15]. This raises concerns regarding the selection of fungicides, as they may lead to a shift in the pathogenic fungi present on banana leaves. To ensure the continued efficacy of fungicide applications in banana cultivation, conducting regular surveys on disease incidence and fungicide sensitivity, specifically targeting *Nigrospora* and *Pseudocercospora* species, is essential.

Among the 160 isolates of the six species of the genus *Nigrospora*, *N. lacticolonia* accounted for 74% of the total (118 isolates). This species was identified as the dominant species at four sites: Little Panay, Panambong, Tugbok, and Ula. Conversely, *N. singularis* was dominant in Patag-Alanib and Dabong-dabong. The isolation frequencies of *N. nigrocolonia*, *N. sphaerica*, and *N. vesicularifera* were comparatively lower than those of the two species, and they are not currently considered primary pathogens of banana leaf spot.

Haplotype analysis revealed that *N. lacticolonia* Hap_1 and Hap_2 were detected in all six regions and five regions, respectively, while *N. singularis* Hap_1 was detected in five regions. The observation that the same haplotype was detected in multiple regions suggests that genetically similar populations are widely distributed across Mindanao Island, indicating past population migration and diffusion.

Furthermore, a comprehensive analysis of the sensitivity to 14 chemical substances using principal component analysis (PCA) revealed that the distribution of PCA on the graph did not reflect the classification of fungal species or haplotypes. Isolates exhibiting different sensitivities to 11 chemicals (excluding tebuconazole, spiroxamine, and pyrimethanil) were found within the species and haplotypes. Notably, the distribution of the isolates did not align with the classification of species or haplotypes. This indicates that fungicide sensitivity among the isolates may have evolved independently of species or haplotype differentiation. This hypothesis is further supported by the findings of Hartmann et al. [16], who demonstrated that mutations in CYP51, linked to azole fungicide resistance, and Cytb, associated with strobilurine (QoI) resistance, emerged independently in distinct geographical regions. Therefore, this observation suggests that the fungicide sensitivity in the isolates arose independently of species and haplotype differentiation. However, because this study did not include a genetic analysis of fungicide resistance, the ensuing discussion focuses on the potential mechanisms underlying the acquisition of fungicide resistance. Hartmann et al. [16] noted that gene flow resulting from human migration and spore dispersal contributes to the ectopic occurrence of resistant isolates. The dispersal of diseases caused by *Nigropsora* spp. on Mindanao Island is not only attributable to human migration resulting from the movement of soil and seedlings but also to soil runoff caused by flooding and atmospheric migration. The potential contribution of soil runoff is supported by the detection of this genus in both soil [17] and atmospheric environments [15]. Consequently, it can be posited that fungi with acquired resistance have a propensity for dissemination.

Therefore, elucidating the mechanisms underlying the dissemination of pathogenic fungi is imperative for effective disease control. This includes controlling resistant isolates and determining the optimal timing for disinfectant application. Meredith [18] demonstrated that the spores of *Nigrospora* spp. are released into the atmosphere in a diurnal cycle. Specifically, in Jamaica, spore release commenced at approximately 7:00 a.m., reaching a peak between 8:00 and 10:00 a.m. This diurnal pattern suggests that *Nigrospora* spp. employ a release mechanism that functions under decreasing vapor pressure [19], which is crucial information for disease control. However, further research is necessary to investigate the environmental factors associated with disease outbreaks in banana plantations in the Philippines.

The findings of this study regarding pathogen sensitivity to fungicides are fundamental data. However, as this is an urgent issue, it is important to note that all 118 isolates of *N. lacticolonia*, the most frequently isolated fungus species, exhibited moderate or low sensitivity to fenpropimorph (inhibition rate 16.9–90.7%). The fungicide is unlikely to be effective because the low in vitro sensitivity suggests that its inhibitory capabilities in the field will likely remain low. In contrast, all fungi showed high sensitivity to spiroxamine, a fungicide of the same class (SBI classification II group), with 100% inhibition of mycelial growth. Notably, the use of these fungicides raises concerns regarding potential cross-resistance, and it is possible that fungi resistant to spiroxamine may emerge in the future. Based on these results, we propose that the emergence of cross-resistant fungi should be monitored.

In this study, we showed that banana leaf diseases occurring in Mindanao are not black Sigatoka caused by *P. fijiensis* but are mainly leaf blight caused by six species belonging to the genus *Nigrospora*. As important information for disease control, it was considered that fungal senescence may occur due to fungicide treatments, as some of these are sensitive to the 14 fungicides used for black Sigatoka disease, while others are not. It is recommended that further surveys of fungal diseases throughout Mindanao Island be conducted and that fungicide selection for disease control be based on a comprehensive understanding of banana leaf diseases.

## Materials and methods

### Isolation of fungi

In July 2024, 175 of banana leaves (27 to 30 leaves from each banana plantation) with Sigatoka-like symptoms were collected at 6 Cavendish banana plantations located in “Barangay Dabong-dabong, Valencia City, Bukidnon (7°58′28″N 125°4′55″E; altitude: 313 m)” (29 leaves), “Barangay Little Panay, Panabo, Davao del Norte (7°17′44″N 125°38′46″E; altitude: 17 m)” (30 leaves), “Barangay Panombon, Mati, Davao Oriental (7°0′32″N 126°16′28″E; altitude: 20 m)” (30 leaves), “Barangay Patag-Alanib, Lantapan, Bukidnon (8°1′5″N 125°1′57″E; altitude: 851 m)” (27 leaves), “Barangay Tugbok, Davao City, Davao del Sur (7°11′45″N 125°27′23″E; altitude: 217 m)” (29 leaves), and “Barangay Ula, Davao City, Davao del Sur (7°8′16″N 125°29′37″E; altitude: 291 m) (30 leaves)” in Mindanao Island, Philippines (Fig. 13). Banana plants selected for sampling were estimated to be between five and eight months of age. Symptoms included small spots with gray centers and yellow halos, as well as spots that turned brown or dark brown as they expanded. The leaves also showed signs of coalescing lesions as the spots expanded. Fungi were isolated from the symptoms, including black and brown spots and yellow hallows (Fig. 7). Plant tissues with apparent symptoms were cut into 3-mm square segments, sterilized with 0.6% (v/v) sodium hypochlorite for 1 min, rinsed twice with sterilized water, and placed on water agar plates. To obtain monocultures of fungal isolates, a single hypha was picked directly from the mycelium that emerged from the tissues on water agar. Colonies with different appearances in a single sample were individually isolated, and then transplanted onto potato dextrose agar (PDA) plates. The monoculture isolates were deposited in the culture collection at Tamagawa University and maintained on PDA plates.

**Fig. 13.**
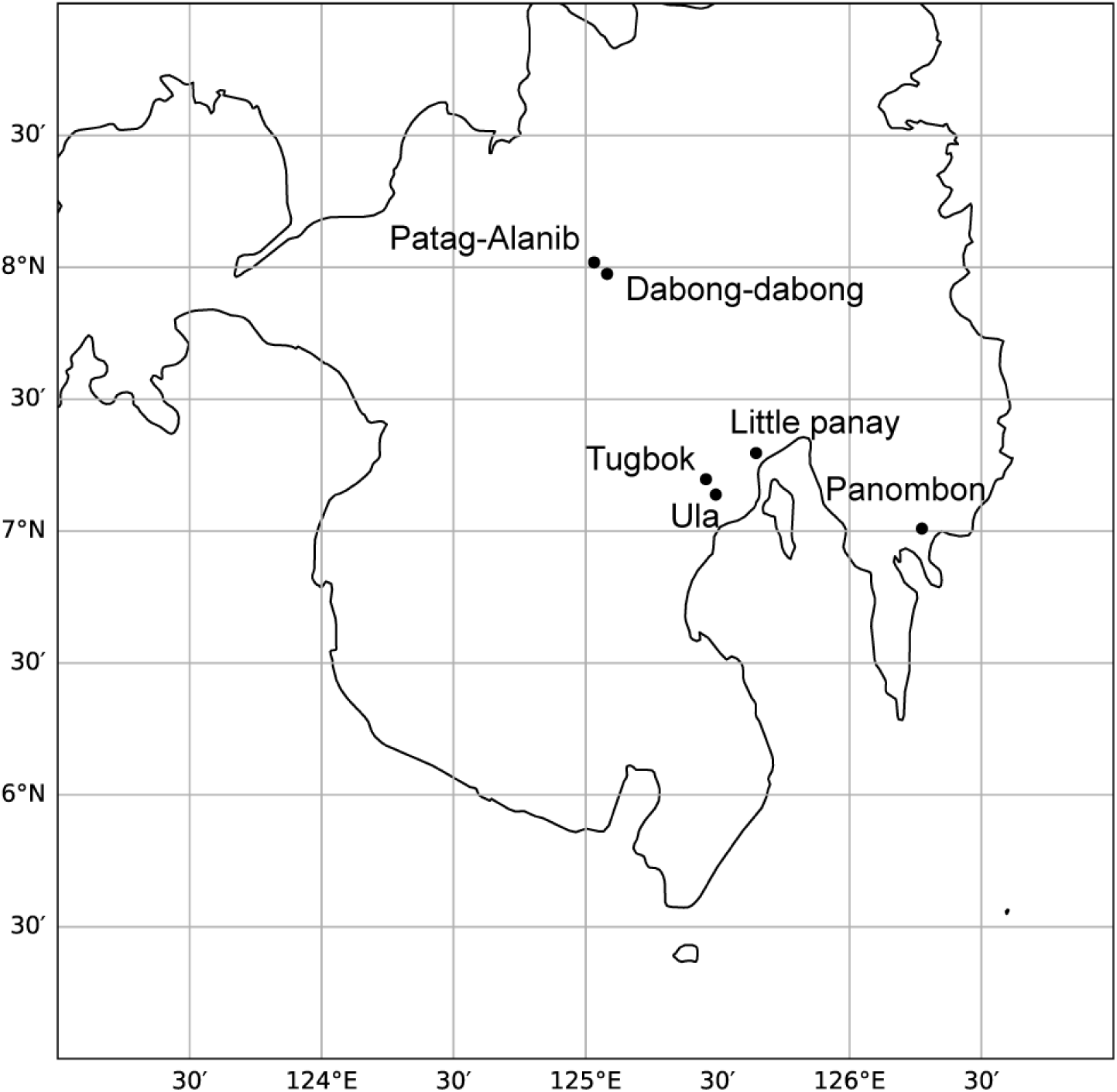
Sampling sites in the Mindanao island, Philippines.

### Polymerase chain reaction (PCR)

Template DNA for PCR was obtained from the mycelia of each isolate grown on potato dextrose agar for 7 d using a CTAB method [20]. PCR was performed using a 10 μL PCR mixture containing 7 μL distilled water, 1 μL 10× Ex Taq buffer with MgCl_2_, 0.8 μL dNTP (10 mM each), 0.1 μL of each primer (50 μM), 0.05 μL Ex Taq DNA polymerase (5 U μL^−1^, Takara, Tokyo, Japan), and 1.0 μL of DNA template. For DNA sequencing of all 160 isolates, the partial 18S rRNA gene, ITS1, 5.8S rRNA gene, ITS2, and partial 28S rRNA gene internal transcribed spacer (ITS) regions were amplified using primer pairs ITS4 and ITS5 [21]. For 107 isolates of *Nigrospora* fungi, in addition to the ITS region, two DNA regions, partial sequences of the *β-tubulin* and the *transcribed elongation factor 1 alfa* (*tef1*) genes, were amplified by using primer pairs Bt2a and Bt2b [22] and Nigro_EF1 (5’-AAC ATC GTC GTT ATC GGC CA-3’) and Nigro_EF2 (5’-TGA TCT CCT TGT AAC GGG CC-3’) (designed in this study), respectively. As the commonly used primers EF1 and EF2 failed to amplify the tef1 region from the isolates obtained in this study, Nigro_EF1 and Nigro_EF2 was newly designed in this study (Supplementary Figure 9).

The PCR was performed under the conditions listed in Table 3. PCR reaction primers were used for sequencing. PCR products were sequenced by the FASMAC Co. Ltd. in Japan (https://fasmac.co.jp/). Obtained forward and reverse sequence reads were assembled using BioEdit v7.7.1.0 software [23] (https://thalljiscience.github.io/page2.html).

**Table 3.**
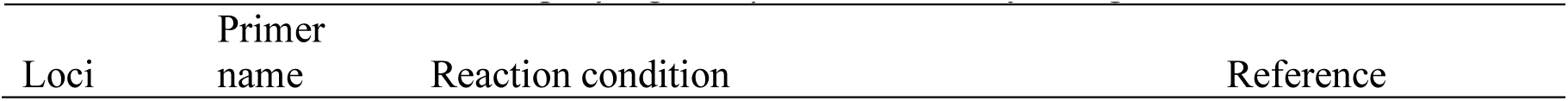
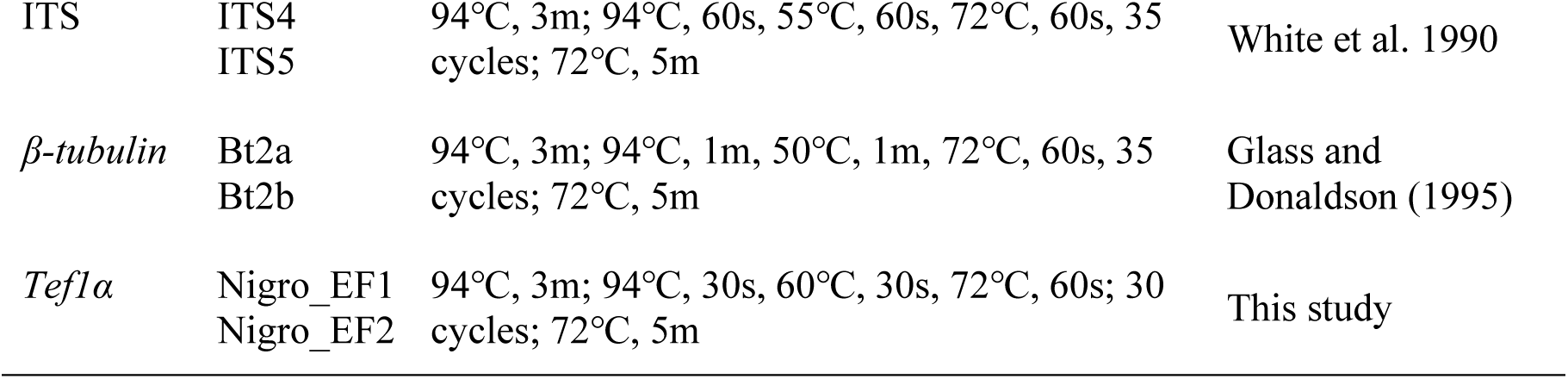
PCR conditions for amplifying ITS, *β-tubulin*, and *tef1α* regions.

**Table 4.**
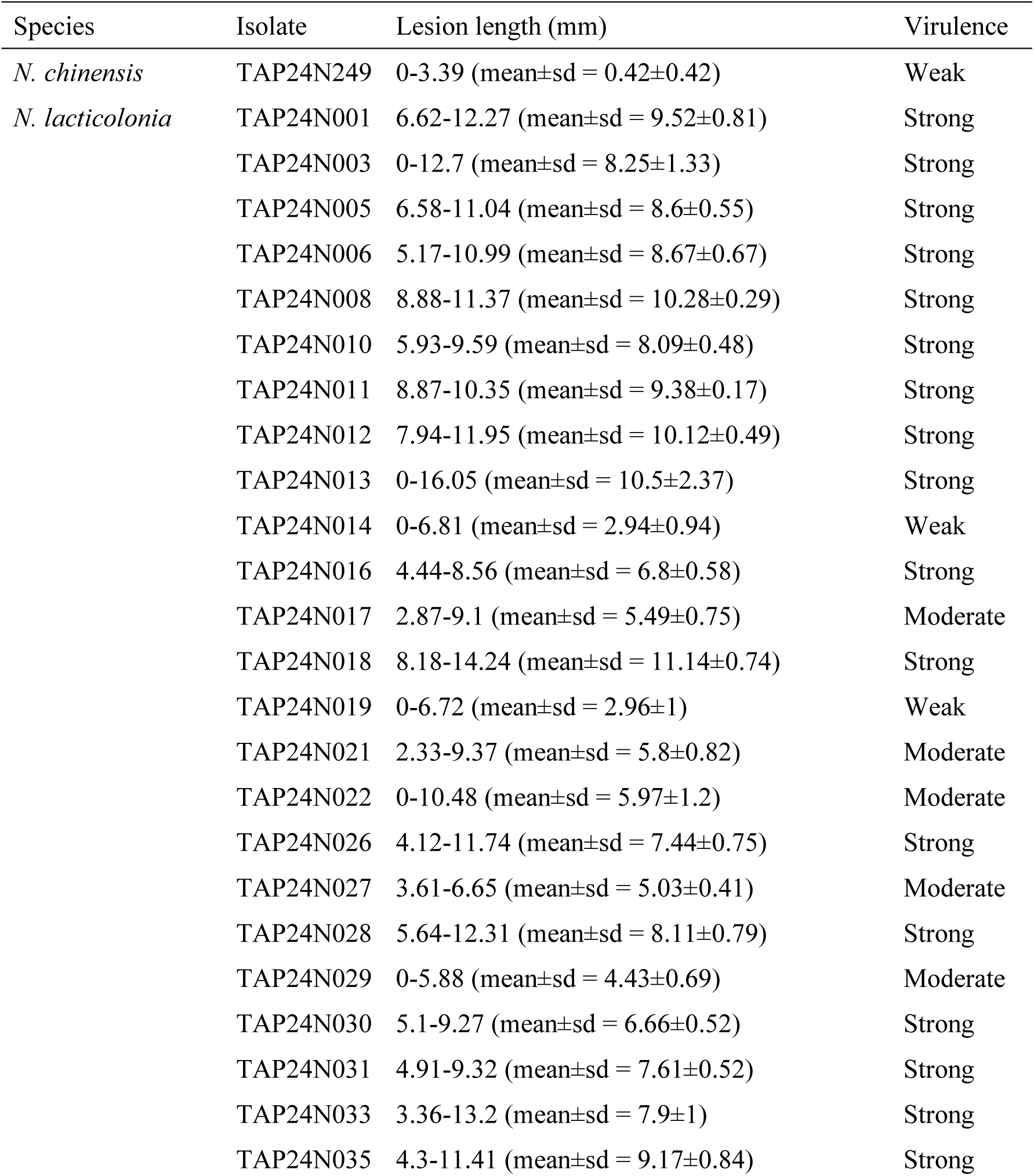
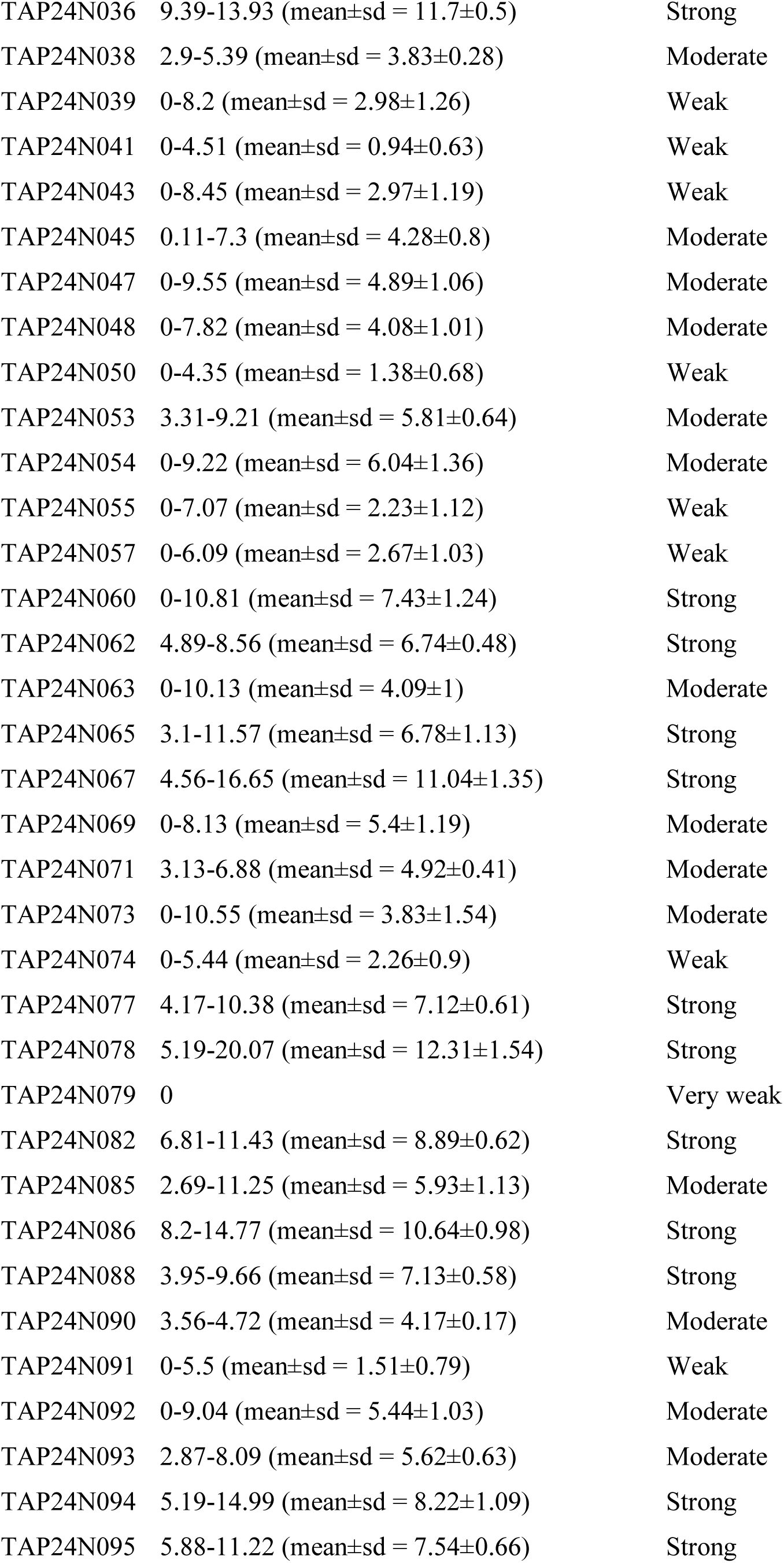
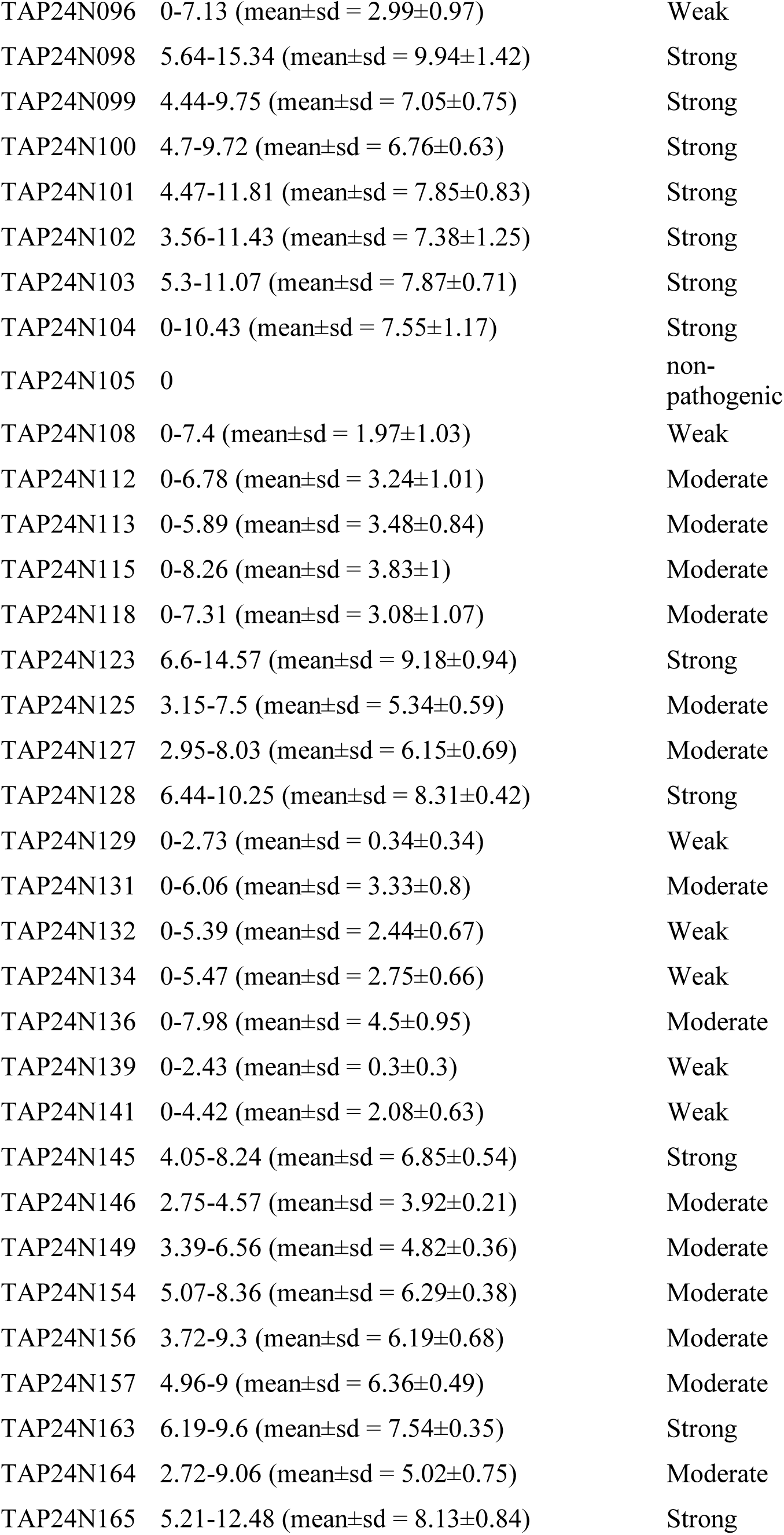
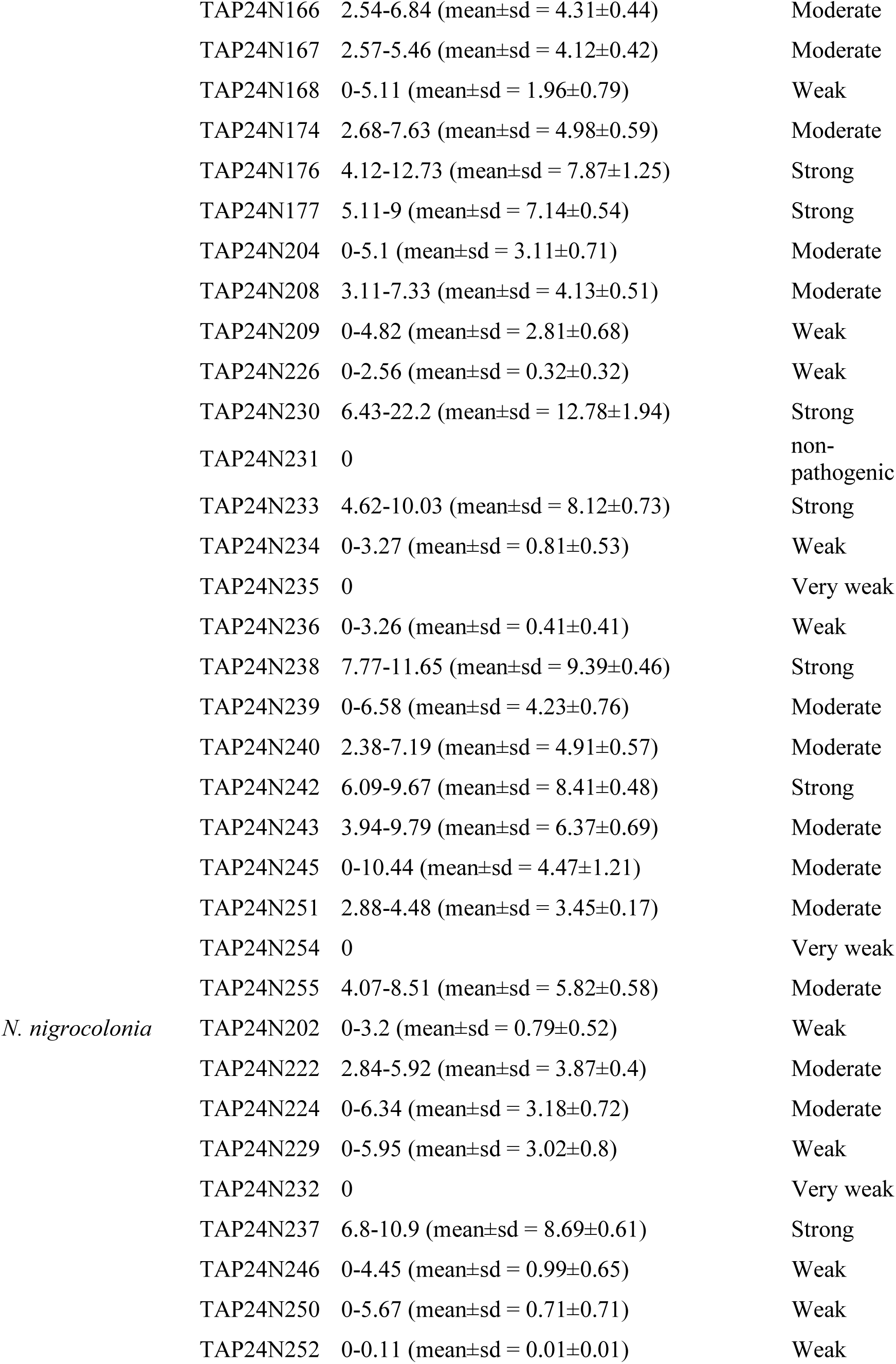
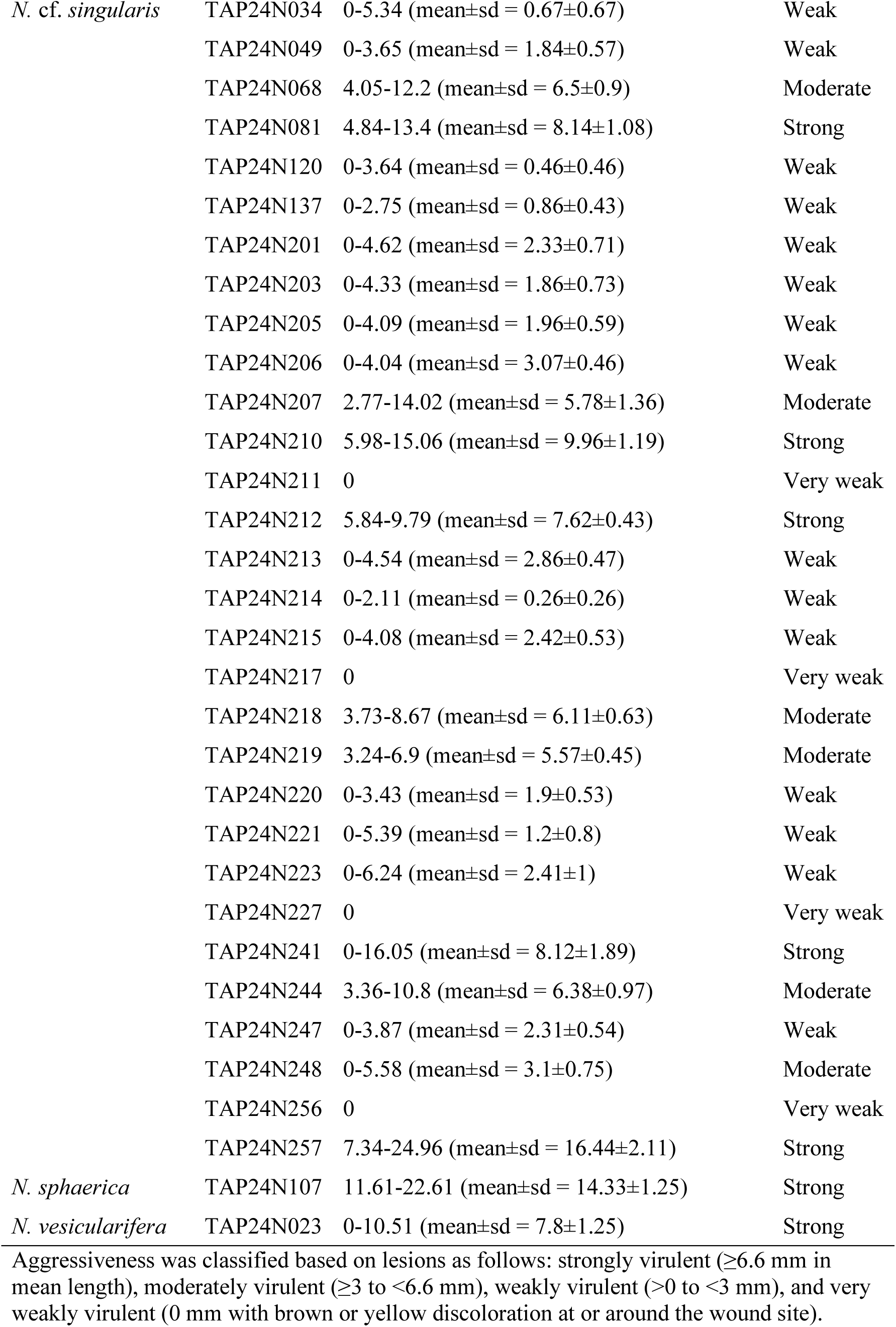
Lesion length and virulence of isolates based on pathogenicity tests.

### Molecular phylogenetic analysis

To identify the 217 isolates at the genus level, we used ITS region sequences to conduct a BLAST search in the NCBI database. Based on the results of the BLAST search, we selected 160 isolates belonging to the genus *Nigrospora* for further analyses to determine the dominant species and diversity of pathogens in the genus *Nigrospora*. To construct phylogenetic trees, 160 isolates and 36 strains of *Nigrospora* spp., including ex-type strains, were used as ingroup taxa (Table 2). As an out group, *Apiospora malaysiana* and *A. pseudoparenchymatica* were selected from the genus *Apiospora*, which is a sister group to the genus *Nigrospora*. All of the ITS, *β-tubulin*, and *tef1* sequences were used for multiple alignments by ClustalW with BioEdit [23] and concatenated into one dataset (Supplementary data 1). Bayesian Inferences (BI) with MrBayes 3.1.2 [24] (https://nbisweden.github.io/MrBayes/index.html) implementing the Markov chain Monto Carlo technique were performed based on the combined dataset of ITS, *β-tubulin*, and *tef1α*. The best-fitted model for each locus for BI was selected with MrMODELTEST2.3 [25]. Each locus was treated as a separate partition. The parameters in MrBayes were set as follows: lset nst = 6, 2, and 6 for ITS, β-tubulin, and *tef1α*, respectively; rates = invgamma, prset statefreqpr = dirichlet (1,1,1,1), nchains = 4, temp = 0.2, swapfreq = 1, and nswaps = 1. The Markov chains were continued until average standard deviation of split frequencies fell below 0.0095 (7,305,000 generations). Only trees sampled after the average standard deviation of split frequencies dropped below 0.01 were used for consensus tree construction. Accordingly, the first 7,244,000 generations were discarded as burn-in. The remaining trees were used to construct a 50% majority-rule consensus tree. Branches that received Bayesian posterior probabilities (BPP) greater than or equal to 0.95 were considered as significantly supported. Bayesian analysis was also performed for each locus with the model described above, and using same parameters. For BI based on ITS and *β-tubulin* sequences, the Markov chains were continued until average standard deviation of split frequencies fell below 0.0095 (27,815,000 and 1,835,000 generations, respectively). Only trees sampled after the average standard deviation of split frequencies dropped below 0.01 were used for consensus tree construction. Accordingly, the first 28,761,000 for ITS and 1,760,000 for *β-tubulin* generations were discarded as burn-in. For BI based on *tef1α*, despite running the analysis for 40,000,000 generations, the average standard deviation of split frequencies did not fall below 0.01. Therefore, we constructed the phylogenetic tree using 1,000 generations of data starting from generation 28,761,000, which yielded the lowest average standard deviation of split frequencies value (0.016862).

Phylogenetic analyses with maximum likelihood method were performed using Molecular Evolutionary Genetics Analysis (MEGA10) software (Kumar 2018) [26]. Tamura-Nei model was selected as the best substitution model based on the Akaike information criterion using MEGA 10. All sites with gaps were treated as missing data. The reliability of each internal branch in the phylogenetic trees was assessed using bootstrap analysis with 1,000 random-addition replicates [27].

### Morphological observation

Our isolates were cultured on synthetic low-nutrient agar (SNA) [28] at 28℃ for 10 days to induce sporulation in the dark. The shape and size of the microscopic structures were observed using light microscopy (B51, Olympus, Tokyo, Japan), and the colonies were assessed. At least 30 conidiogenous cells and conidia were measured to determine mean colony size. Three SNA plates were examined, and from each plate, ten or more conidiophores and conidia were observed for measurement. These measurements were conducted using ImageJ 1.52a software (Rasband, WS, ImageJ; US National Institutes of Health: https://imagej.net/ij/index.html).

### Pathogenicity test

Each of our 160 isolates was tested for pathogenicity using mycelial discs (5 mm diameter) obtained from the edges of colonies grown on PDA media for three days. These discs were then applied to four sites on non-wounded and wounded segments of healthy banana leaves (approx. 5×5 cm in size; cv. Dwarf Cavendish). For each fungal strain, two non-wounded and two wounded leaf segments were used. Three puncture sites were created on the wounded leaves using a heated sewing needle. Each puncture site was spaced approximately 2 mm apart. The inoculated leaves were covered with plastic boxes to maintain high humidity (85–90%) with wet paper towel and maintained at approximately 25 °C under natural light condition for three days. PDA was used as a control. Pathogenicity was evaluated based on the size and coloration of lesions observed at 4 days after inoculation (dpi). Isolates that did not produce elliptical necrotic lesions but caused brown or yellow discoloration at or around the wound site were considered extremely weak in pathogenicity and were classified as “very weakly virulent”. For isolates that formed brown, spindle-shaped necrotic lesions, the lesion length at each of the eight inoculation points was measured, and the mean lesion length was used as an index of pathogenicity. To further categorize the degree of pathogenicity, the 33rd and 66th percentiles of the mean lesion lengths were calculated. Isolates with mean values below the 33rd percentile were classified as “weakly virulent”, those between the 33rd and 66th percentiles as “moderate virulent”, and those above the 66th percentile as “strongly virulent”. This classification was performed using R (version 4.2.2), with data processing conducted via the dplyr package. Values of zero were excluded prior to quantile calculation. Based on these criteria, all isolates were ultimately categorized into five levels of pathogenicity: “non-pathogenic”, “very weakly virulent”, “weakly virulent”, “moderate virulent”, and “strongly virulent”. At 7 dpi, the symptoms appeared, the fungi were re-isolated from the diseased tissue using the same method as that of the description in “isolation of fungi” and re-identified based on morphologies of colony on PDA.

### Haplotype analyses

The haplotypes of *N. lacticolonia*, *N. singuralis*, and *N. nigrocolonia* were analyzed to reveal the spread of pathogens on Mindanao Island. The sequence of the *tef1α* gene was used for the analyses because mutations in each species were observed in the gene. The *tef1α* gene sequences for each species were aligned using MEGA 10, and sites containing gaps at both ends were trimmed. The total number of variable sites (S) and the nucleotide diversity (Pi) were calculated using MEGA 10. The trimmed alignment data were exported to PopArt v1.7 [29] to construct a haplotype network. The geographic distribution of the haplotypes was visualized using the median-joining option in the PopArt program.

### In vitro Fungicide Sensitivity test

In vitro tests were conducted to evaluate the inhibitory effects of the active ingredients in fungicides, currently used in the Philippines, on the mycelial growth of *Nigrospora* spp. Mycelial growth inhibition was used as an indicator of sensitivity. The active ingredients of 14 fungicides belonging to 6 classes of the Fungicide Resistance Action Committee (FRAC) were dissolved in acetone or sterile distilled water and adjusted to a concentration of 1/5 in PDA medium. Aryl hydroxamic acid (SHAM; Fluorochem Ltd.) was used in conjunction with the QoI fungicide pyraclostrobin to inhibit alternative oxidase (AOX), which does not react with cyano groups [30,31]. The final concentrations of the fungicides in the medium were as follows: 100 ppm for bitertanol, difenoconazole, epoxiconazole, flutriafol, tebuconazole, fenpropimorph, spiroxamine, boscalid, and flusapirox; and 200 ppm for mancozeb and chlorothalonil. Media without fungicides (0 ppm) were used as control to formulate inhibition rate. These fungicide concentrations were determined based on the following reasons. In FRAC monitoring assays, the upper concentration range typically falls between 1 and 30 ppm; however, concentrations as high as 100 ppm are occasionally employed. By intentionally setting the concentration higher than the standard FRAC monitoring range, we aimed to detect isolates with clearly low sensitivity. Isolates were classified as sensitive or low-sensitive based on their growth response at 100 ppm or 200 ppm. Those exhibiting visible growth at these concentrations were designated as having low sensitivity.

Mycelial disks (6 mm in diameter) were obtained from the periphery of pre-cultured colonies, which were cultured at 25°C for 3 days in 1/5 PDA. These disks were transferred to 1/5 PDA containing the chemicals at avobe mentioned concentrations. Following a 24-h incubation period at 25°C, the diameters of the colonies were measured, and 6 mm was subtracted from the diameter of the mycelial disc. The inhibition rate was defined as the rate of growth when the control was 100%, and was calculated from the values of the four colonies.

### Principal component analysis (PCA)

The PCA of standardized variables was performed using R (v4.2.2) to determine whether there was a difference in the inhibition rate among species/haplotypes and sampling sites. We conducted PCA using the inhibition rate data obtained from the 14 fungicides as variants. The final principal components (PCs) were determined by excluding the data of individuals that fell outside the ±3σ range as outliers. Euclidean distances were calculated based on the obtained coordinates of PC1 and PC2, and a dendrogram was constructed using Ward’s method. Outliers were defined based on principal component scores obtained from PCA, using a threshold of three standard deviations from the mean for PC1 and PC2. This criterion is grounded in the statistical property of the normal distribution, where approximately 99.7% of data points fall within ±3 standard deviations. Values exceeding this range are considered statistically rare and thus indicative of potential outliers. A sample was classified as an outlier if either its PC1 or PC2 score exceeded the threshold.

## Acknowledgments

We are grateful to Ms Sayuri Sakurai (Tamagawa University BacaDM Project) for her assistance in preparing the culture medium for this research.

## Data Availability

The datasets generated during the current study are available in the GenBank provided by NCBI/DDBJ. The GenBank accession numbers are listed in Table 2.

## Author Contribution

S.N. conceived the study, designed the methodology, collected field samples, conducted fungicide sensitivity tests, conducted all analyses, visualization of the data, and led the manuscript preparation. Y.H. and Y.T. conducted fungicide sensitivity tests. K.U. designed the methodology for fungicide sensitivity tests. M.A.M., R.V., S.M.C., and A.F.P. collected field samples. K.W. supervised this study, acquired funding, conducted fungicide sensitivity tests, and assisted in manuscript editing and project coordination. All authors reviewed and approved the final manuscript

## References

1. Nozawa, S. et al. *Fusarium mindanaoense* sp. nov., a new fusarium wilt pathogen of cavendish banana from the Philippines belonging to the F. fujikuroi species complex. J. Fungi (Basel) 9, 443; 10.3390/jof9040443, PubMed: 37108898 (2023).

2. Ploetz, R. C. Fusarium wilt of banana. Phytopathology 105, 1512–1521; 10.1094/PHYTO-04-15-0101-RVW, PubMed: 26057187 (2015).

3. Wang, B. et al. A new leaf blight disease caused by *Alternaria jacinthicola* on banana in China. Horticulturae 8, 12 (2021).

4. Fu, B. Z. et al. First report of leaf spot caused by *Alternaria alternata* on Chinese dwarf banana in China. Plant Dis. 98, 691–691; 10.1094/PDIS-08-13-0831-PDN, PubMed: 30708541 (2014).

5. Huang, R. et al. Identification and characterization of Colletotrichum species associated with anthracnose disease of banana. Plant Pathol. 70, 1827–1837; 10.1111/ppa.13426 (2021).

6. Crous, P. W. & Mourichon, X. Mycosphaerella eumusae and its anamorph *Pseudocercospora eumusae* spp. nov: causal agent of eumusae leaf spot disease of banana. Sydowia 54, 23–34 (2002).

7. Sadid, K. N. E., Hossain, M. A., Munshi, A. R. & Karim, M. R. First detection, isolation and molecular characterization of banana leaf spot diseases caused by *Nigrospora sphaerica* and *Neocordana musae* in Bangladesh. Arch. Phytopathol. Plant Prot. 56, 1112–1125 (2023).

8. Fortune, M. P. et al. First report of black Sigatoka disease (causal agent *Mycosphaerella fijiensis*) from Trinidad. Plant Pathol. 54, 246; 10.1111/j.1365-3059.2005.01123.x (2005).

9. Strobl, E. & Mohan, P. Climate and the global spread and impact of bananas’ black leaf sigatoka disease. Atmosphere 11, 947; 10.3390/atmos11090947 (2020).

10. Esguera, J. G., Balendres, M. A. & Paguntalan, D. P. Overview of the Sigatoka leaf spot complex in banana and its current management. Trop. Plant 3, e002; 10.48130/tp-0024-0001 (2024).

11. Rhodes, P. L. A new banana disease in Fiji. CPN 10, 38–41 (1964).

12. Pordel, A., Dehghani, K., Amirmijani, A. & Miri, K. Identification and pathogenicity of Fusarium sensu lato species causeing wilting and leaf spot of Banana in South of Iran. Iran. J. Plant Pathol. 58, 96–108 (2022).

13. Raza, M. et al. Culturable plant pathogenic fungi associated with sugarcane in southern China. Fungal Divers. 99, 1–104; 10.1007/s13225-019-00434-5 (2019).

14. Wang, M et al. Phylogenetic reassessment of *Nigrospora*: Ubiquitous endophytes, plant and human pathogens. Persoonia 39, 118–142; 10.3767/persoonia.2017.39.06 (2017).

15. Karlsson, I., Friberg, H., Steinberg, C. & Persson, P. Fungicide effects on fungal community composition in the wheat phyllosphere. PLOS One 9, e111786; 10.1371/journal.pone.0111786, PubMed: 25369054 (2014).

16. Hartmann, F. E. et al. The complex genomic basis of rapid convergent adaptation to pesticides across continents in a fungal plant pathogen. Mol. Ecol. 30, 5390–5405; 10.1111/mec.15737, PubMed: 33211369 (2021).

17. Zhang, Y. Y., et al. *Nigrospora humicola* (Apiosporaceae, Amphisphaeriales), a new fungus from soil in China. Diversity 16, 118 (2024).

18. Meredith, D. S. Atmospheric content of nigrospora spores in Jamaican banana plantations. J. Gen. Microbiol. 26, 343–349; 10.1099/00221287-26-2-343, PubMed: 14472772 (1961).

19. Hirst, J. M. Changes in atmospheric spore content: diurnal periodicity and the effects of weather. Trans Br Mycol Soc. 36, 375–IN8; 10.1016/S0007-1536(53)80034-3 (1953).

20. Doyle, J. J. & Doyle, J. L. A rapid DNA isolation procedure for small quantities of fresh leaf tissue. Phytochem Bull. 19A1 A27, 11–15 (1987).

21. White, T. J., Bruns, T., Lee, S. & Taylor, J. W. Amplification and direct sequencing of fungal ribosomal RNA genes for phylogenetics. in 15 (ed. Gelfand, D. H., Shinsky, J. J. & White, T. J.) PCR Protocols: A Guide to Methods and Applications Ch. 315–322 (M.A., Innis) (Acad. Press (San Diego, 1990)).

22. Glass, N. L. & Donaldson, G. C. Development of primer sets designed for use with the PCR to amplify conserved genes from filamentous Ascomycetes. Appl Environ Microbiol. 61, 1323–1330; 10.1128/aem.61.4.1323-1330.1995, PubMed: 7747954 (1995).

23. Hall, T. A. BioEdit: a user-friendly biological sequence alignment editor and analysis program. for Windows 95/98/NT. Nucleic Acids Symp Ser. 41 95–98, (1999).

24. Ronquist, F. & Huelsenbeck, J. P. MrBayes 3: Bayesian phylogenetic inference under mixed models. Bioinformatics 19, 1572–1574 (2003).

25. Nylander, J.A. Modeltest v2. Program distributed by the author. Evolutionary Biology Centre, Uppsala University (2004).

26. Kumar, S., Stecher, G., Li, M., Knyaz, C., Tamura, K. et al. MEGA X: Molecular evolutionary genetics analysis across computing platforms. Mol. Biol. Evol. 35, 1547–1549; 10.1093/molbev/msy096, PubMed: 29722887 (2018).

27. Felsenstein, J. Confidence limits on phylogenies: an approach using the bootstrap. Evolution 39, 783–791; 10.1111/j.1558-5646.1985.tb00420.x, PubMed: 28561359 (1985).

28. Nirenberg, H. I. A simplified method for identifying Fusarium spp. occurring on wheat. Can. J. Bot. 59, 1599–1609 (1981).

29. Leigh, J. W. & Bryant, D. POPART: full-feature software for haplotype networkconstruction. Methods Ecol. Evol. 6, 1110–1116; 10.1111/2041-210X.12410 (2015).

30. Ishii, H. et al. Characterisation of QoI-resistant field isolates of Botrytis cinerea from citrus and strawberry. Pest Manag. Sci. 65, 916–922; 10.1002/ps.1773, PubMed: 19444805 (2009).

31. Hollomon, D. W., Wood, P. M., Reeve, C. & Miguez, M. Alternative oxidase and its impact on the activity of Qo and Qi site inhibitors. in Modern Fungicides and Antifungal Compounds IV (ed. Dehne, H. W., Gisi, U., Kuck, K. H., Russell, P. E. & Lyr, H.) 31–34 (BCPC, Alton, Hants, UK, 2005).

