## Supplementary material for "Occurrence of *Nigrospora spp.* as the predominant causal agents of leaf spot disease in Cavendish banana in banana plantations in Mindanao Island, Philippines": Supplementary fig 1.pdf

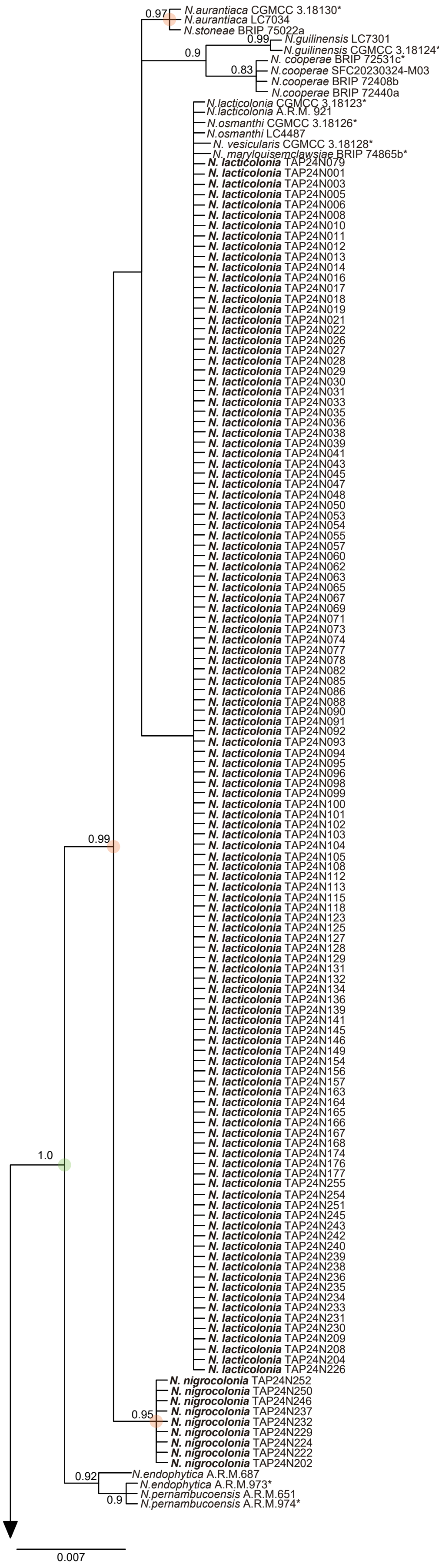


**Supplementary Fig. 1** Phylogenetic trees constructed based on the DNA sequences of ITS region inferred using MrBayes. The numbers on each node are posterior probabilities (PPB) estimated using the software MrBayes. Strain number with asterisk is ex-type strain. Species name and strains with boldface is our isolate. Nodes with a PPB of 1.0 are highlighted with green circles, while those with PPB ≥ 0.95 are indicated by orange circles.

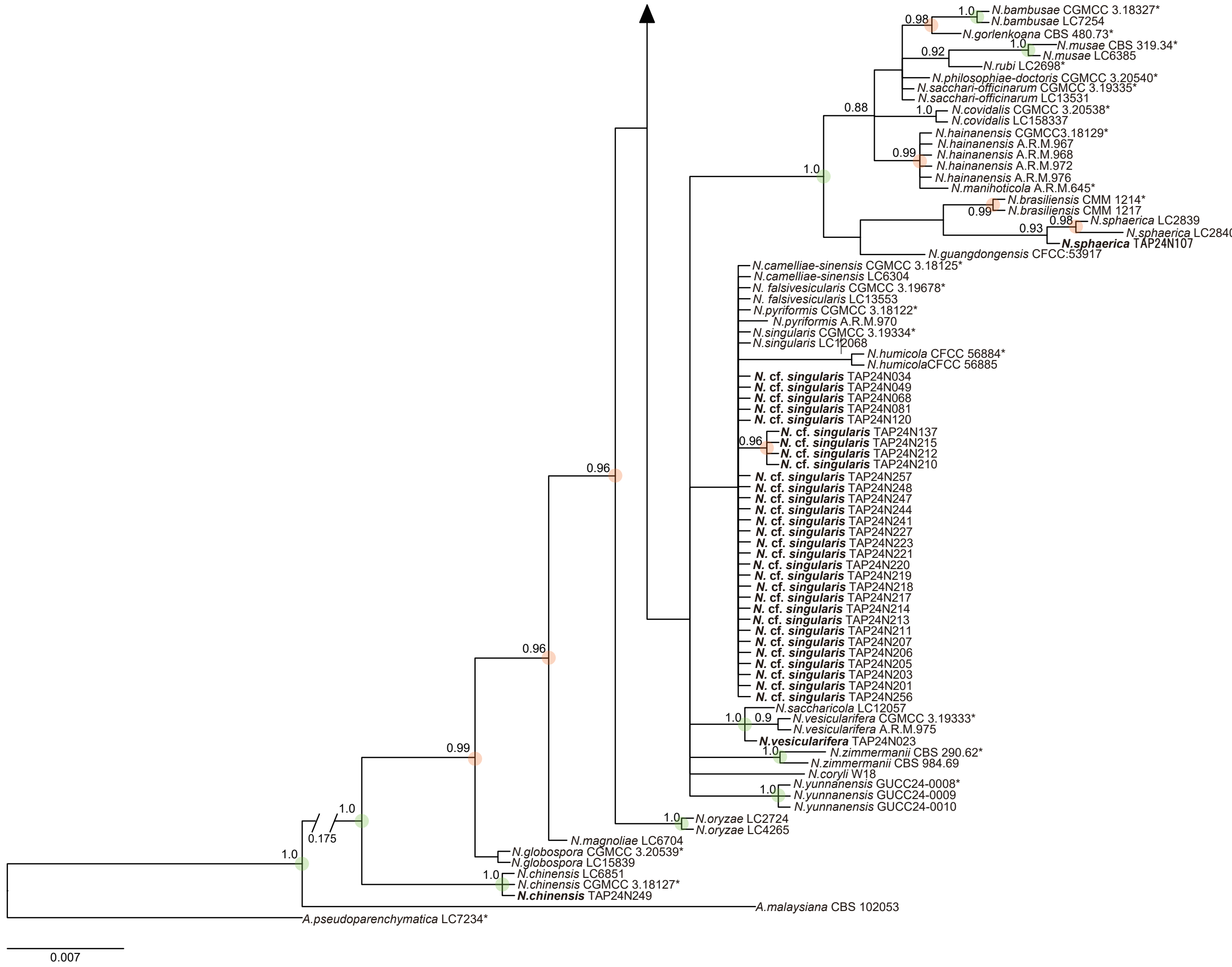


**Supplementary Fig. 1** Continued.
