## Supplementary material for "Occurrence of *Nigrospora spp.* as the predominant causal agents of leaf spot disease in Cavendish banana in banana plantations in Mindanao Island, Philippines": Supplementary fig 3.pdf

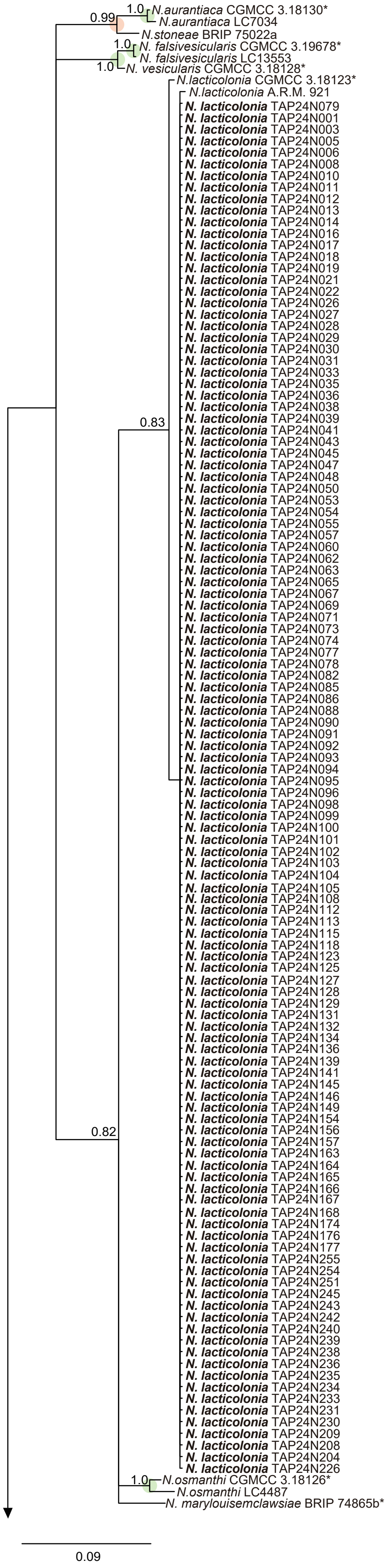


**Supplementary Fig. 3** Phylogenetic tree constructed based on the DNA sequences of *tef1α* inferred using MrBayes. The numbers on each node are posterior probabilities (PPB) estimated using the software MrBayes. Strain number with asterisk is ex-type strain. Species name and strains with boldface is our isolate. Nodes with a PPB of 1.0 are highlighted with green circles, while those with PPB ≥ 0.95 are indicated by orange circles.

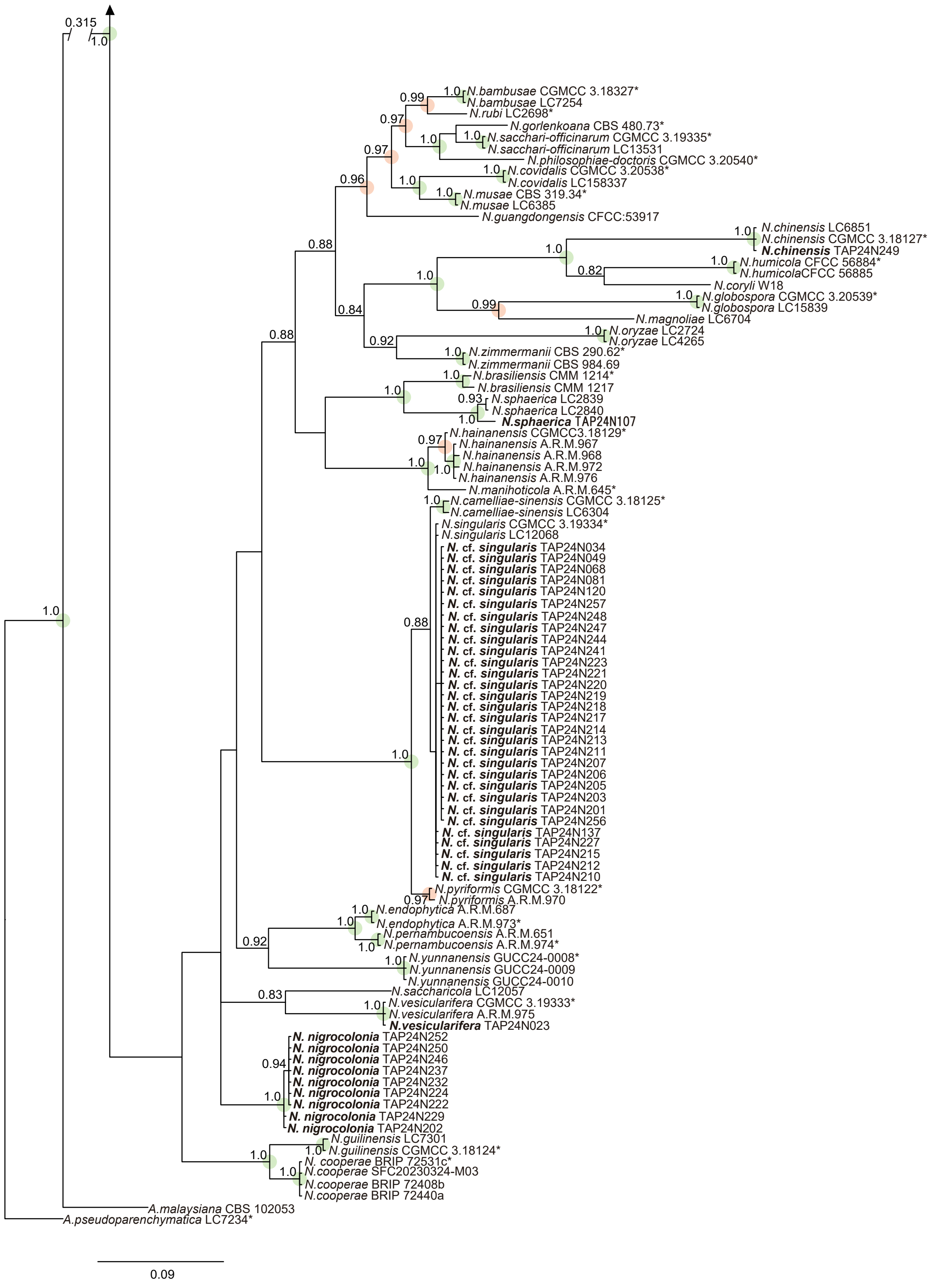


**Supplementary Fig. 3** Continued.
