## Supplementary material for "Occurrence of *Nigrospora spp.* as the predominant causal agents of leaf spot disease in Cavendish banana in banana plantations in Mindanao Island, Philippines": Supplementary fig 4.pdf

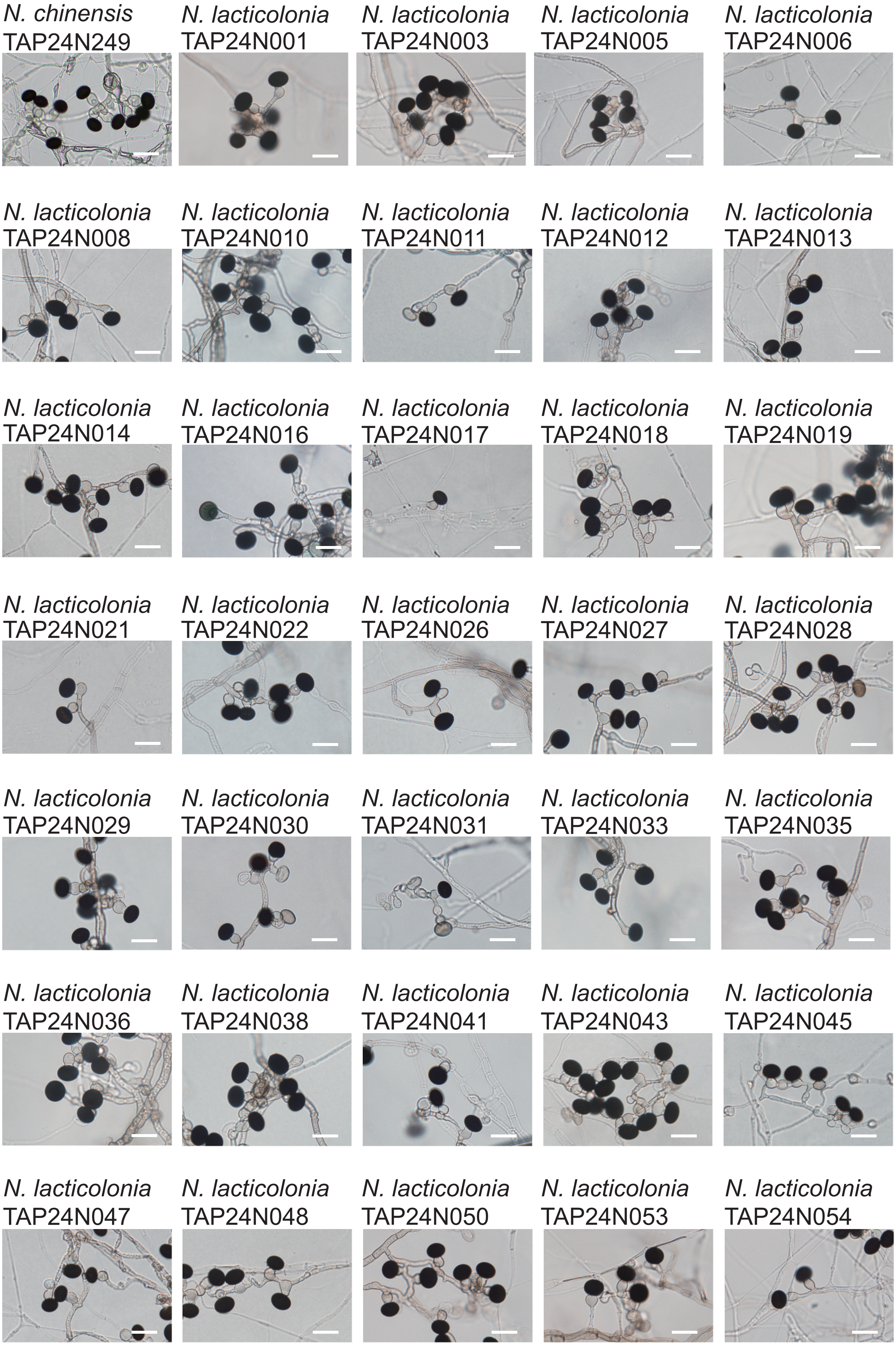


**Supplementary Fig. 4** Conidiogenous cells and conidia of isolates on SNA plates. Scale bars: = 20 μm. No conidial images are available for isolates that did not produce conidia.

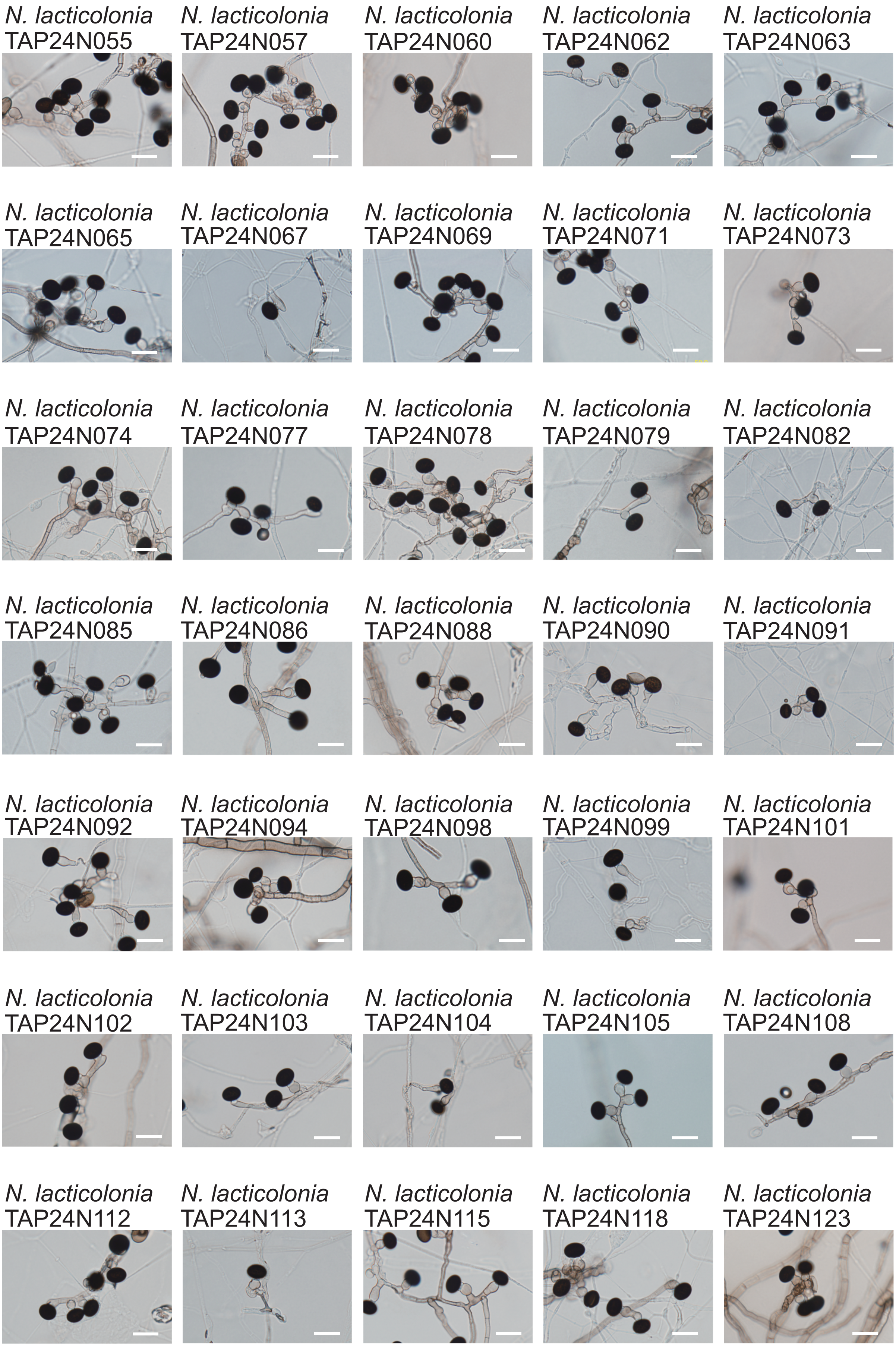


**Supplementary Fig. 4** Continued.

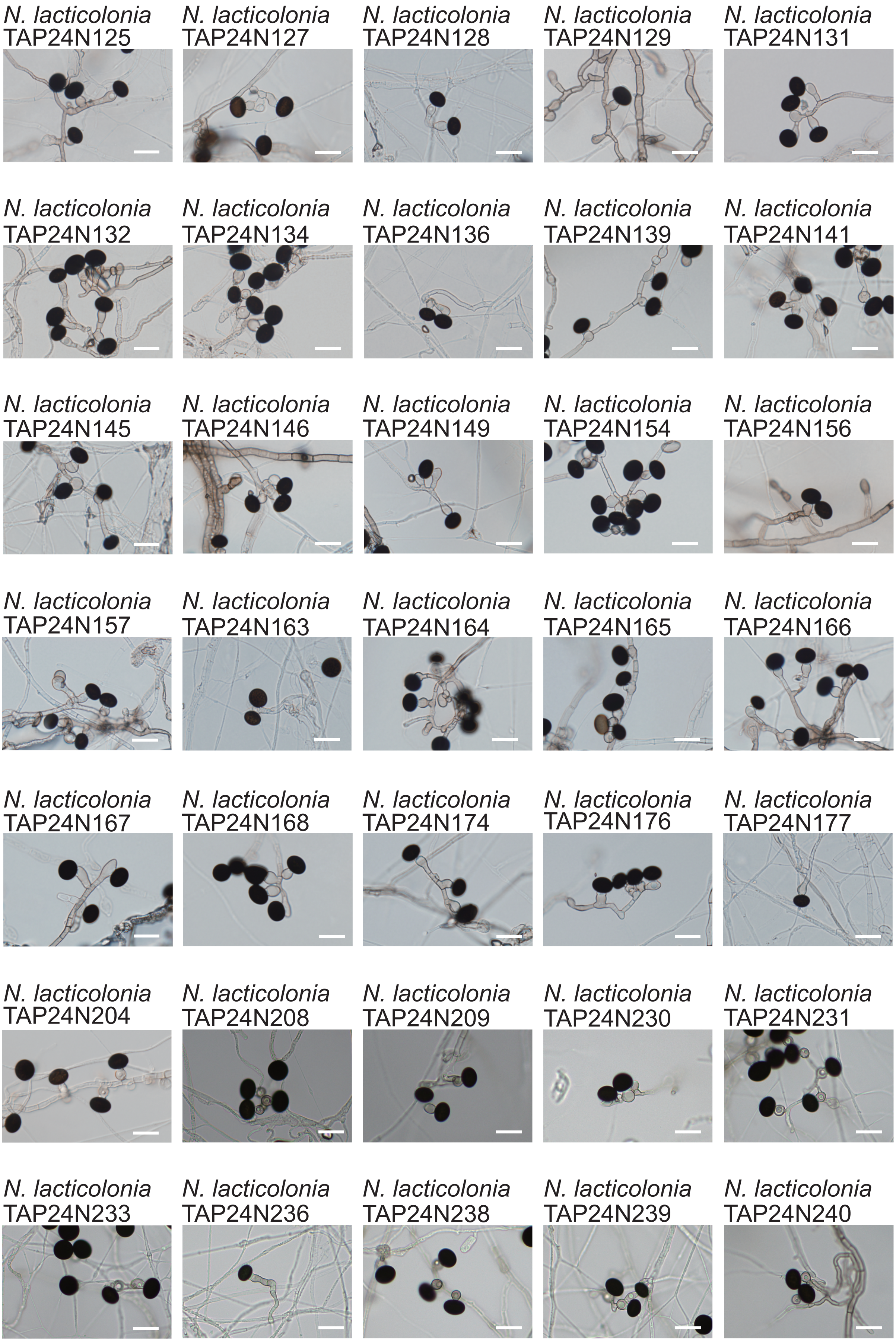


**Supplementary Fig. 4** Continued.

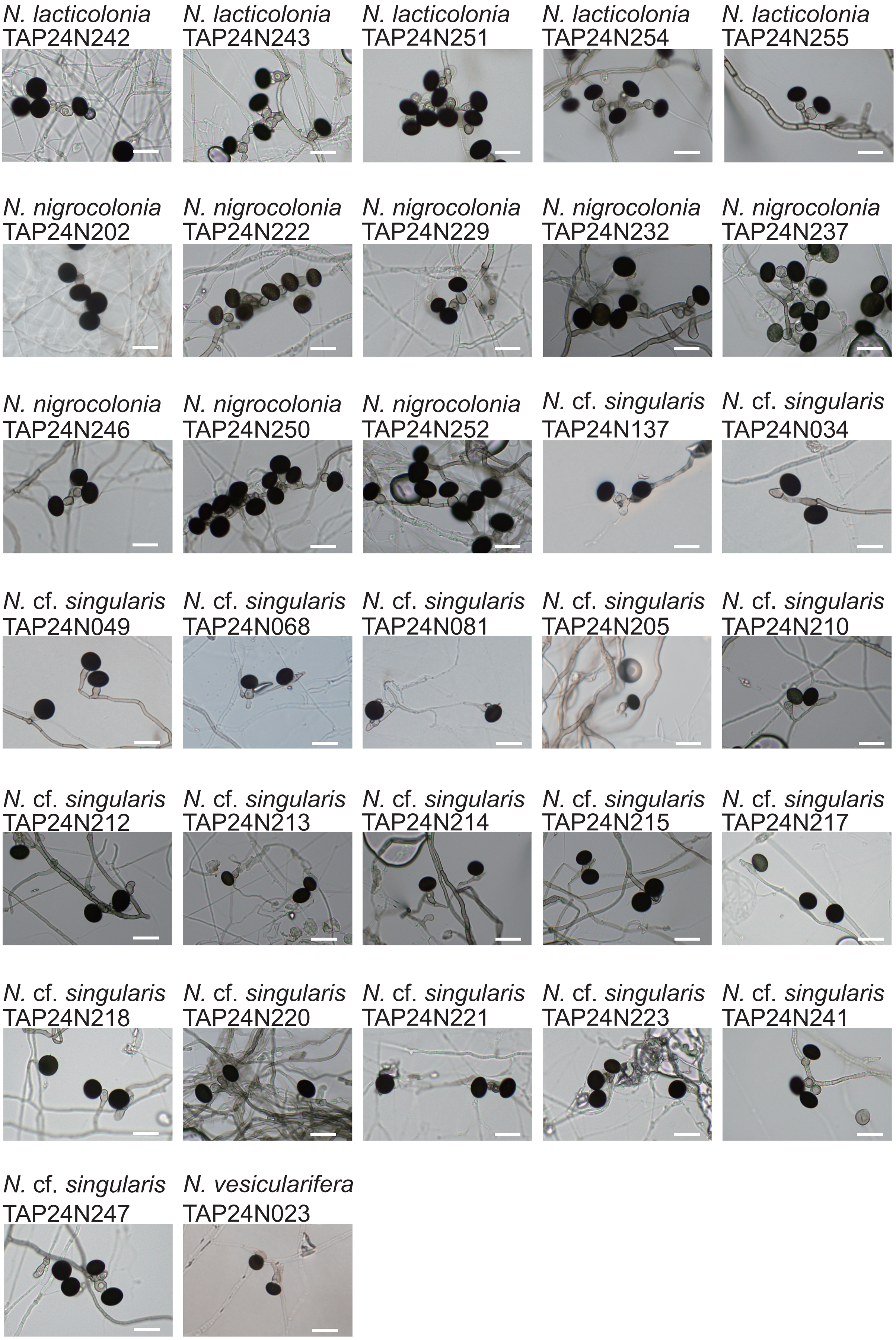


**Supplementary Fig. 4** Continued.
