## Supplementary material for "Occurrence of *Nigrospora spp.* as the predominant causal agents of leaf spot disease in Cavendish banana in banana plantations in Mindanao Island, Philippines": Supplementary fig 5.pdf

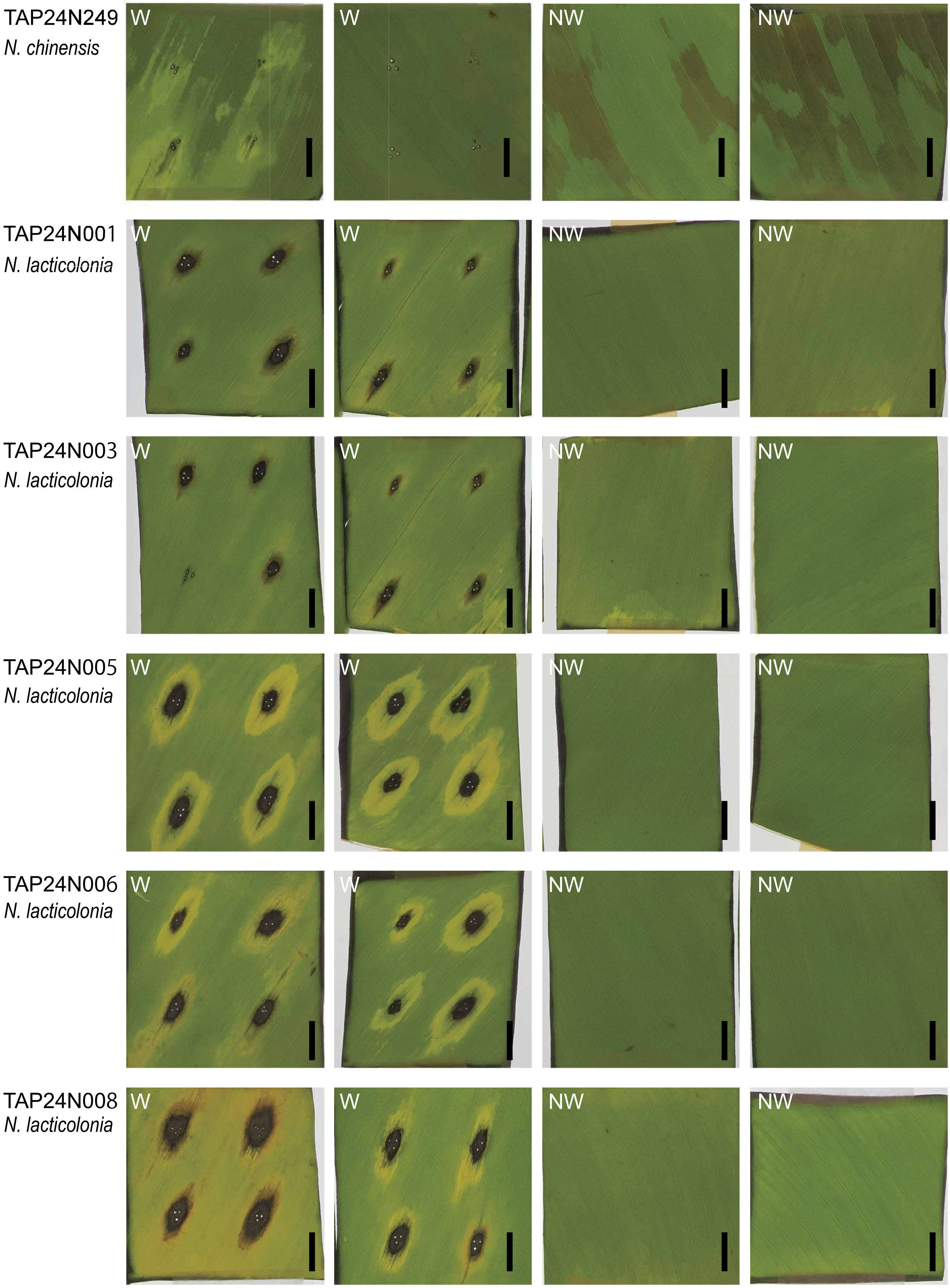


**Supplementary Fig 5** Symptoms on cut leaves of banana (cv. Dwarf Cavendish) 4 days after inoculation with our *Nigrospora* isolates. W: leaves with wounds; NW: leaves without wound. Bars: 1 cm.

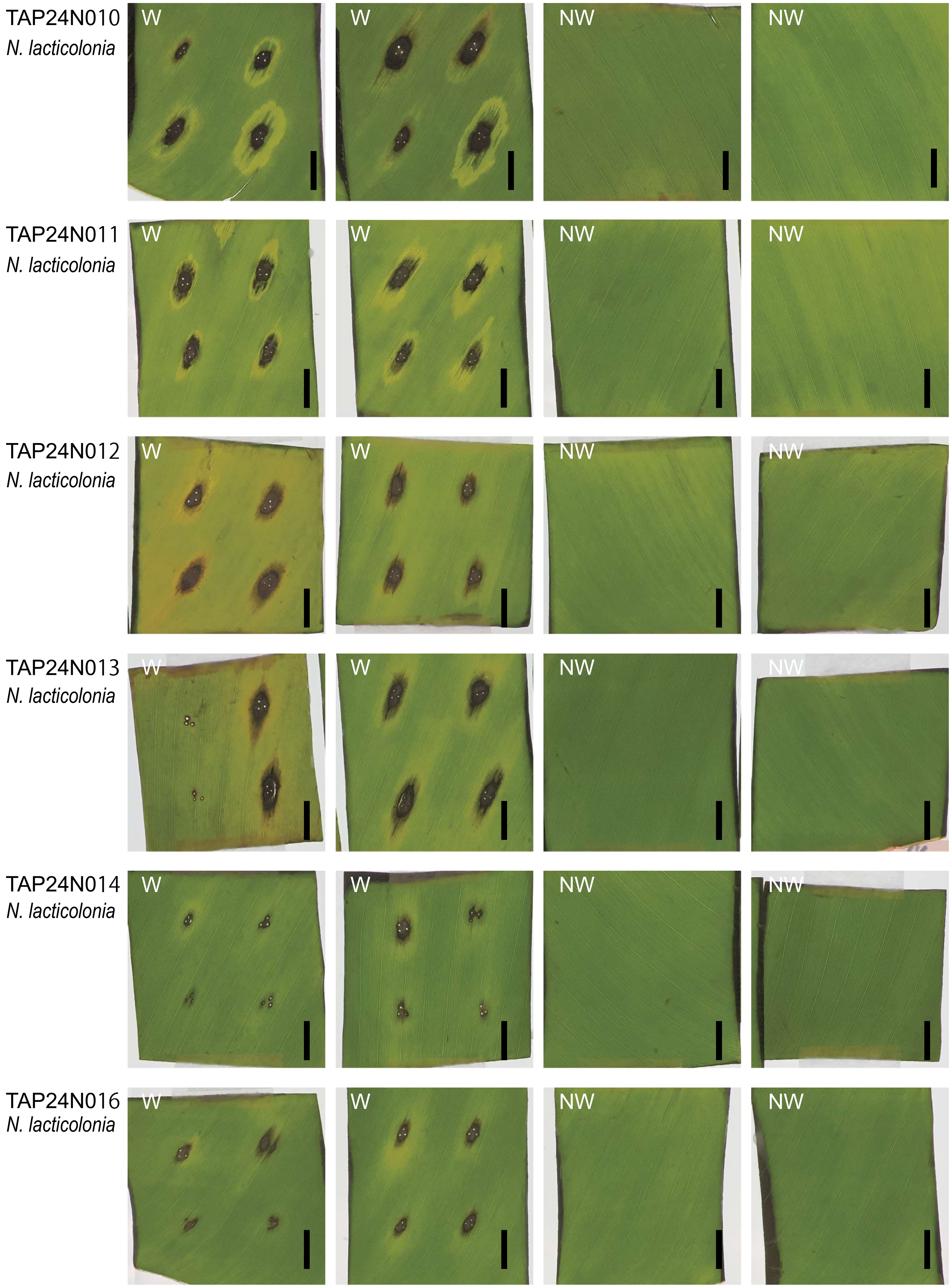


**Supplementary Fig 5** Continued.

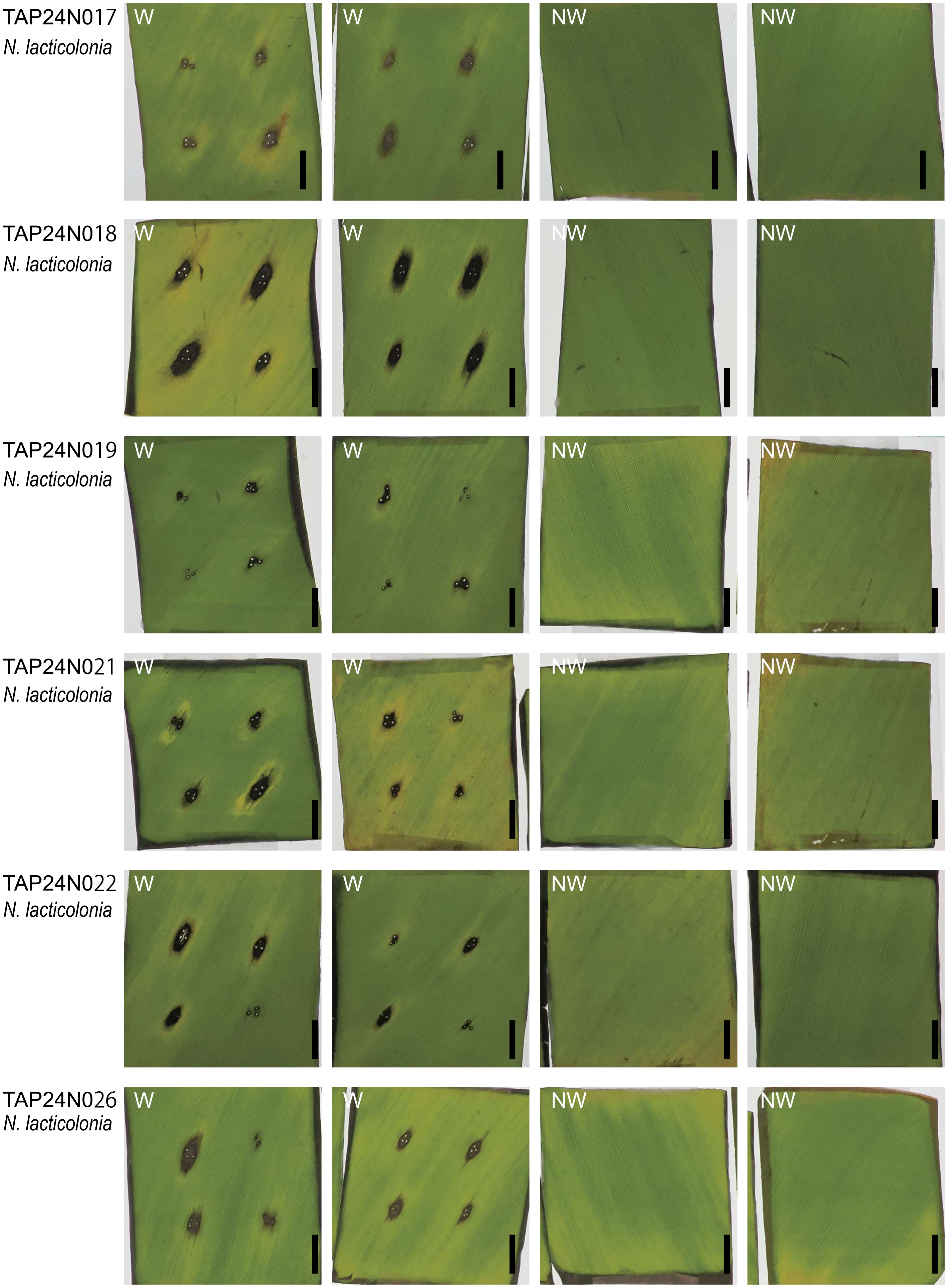


**Supplementary Fig 5** Continued.

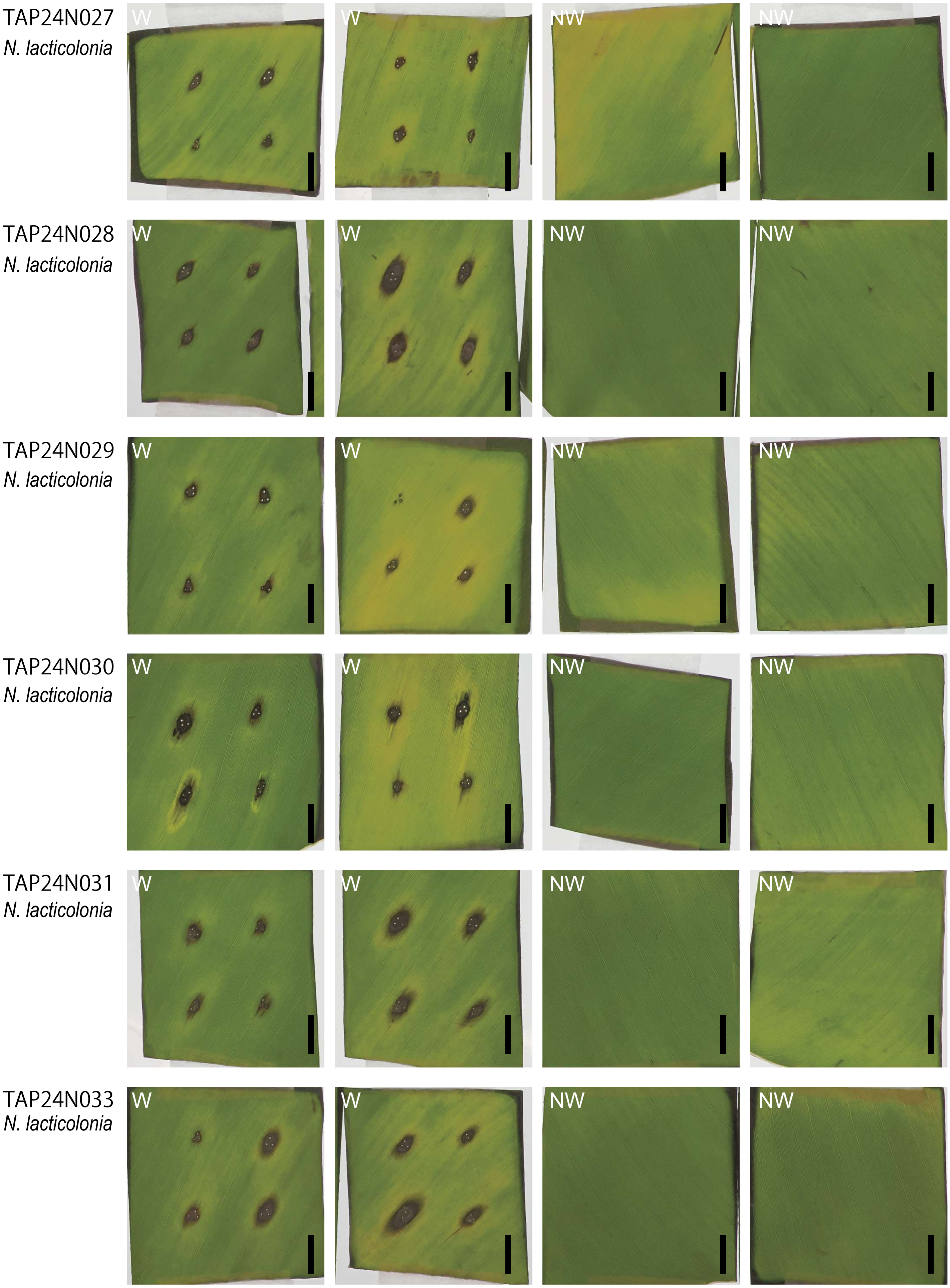


**Supplementary Fig 5** Continued.

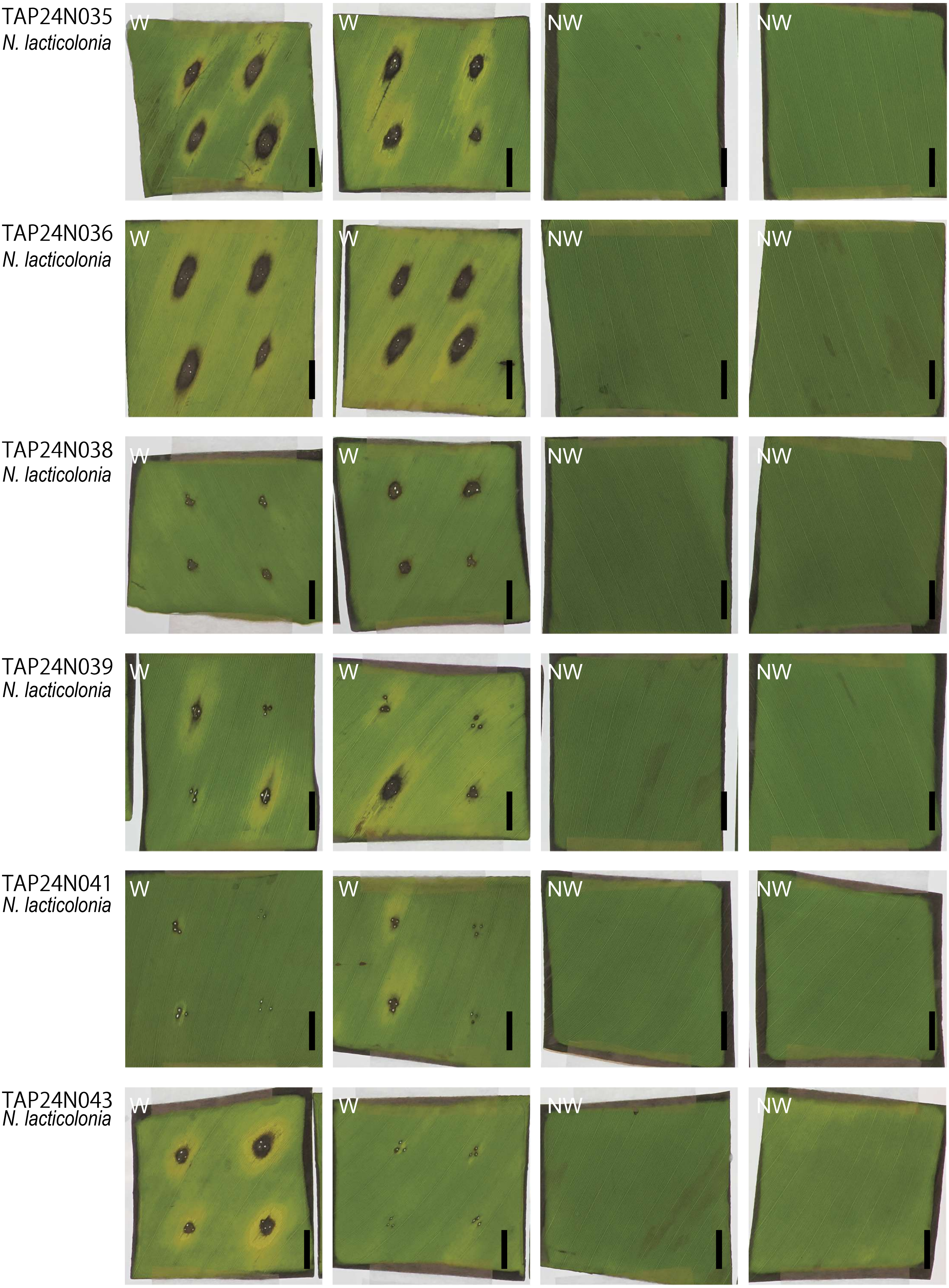


**Supplementary Fig 5** Continued.

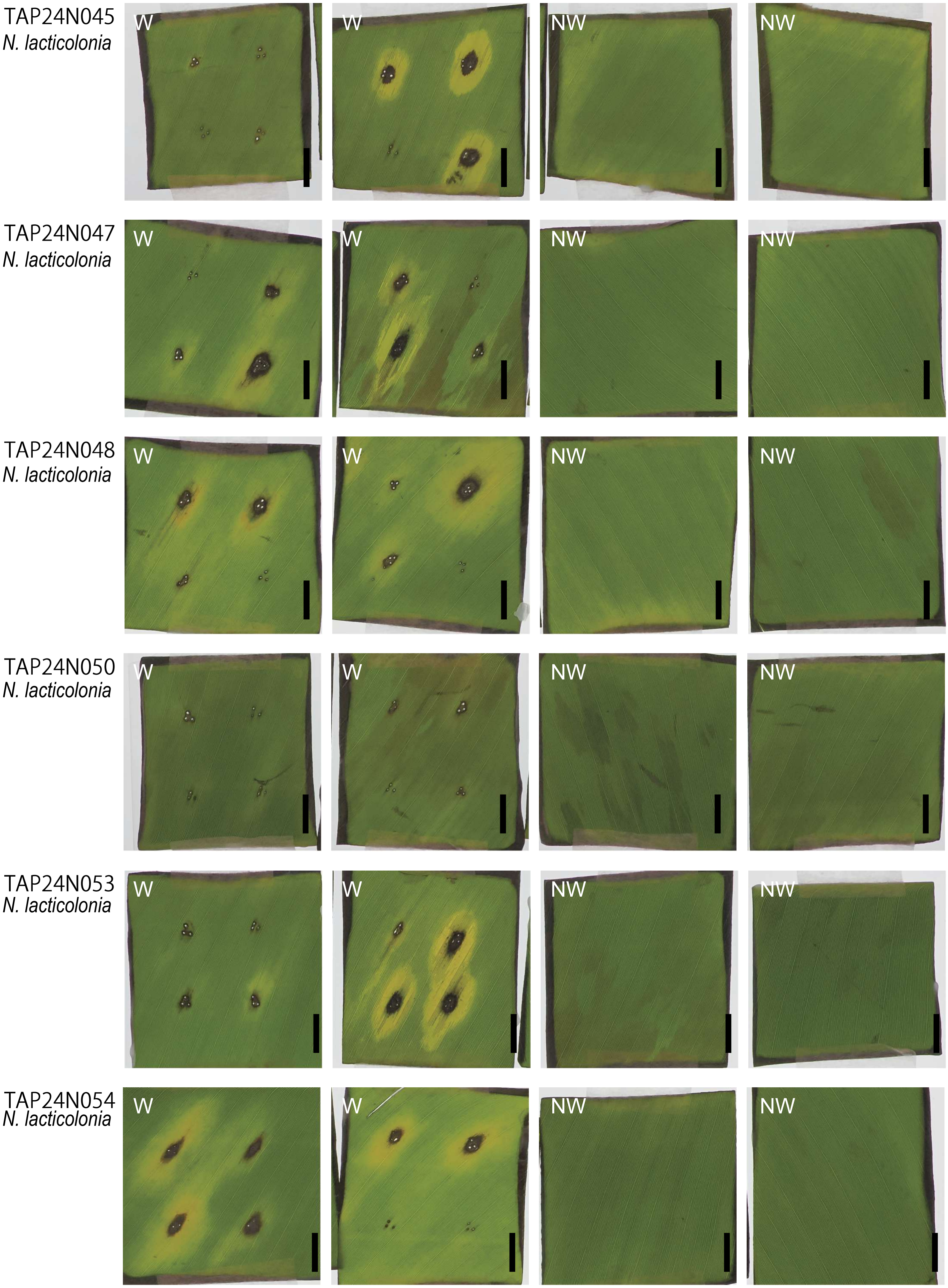


**Supplementary Fig 5** Continued.

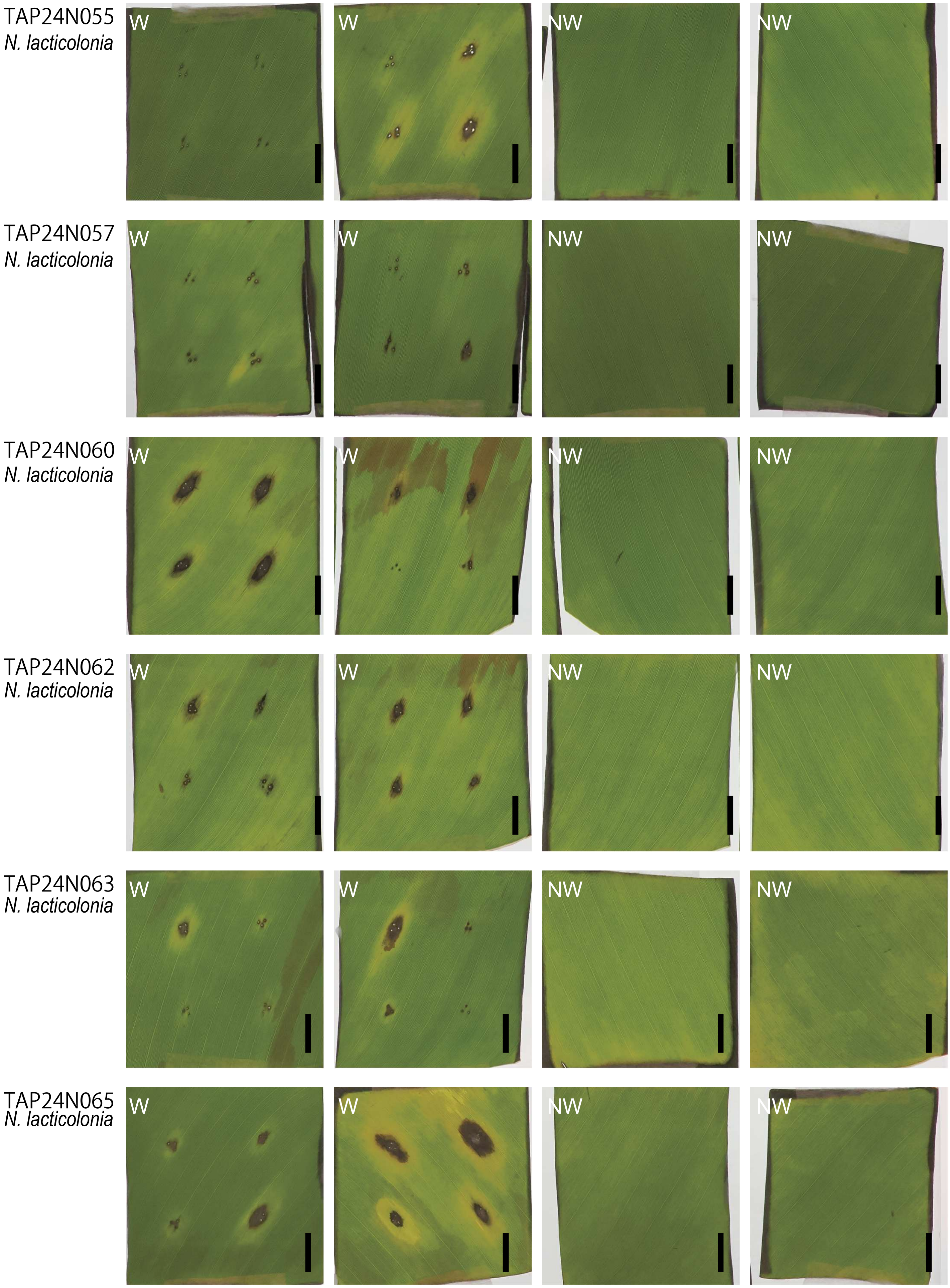


**Supplementary Fig 5** Continued.

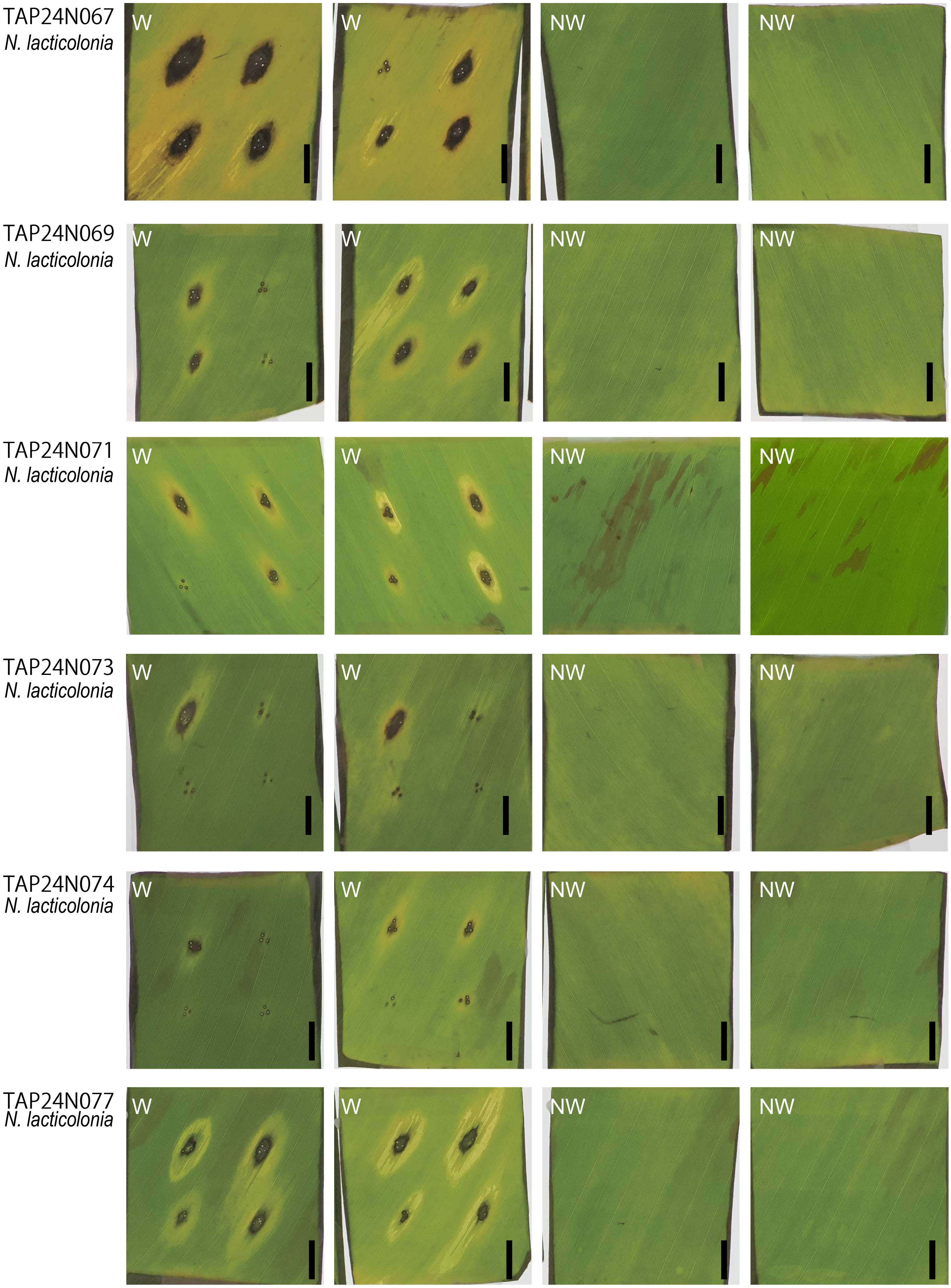


**Supplementary Fig 5** Continued.

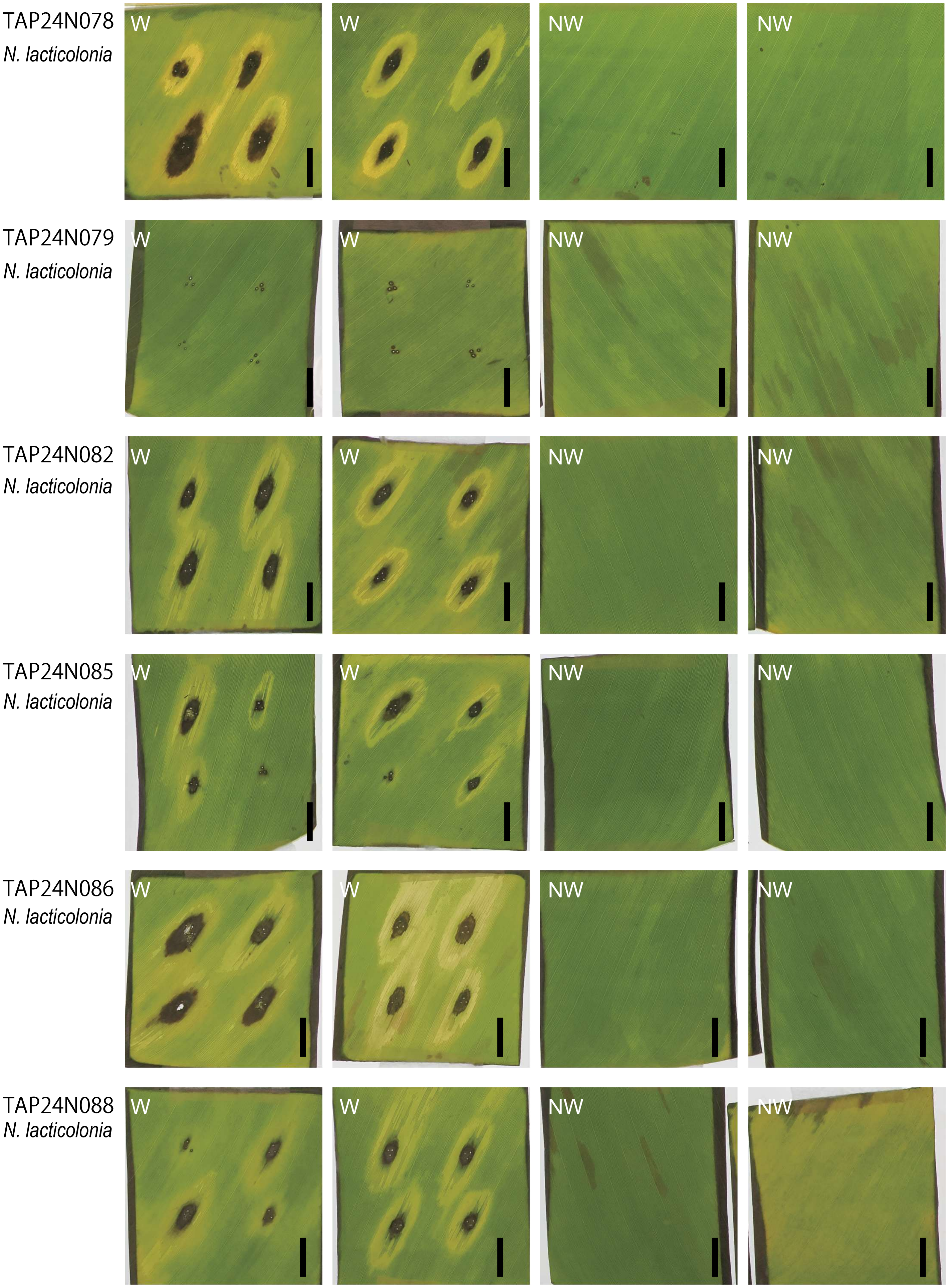


**Supplementary Fig 5** Continued.

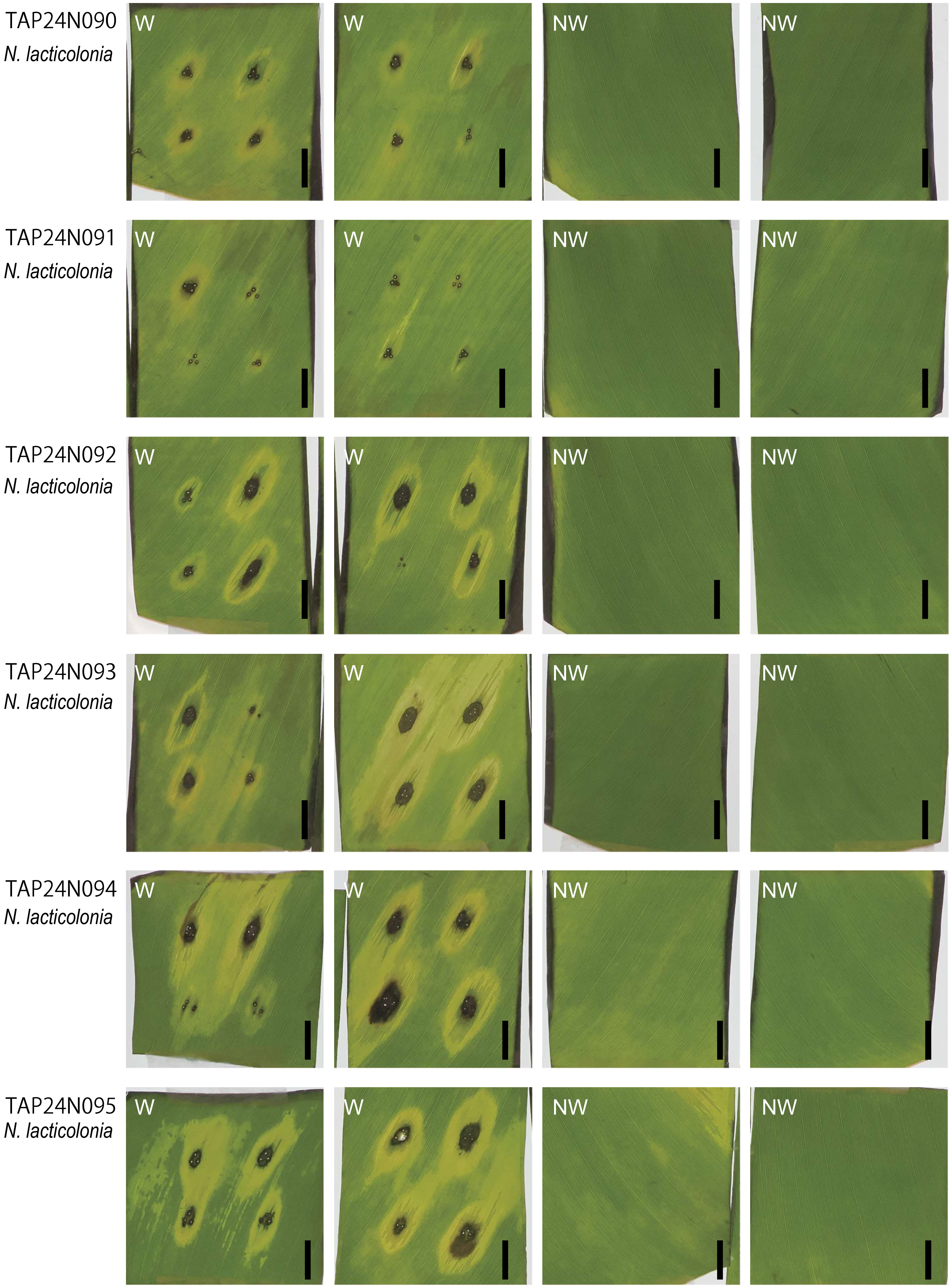


**Supplementary Fig 5** Continued.

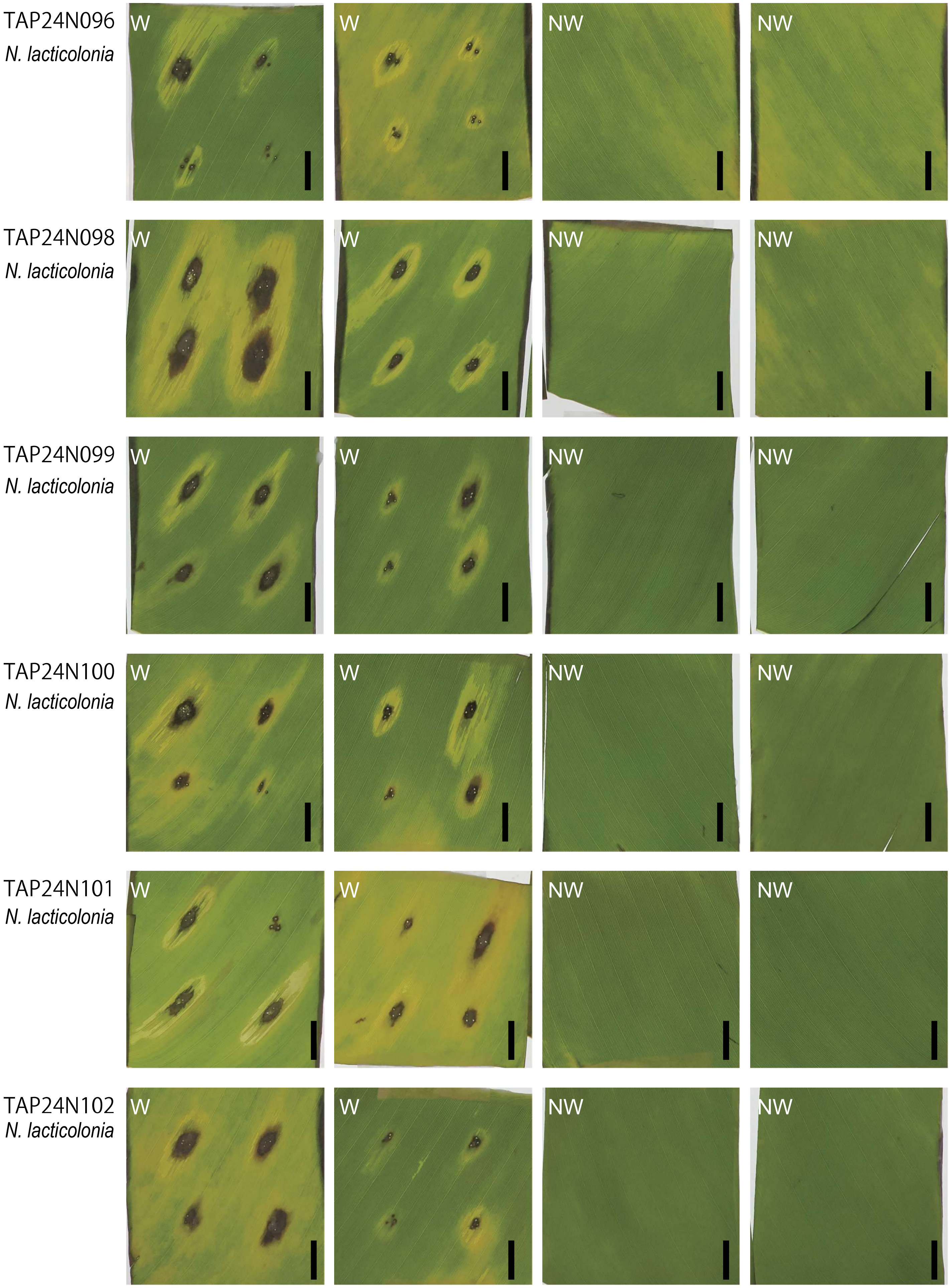


**Supplementary Fig 5** Continued.

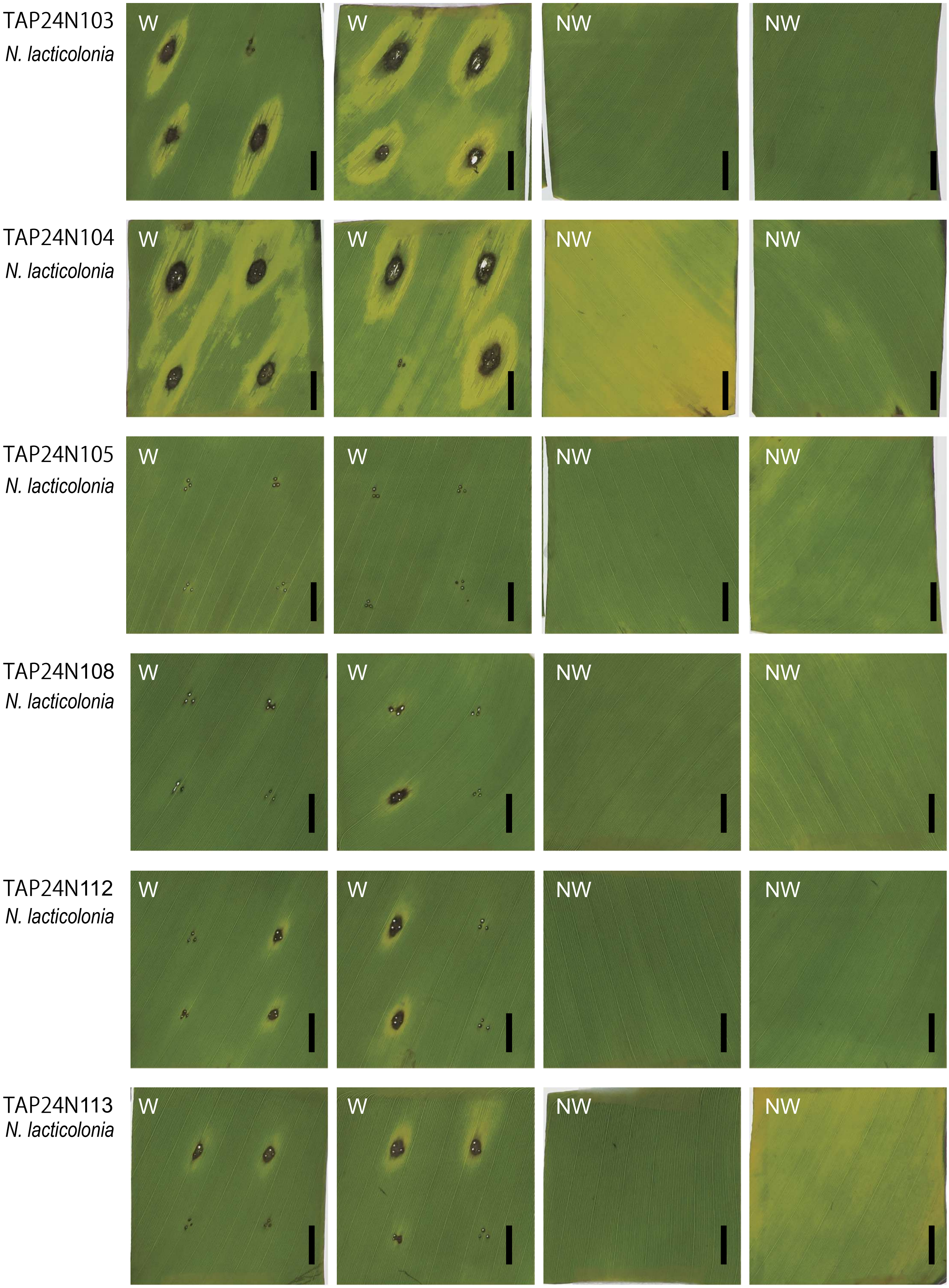


**Supplementary Fig 5** Continued.

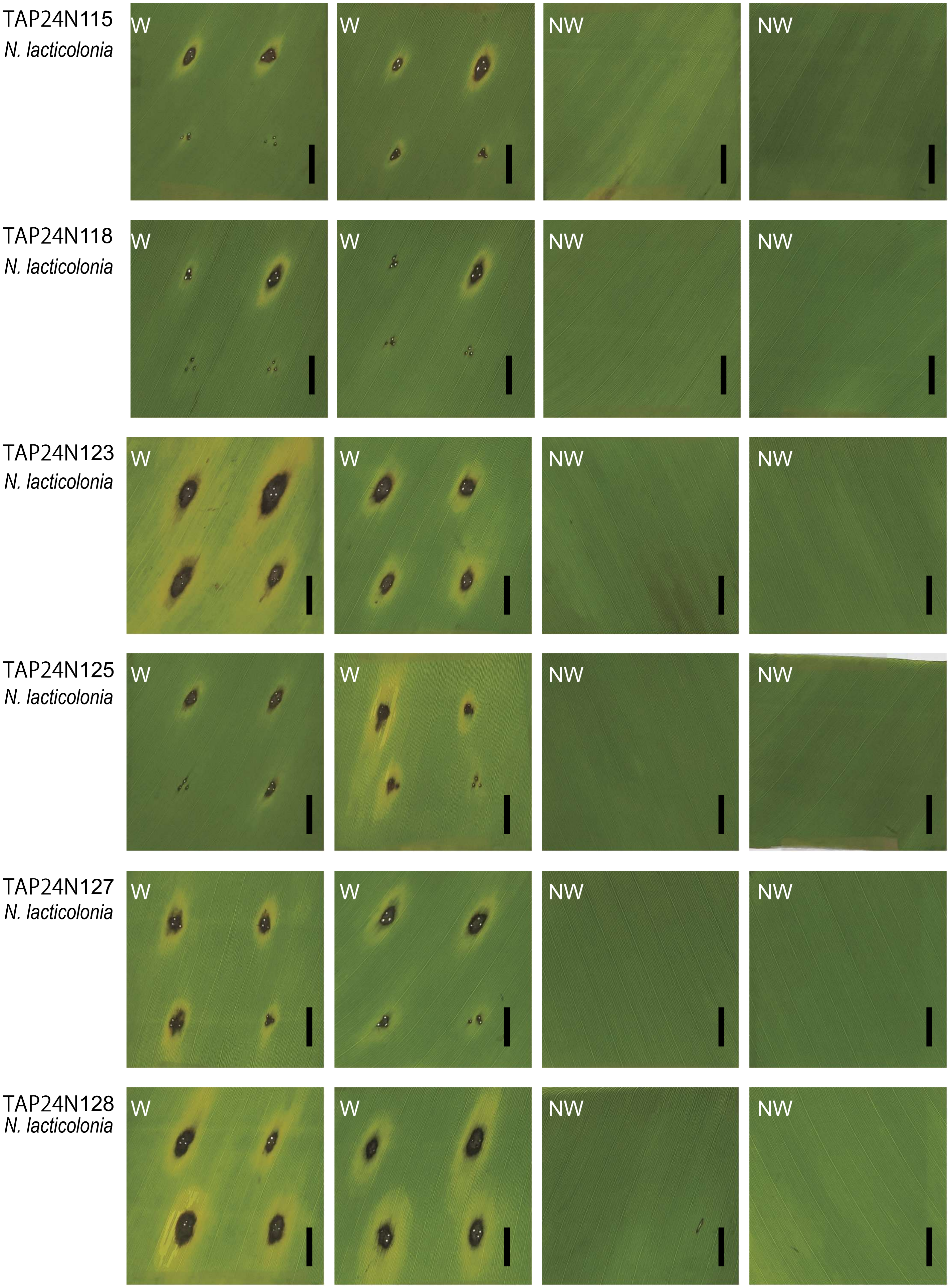


**Supplementary Fig 5** Continued.

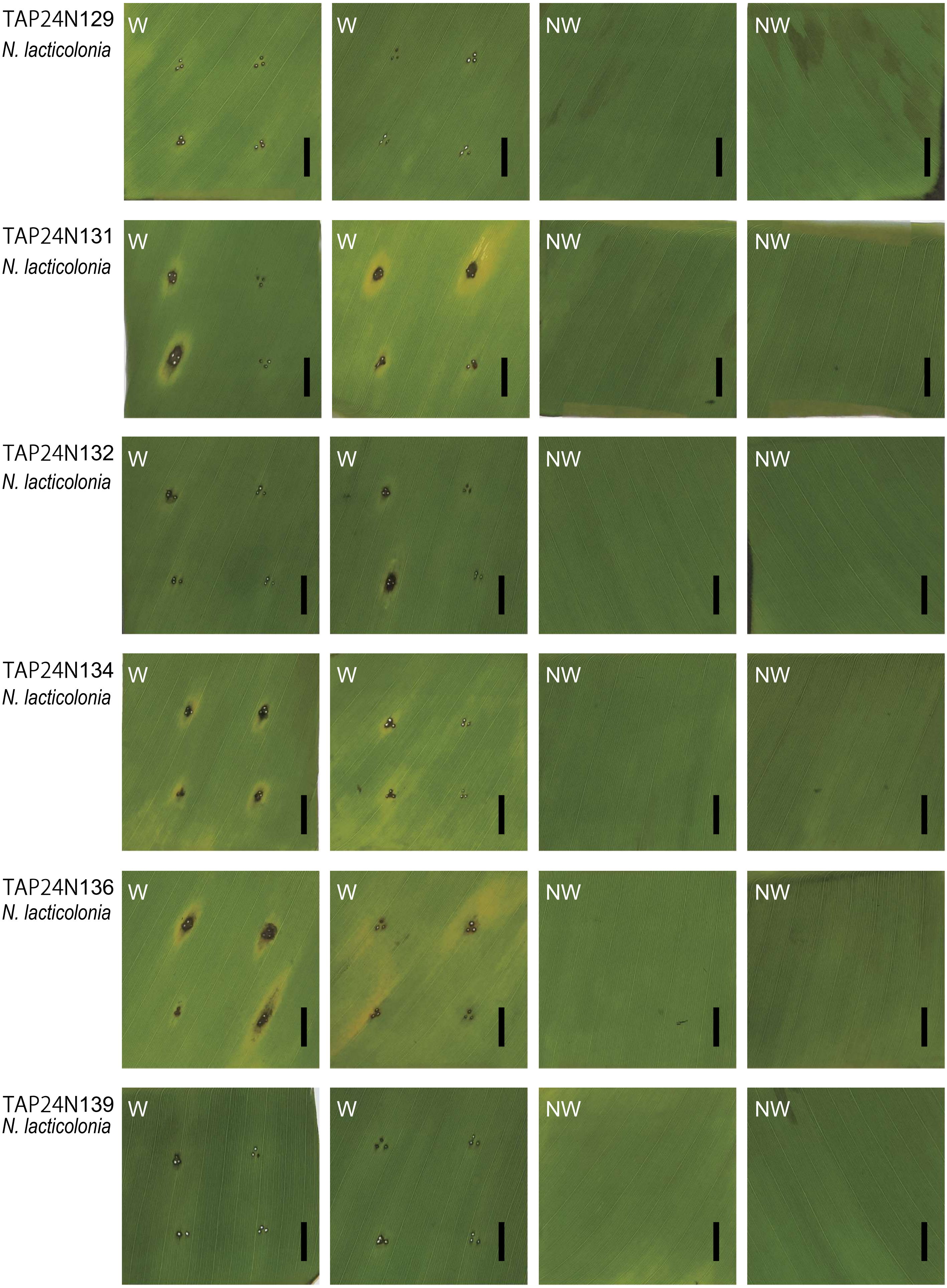


**Supplementary Fig 5** Continued.

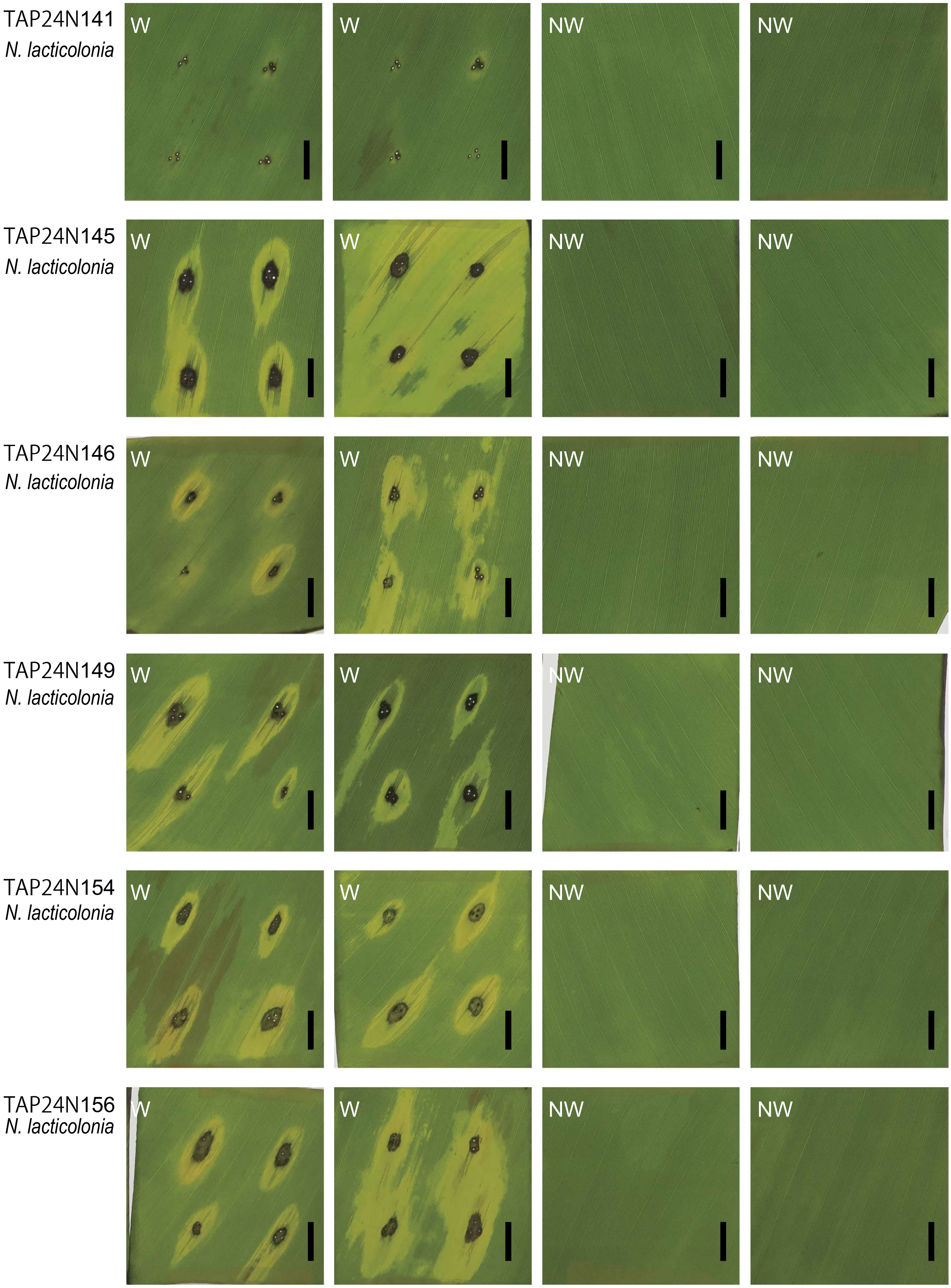


**Supplementary Fig 5** Continued.

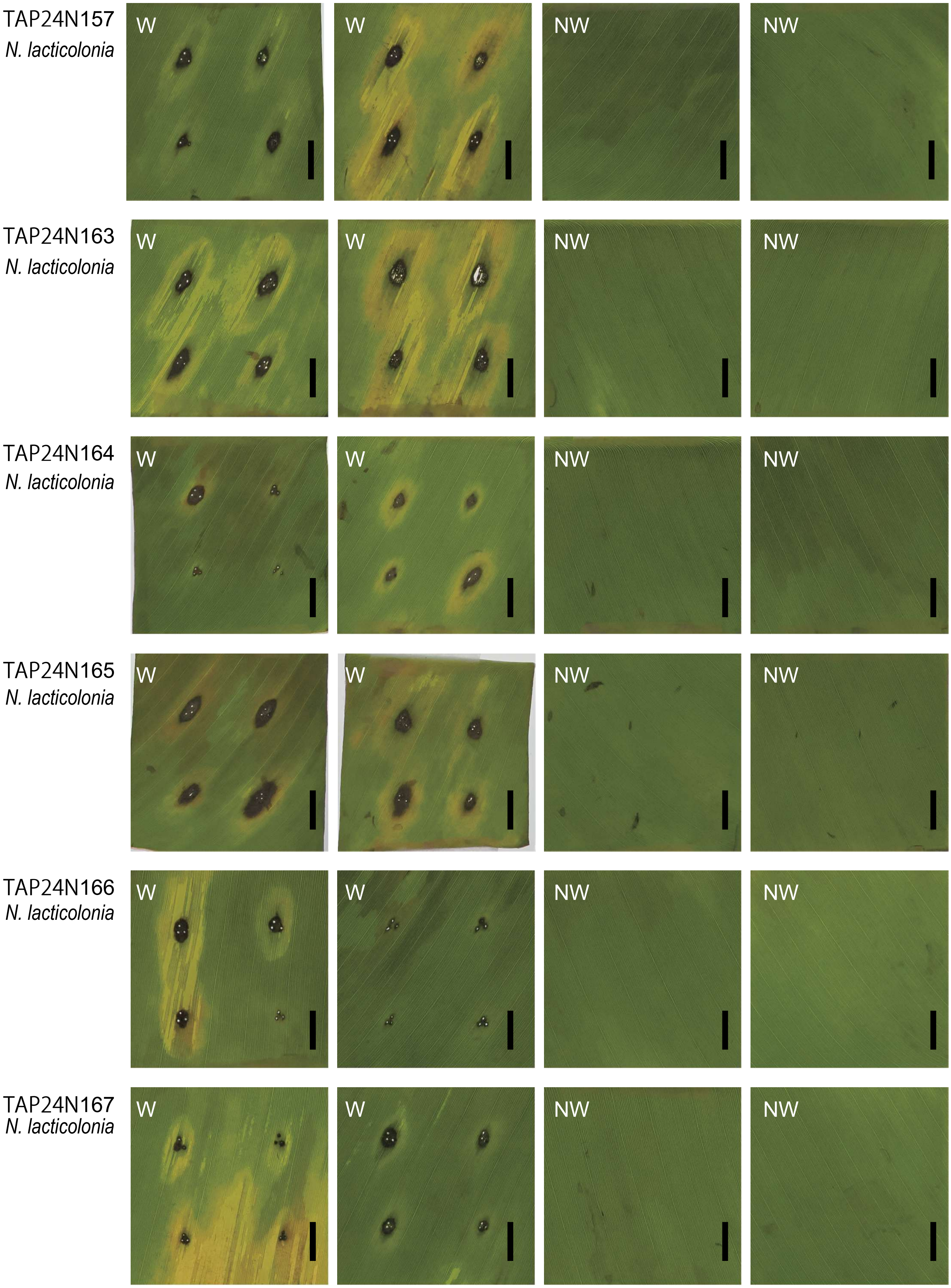


**Supplementary Fig 5** Continued.

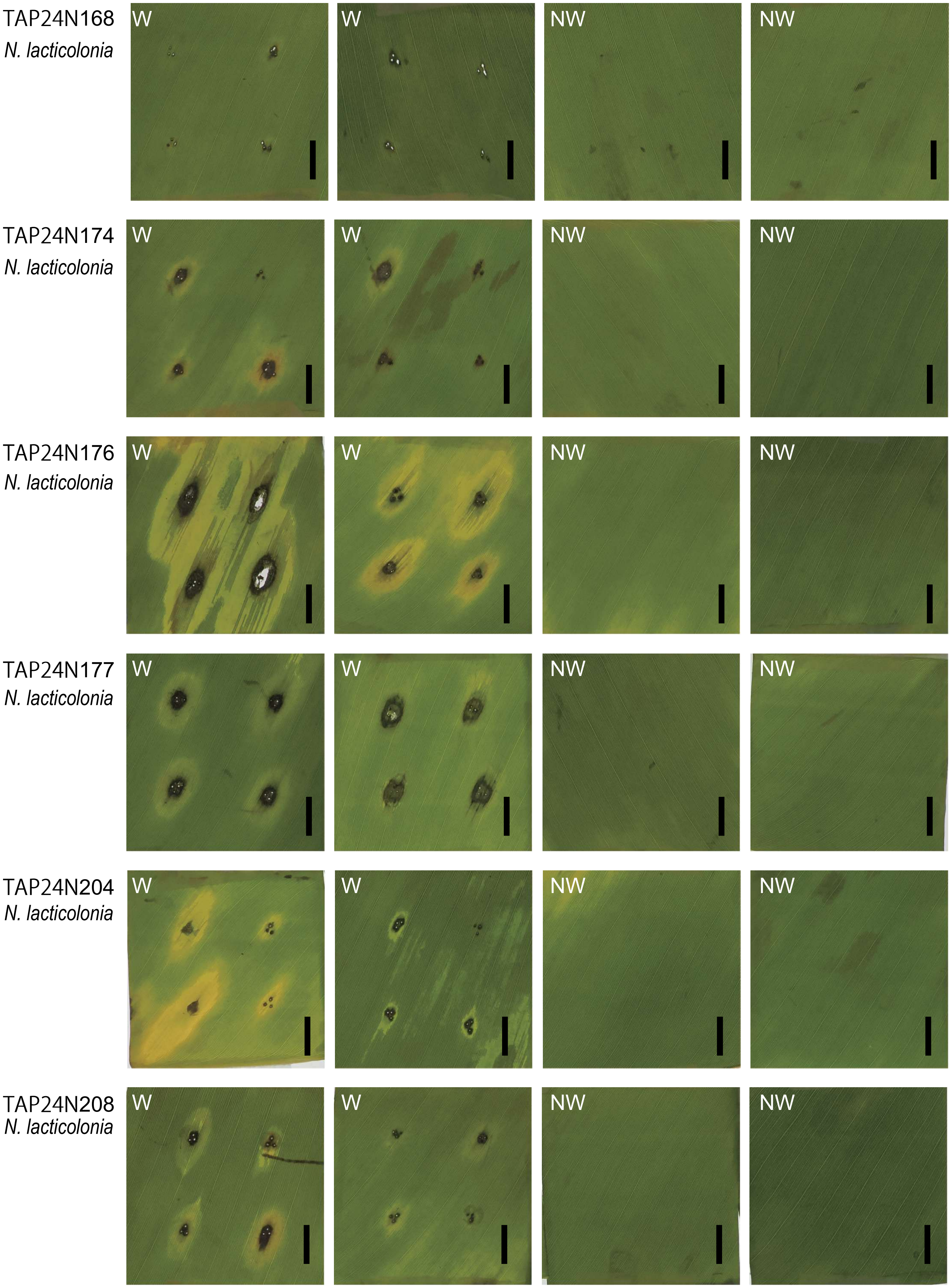


**Supplementary Fig 5** Continued.

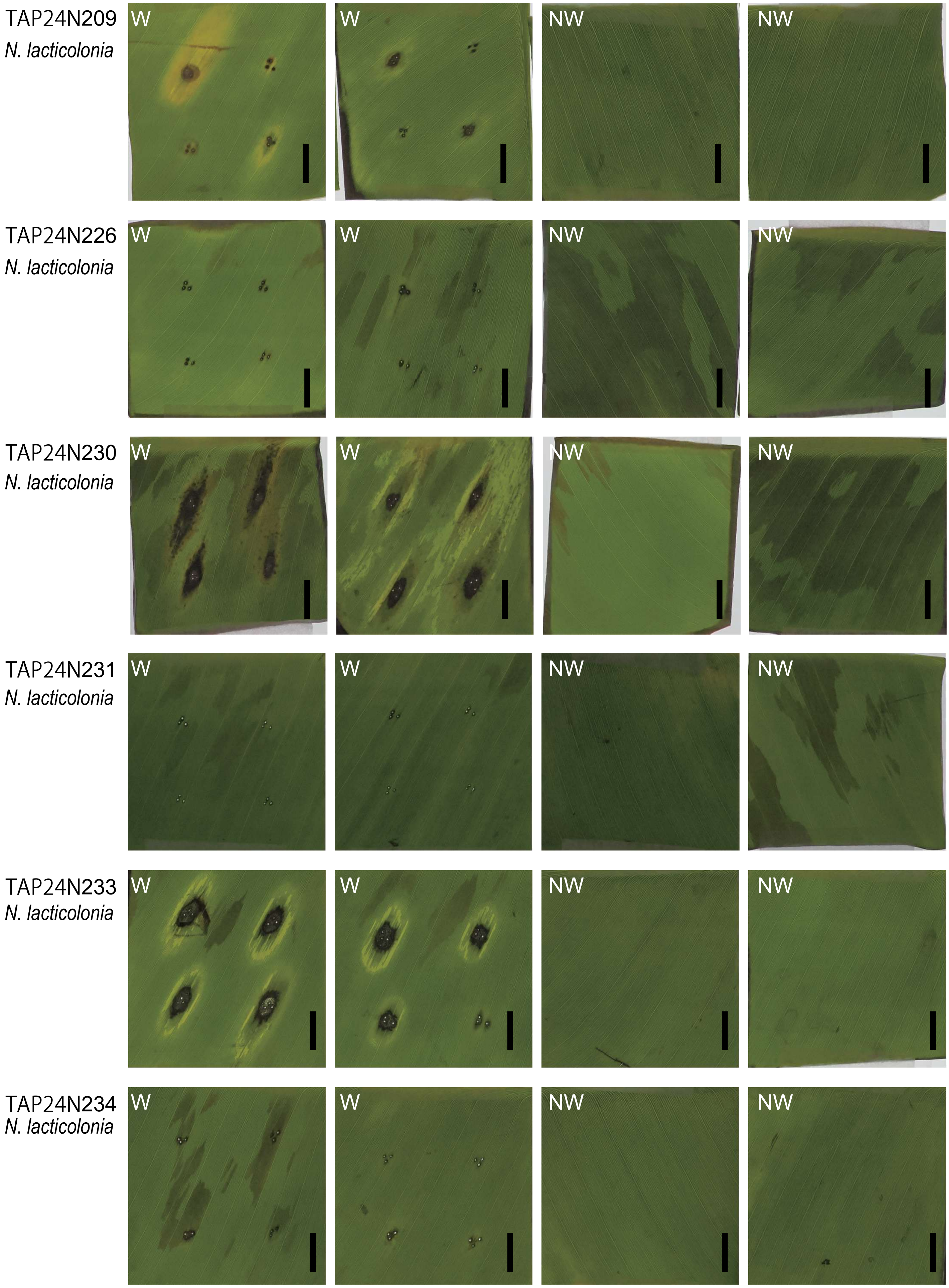


**Supplementary Fig 5** Continued.

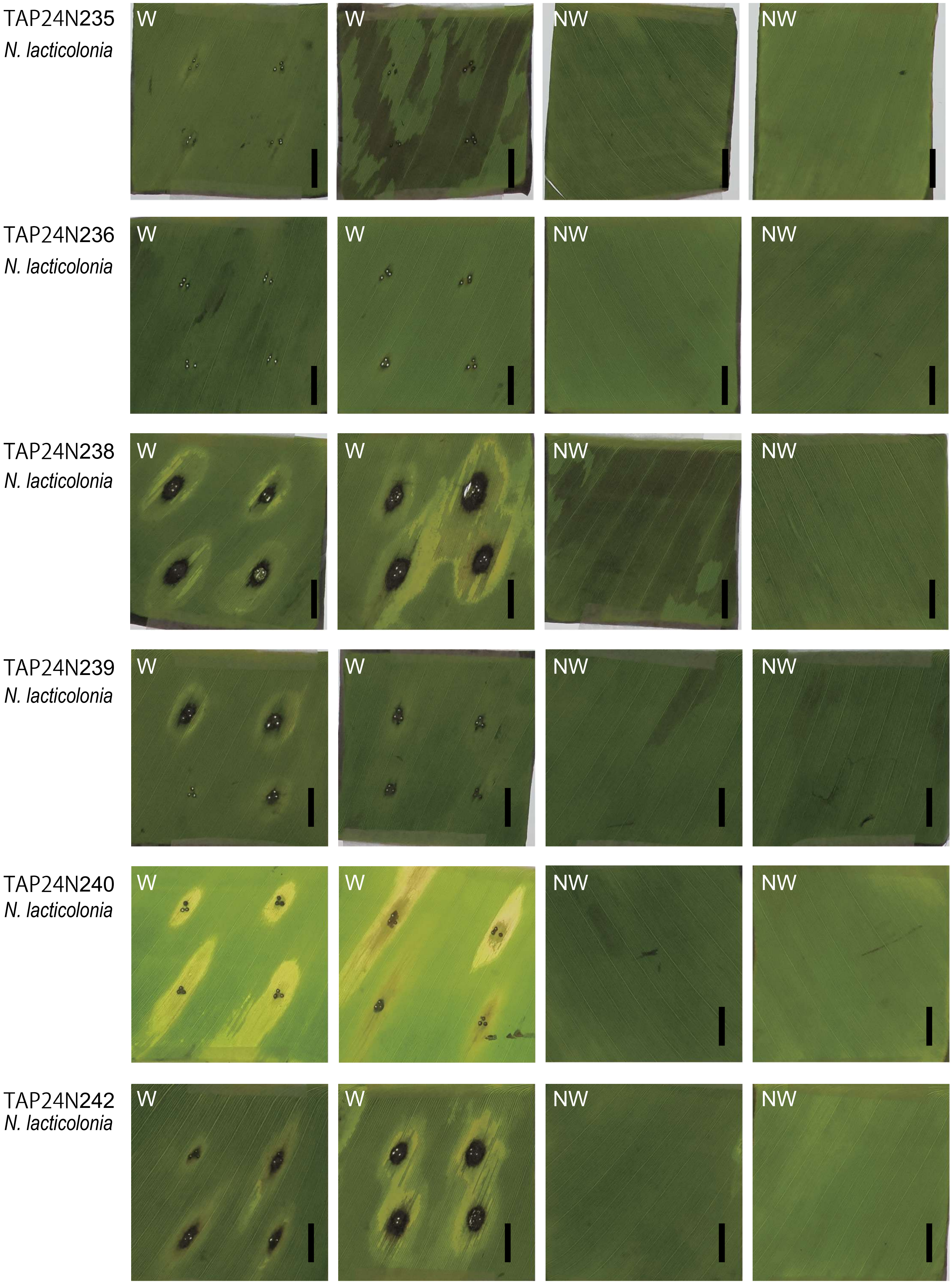


**Supplementary Fig 5** Continued.

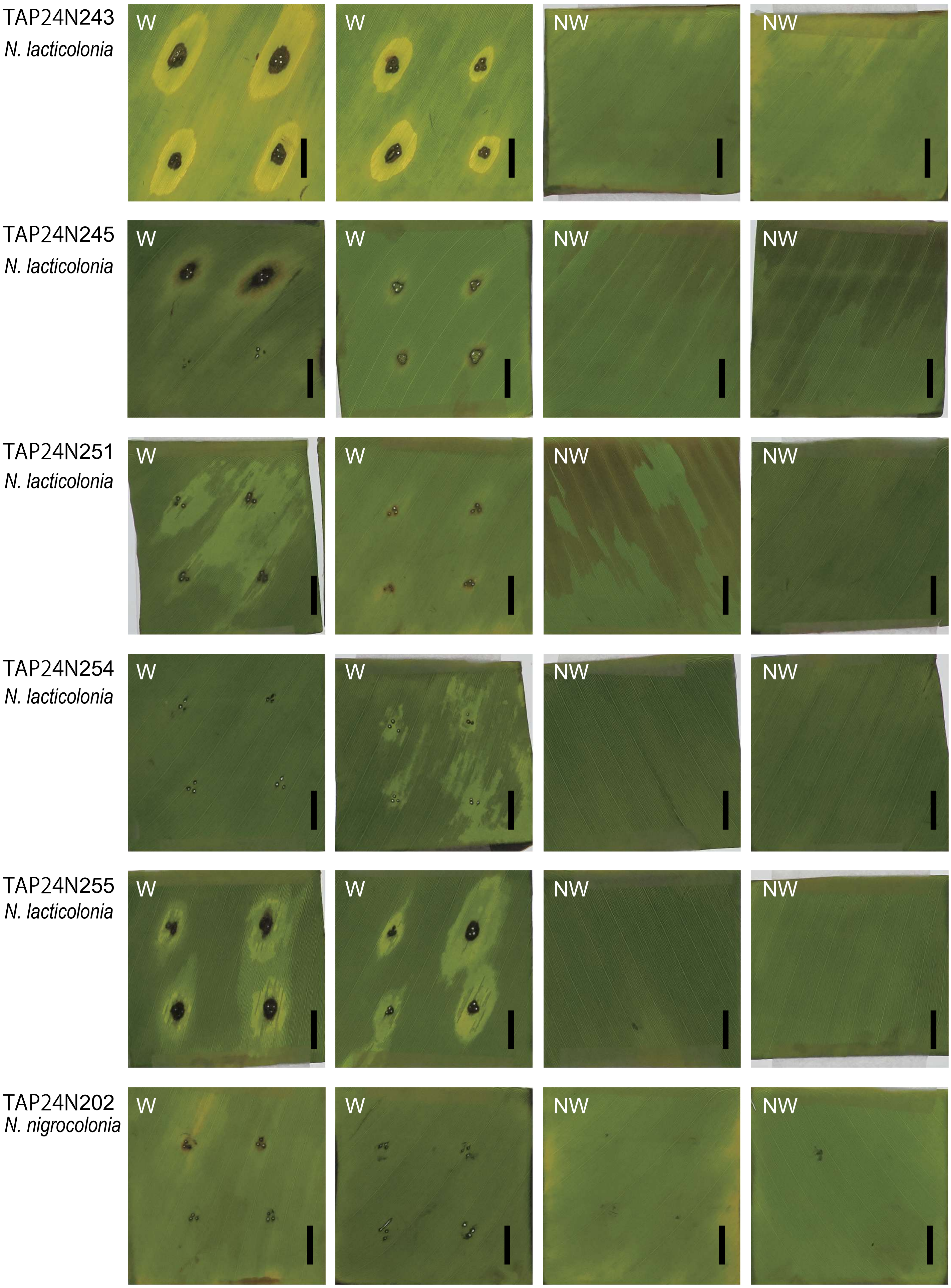


**Supplementary Fig 5** Continued.

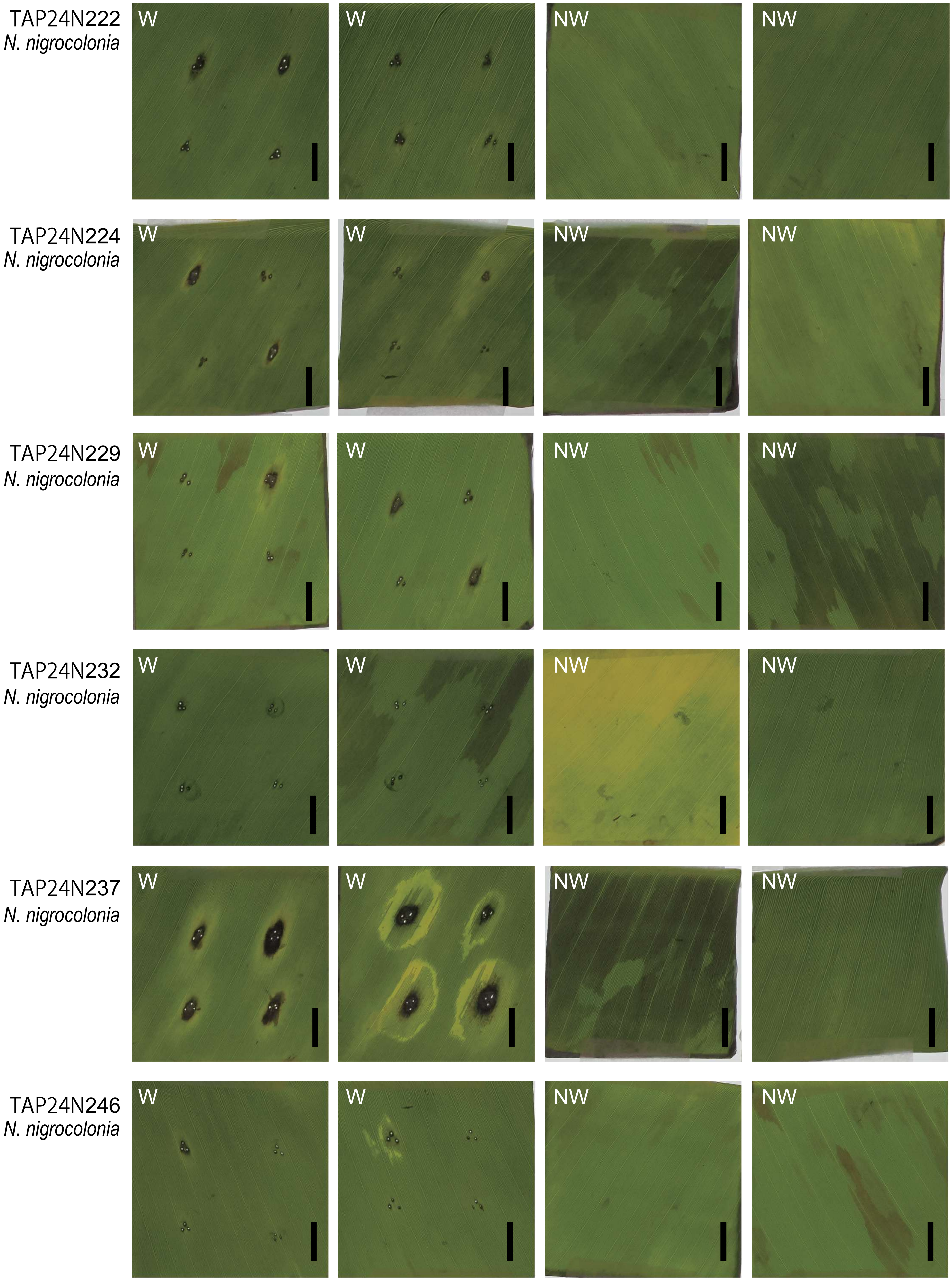


**Supplementary Fig 5** Continued.

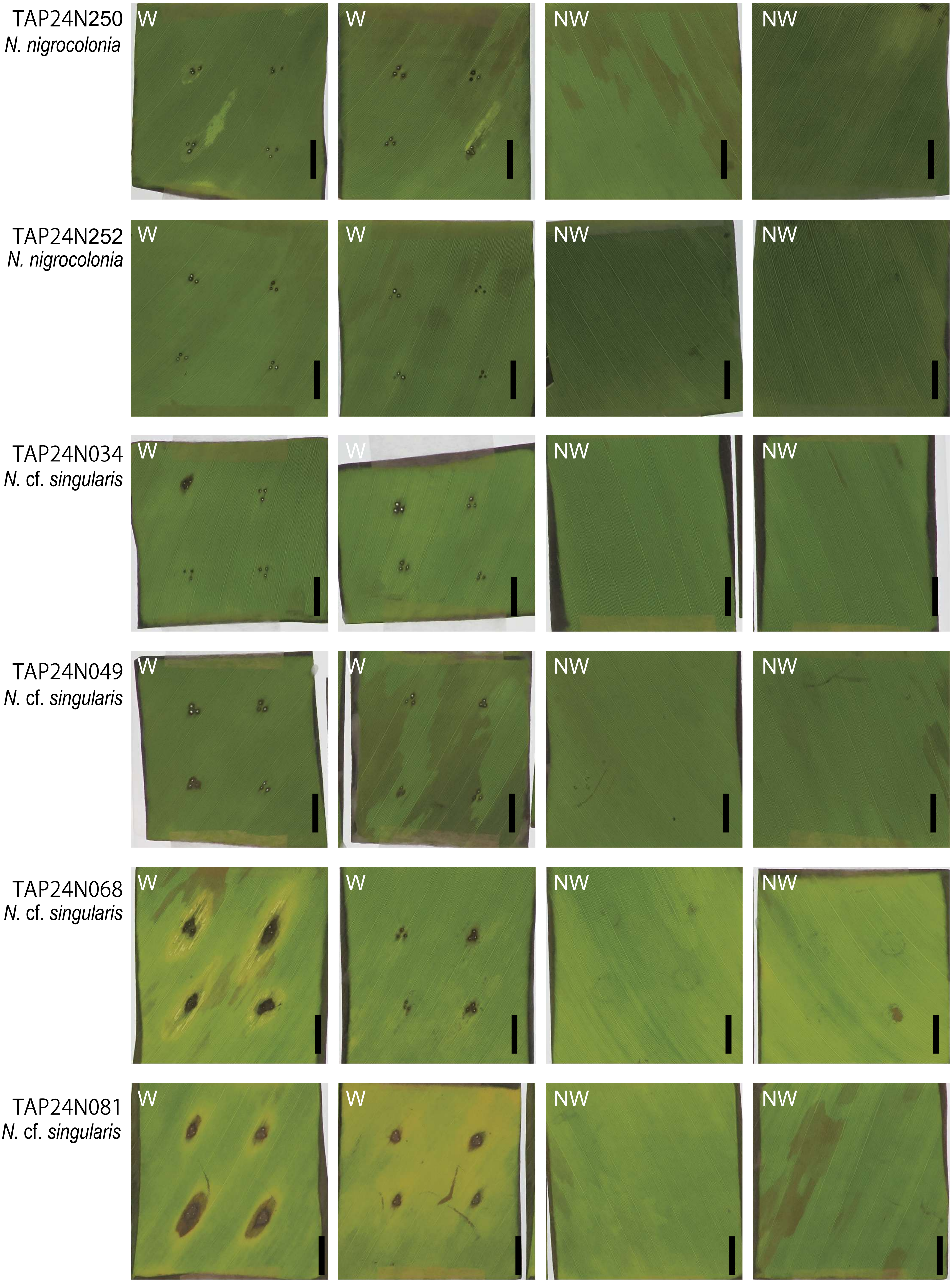


**Supplementary Fig 5** Continued.

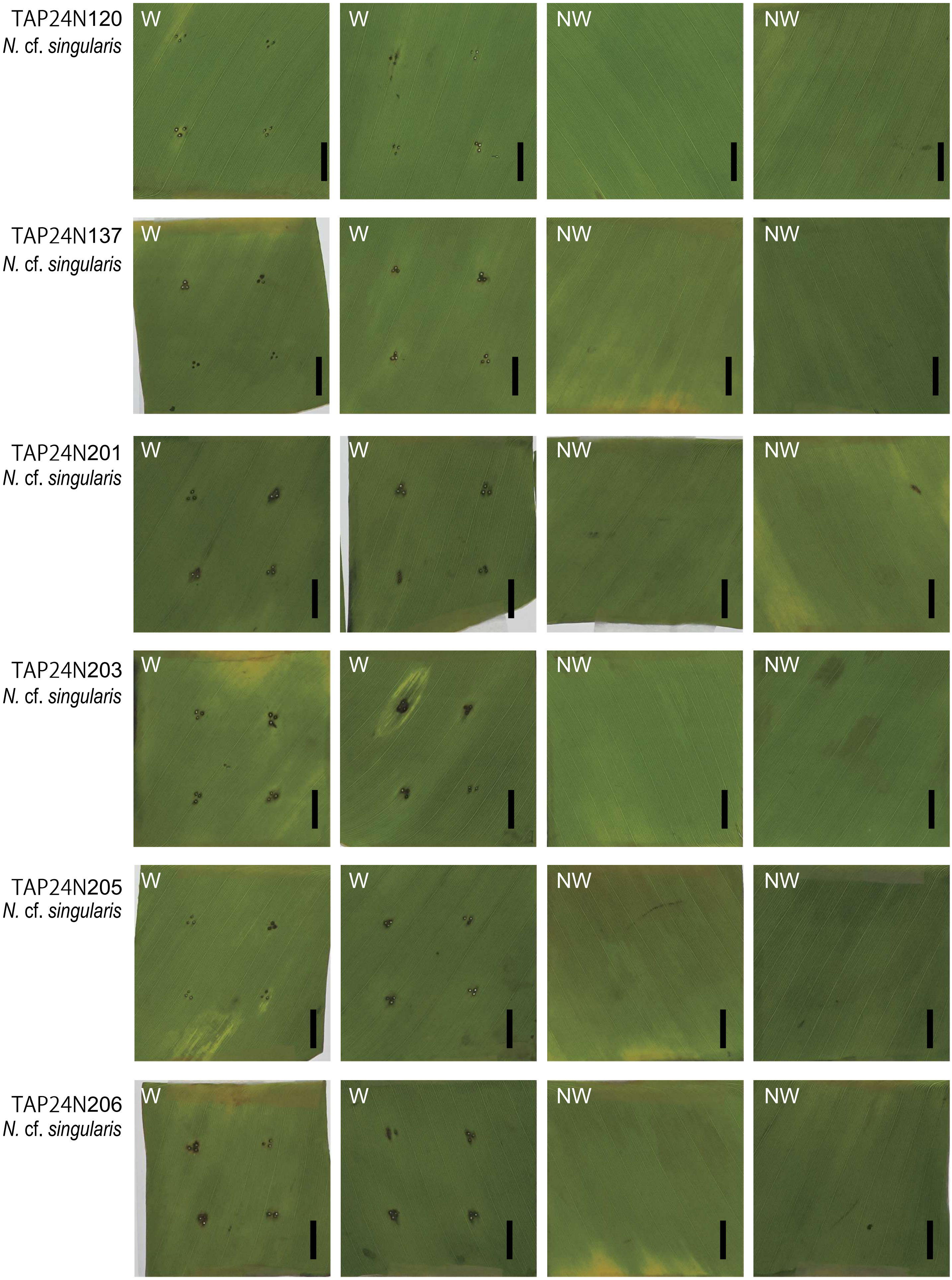


**Supplementary Fig 5** Continued.

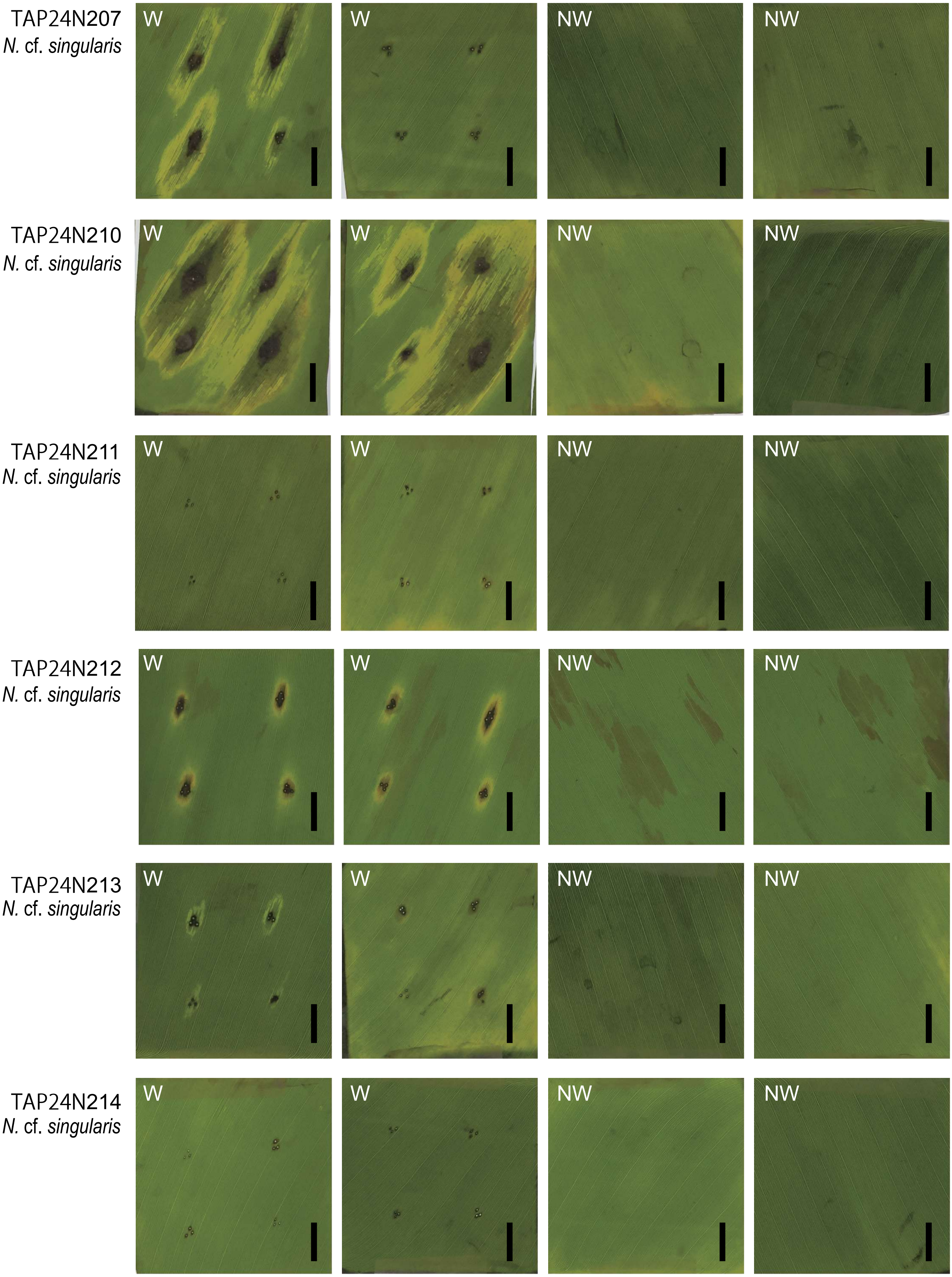


**Supplementary Fig 5** Continued.

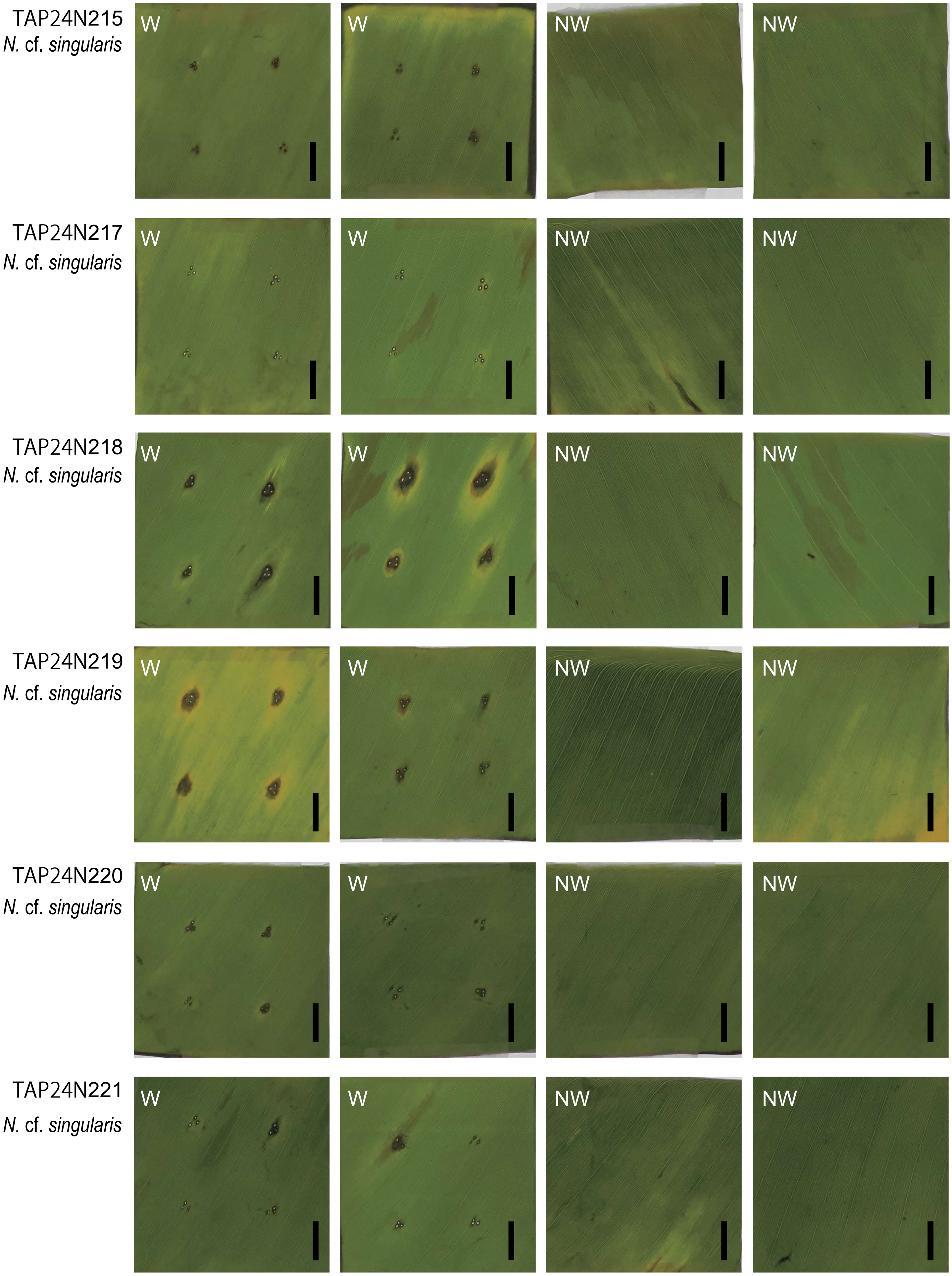


**Supplementary Fig 5** Continued.

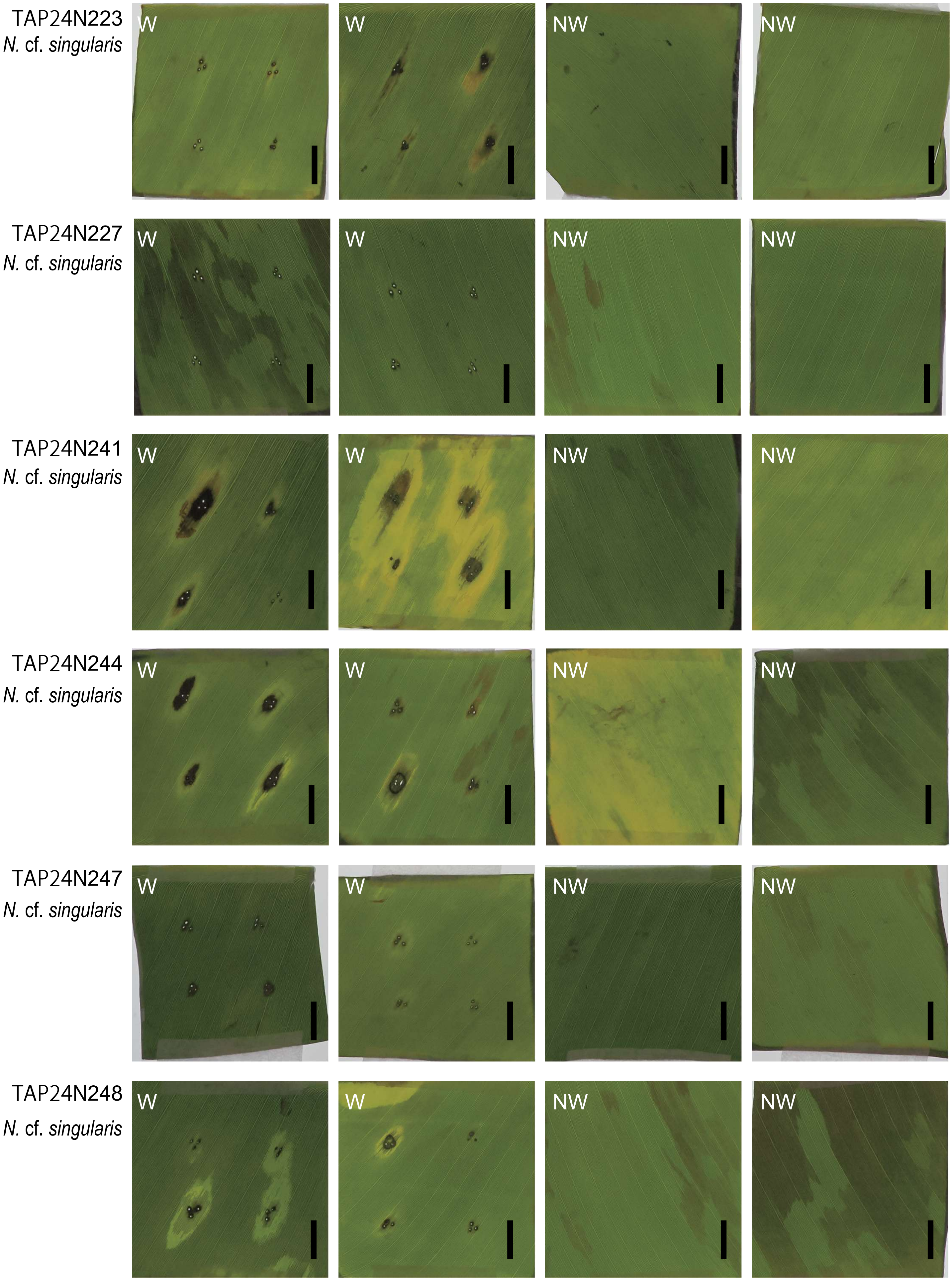


**Supplementary Fig 5** Continued.

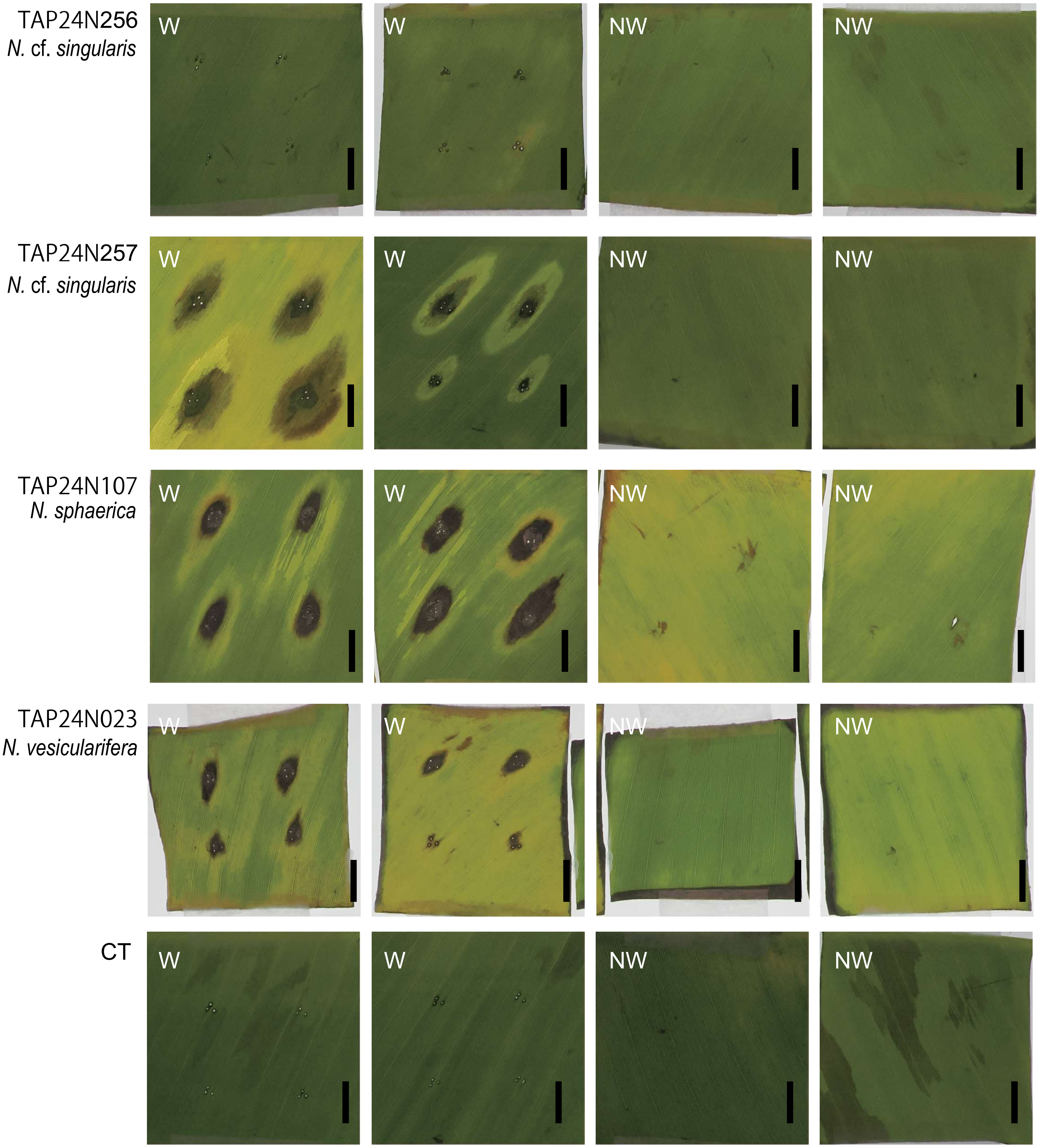


**Supplementary Fig 5** Continued.
