## Supplementary material for "Occurrence of *Nigrospora spp.* as the predominant causal agents of leaf spot disease in Cavendish banana in banana plantations in Mindanao Island, Philippines": Supplementary fig 6.pdf

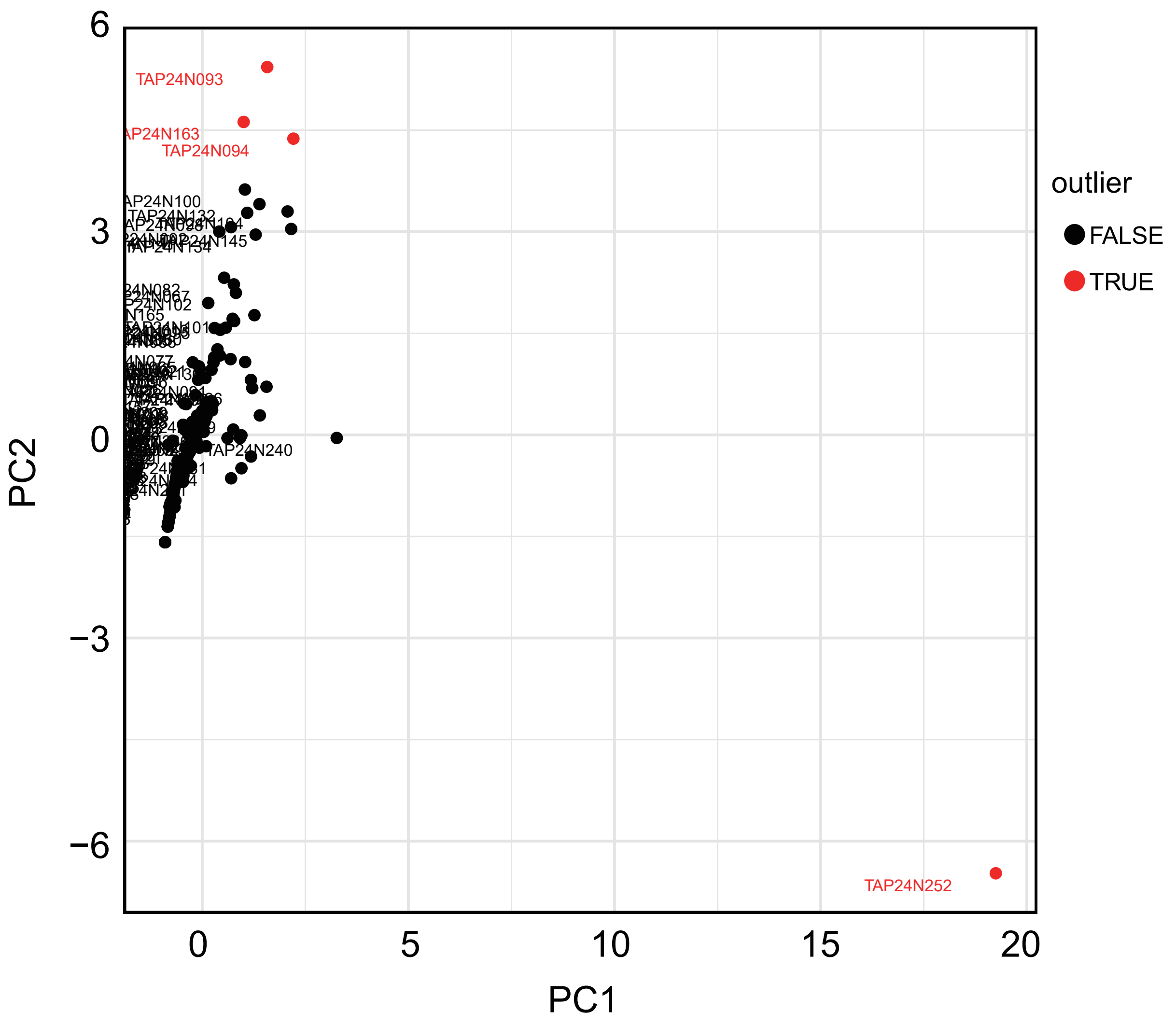


**Supplementary Fig. 6** Detection of outliers in the Principal Component Analysis based on the inhibition rates of isolates of 14 fungicides.
