## Supplementary material for "Occurrence of *Nigrospora spp.* as the predominant causal agents of leaf spot disease in Cavendish banana in banana plantations in Mindanao Island, Philippines": Supplementary fig 7.pdf

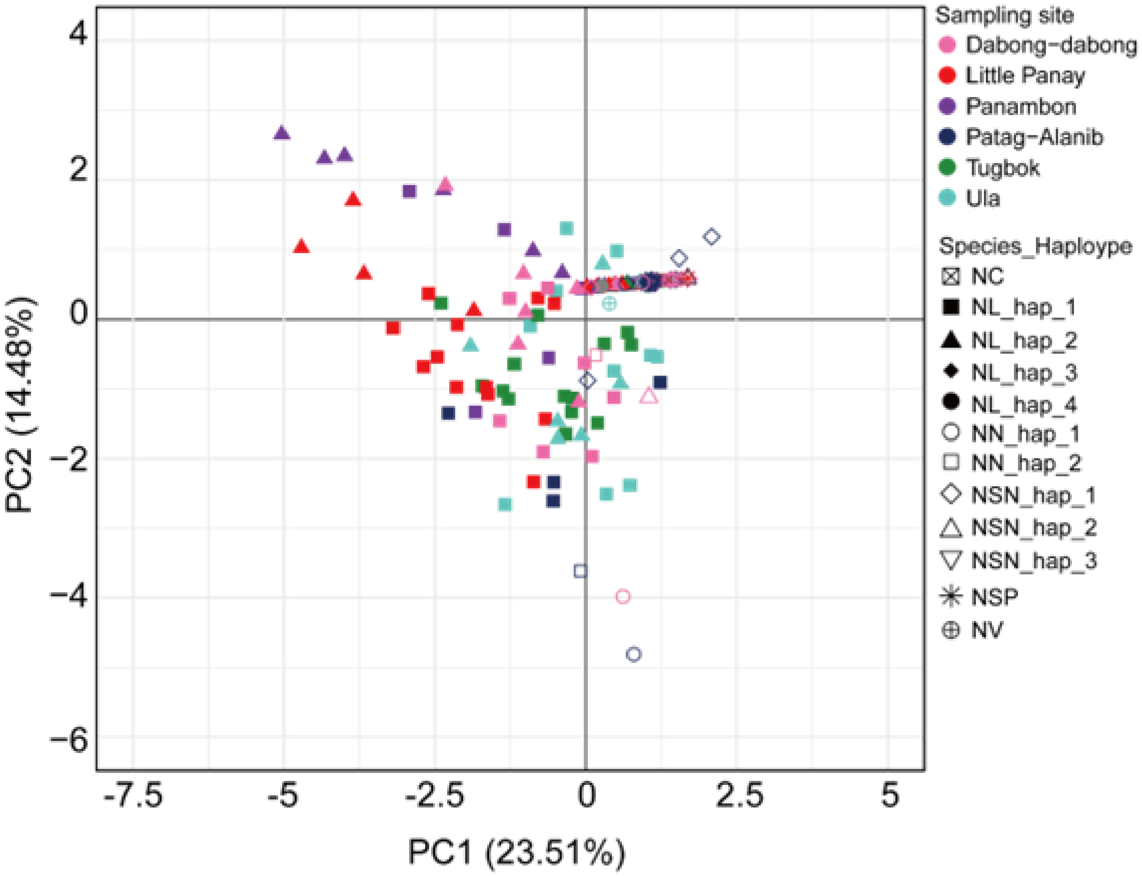


**Supplemental Fig. 7** Principal Component Analysis based on the inhibition rates of of 14 fungicides against fungal isolates.
