## Supplementary material for "Occurrence of *Nigrospora spp.* as the predominant causal agents of leaf spot disease in Cavendish banana in banana plantations in Mindanao Island, Philippines": Supplementary fig 8.pdf

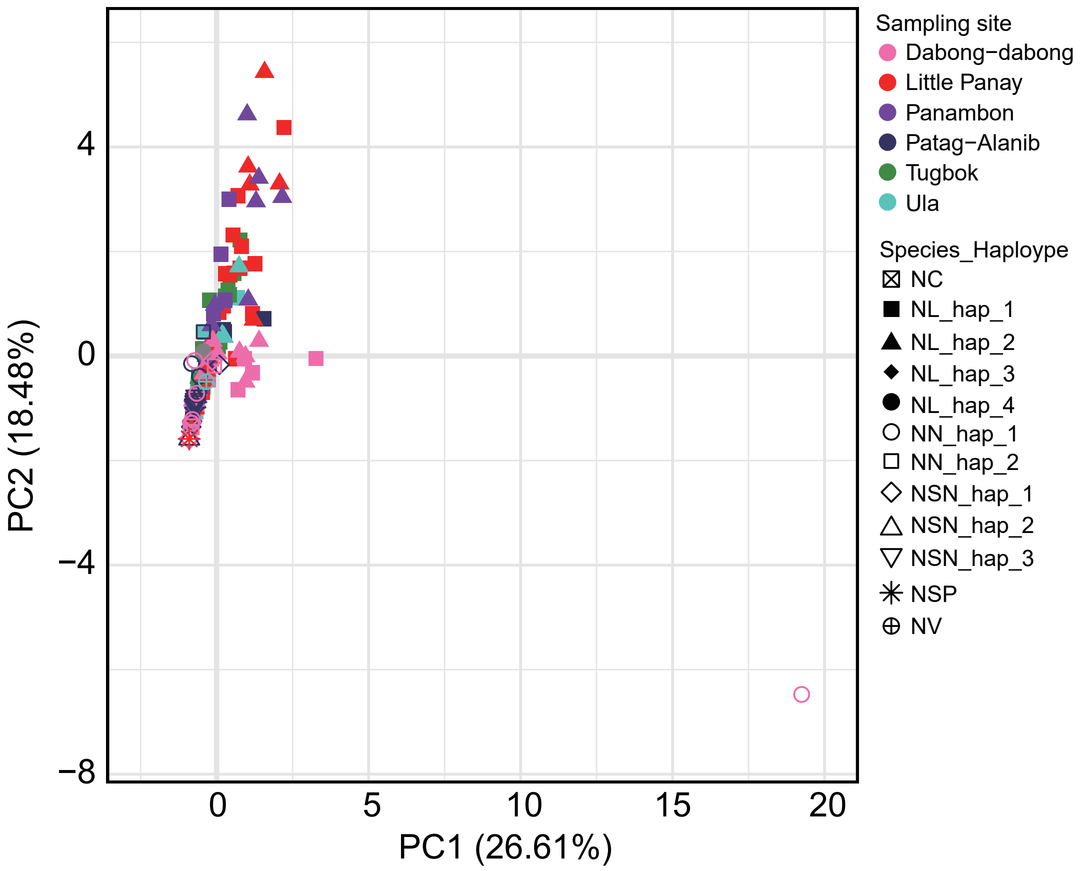


**Supplementary Fig. 8** Principal Component Analysis based on the inhibition rates of 14 fungicides against fungal isolates with outliers.
