## Supplementary material for "Occurrence of *Nigrospora spp.* as the predominant causal agents of leaf spot disease in Cavendish banana in banana plantations in Mindanao Island, Philippines": Supplementary fig 9.pdf

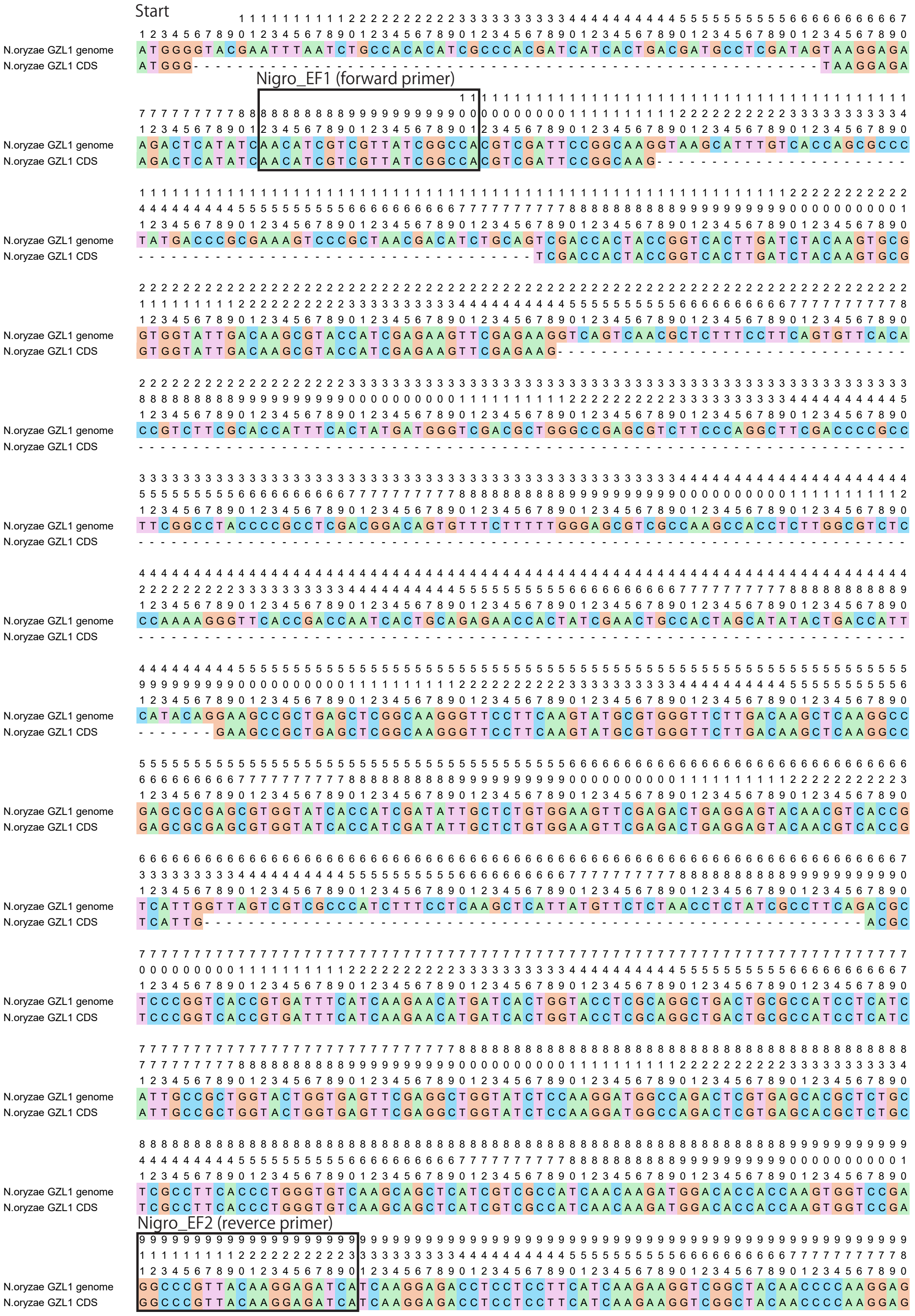


**Supplementary Fig 9** Positions of primers Nigro_EF1 (5'-AAC ATC GTC GTT ATC GGC CA-3') and Nigro_EF2 (5'-TGA TCT CCT TGT AAC GGG CC-3') designed for PCR amplification of the tef1a region. "*N. oryzae* GZL1 genome" contains exons and introns. "*N. oryzae* GZL1 CDS" sequence comprises only the coding region. Each site is numbered from the start codon at the top. Each sequence originates from the genome data of *N. oryzae* strain GZL1 deposited in the NCBI database (accession number: GCA_029489945). The positions where primers were designed are enclosed in black rectangular boxes, with the primer name indicated above. Primer design was performed using the primer design tool primer3plus (https://www.primer3plus.com/) based on sequences obtained from the genome data. Default settings were used during design.

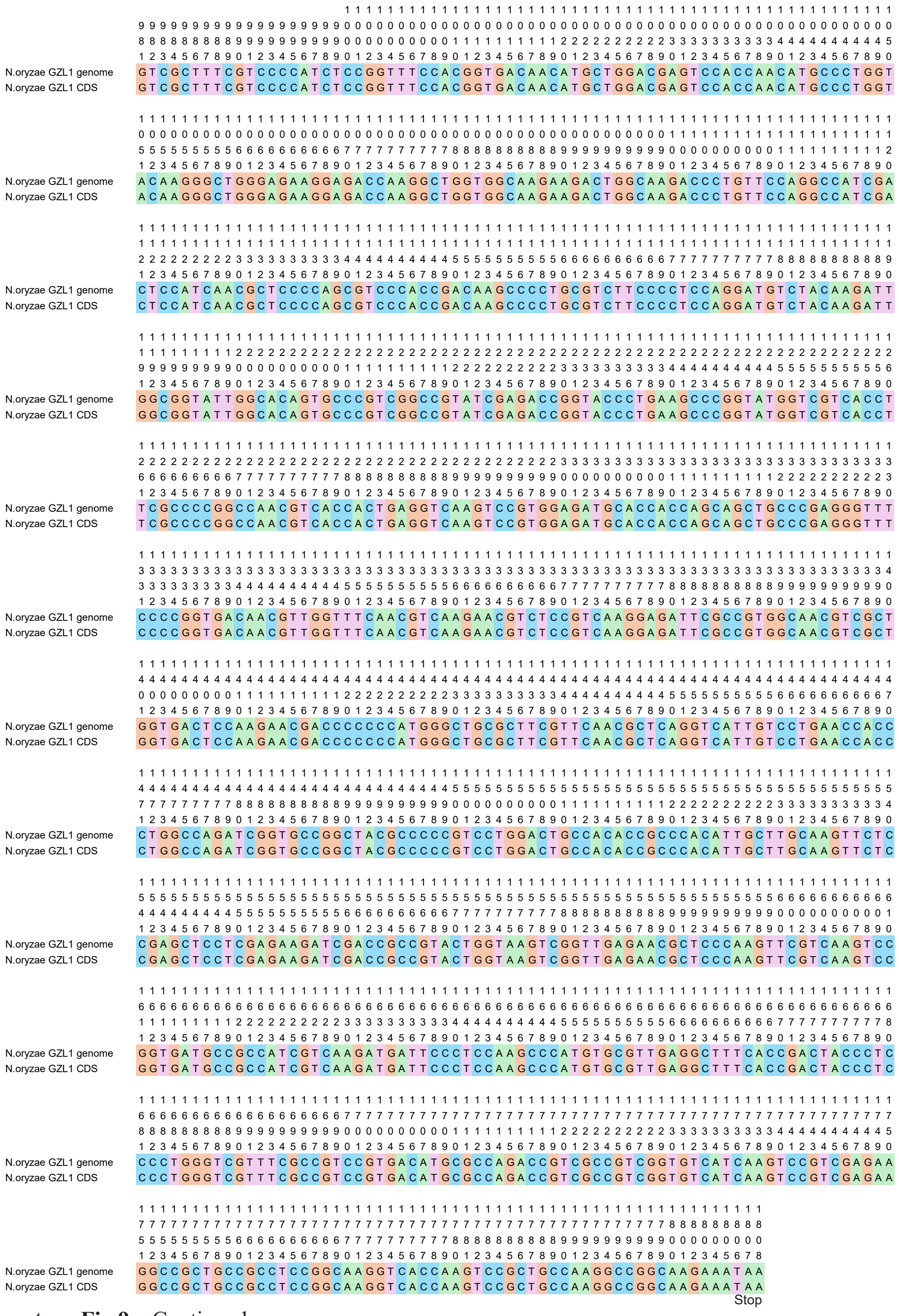


**Supplementary Fig 9** Continued
